# Global landscape of the host response to SARS-CoV-2 variants reveals viral evolutionary trajectories

**DOI:** 10.1101/2022.10.19.512927

**Authors:** Mehdi Bouhaddou, Ann-Kathrin Reuschl, Benjamin J. Polacco, Lucy G. Thorne, Manisha R. Ummadi, Chengjin Ye, Romel Rosales, Adrian Pelin, Jyoti Batra, Gwendolyn M. Jang, Jiewei Xu, Jack M. Moen, Alicia Richards, Yuan Zhou, Bhavya Harjai, Erica Stevenson, Ajda Rojc, Roberta Ragazzini, Matthew V.X. Whelan, Wilhelm Furnon, Giuditta De Lorenzo, Vanessa Cowton, Abdullah M. Syed, Alison Ciling, Noa Deutsch, Daniel Pirak, Giulia Dowgier, Dejan Mesner, Jane L. Turner, Briana L. McGovern, M. Luis Rodriguez, Rocio Leiva-Rebollo, Alistair S. Dunham, Xiaofang Zhong, Manon Eckhardt, Andrea Fossati, Nicholas Liotta, Thomas Kehrer, Anastasija Cupic, Magda Rutkowska, Nacho Mena, Sadaf Aslam, Alyssa Hoffert, Helene Foussard, John Pham, Molly Lyons, Laura Donahue, Aliesha Griffin, Rebecca Nugent, Kevin Holden, Robert Deans, Pablo Aviles, José Antonio López-Martín, Jose M. Jimeno, Kirsten Obernier, Jacqueline M. Fabius, Margaret Soucheray, Ruth Hüttenhain, Irwin Jungreis, Manolis Kellis, Ignacia Echeverria, Kliment Verba, Paola Bonfanti, Pedro Beltrao, Roded Sharan, Jennifer A. Doudna, Luis Martinez-Sobrido, Arvind Patel, Massimo Palmarini, Lisa Miorin, Kris White, Danielle L. Swaney, Adolfo García-Sastre, Clare Jolly, Lorena Zuliani-Alvarez, Greg J. Towers, Nevan J. Krogan

## Abstract

A series of SARS-CoV-2 variants of concern (VOCs) have evolved in humans during the COVID-19 pandemic—Alpha, Beta, Gamma, Delta, and Omicron. Here, we used global proteomic and genomic analyses during infection to understand the molecular responses driving VOC evolution. We discovered VOC-specific differences in viral RNA and protein expression levels, including for N, Orf6, and Orf9b, and pinpointed several viral mutations responsible. An analysis of the host response to VOC infection and comprehensive interrogation of altered virus-host protein-protein interactions revealed conserved and divergent regulation of biological pathways. For example, regulation of host translation was highly conserved, consistent with suppression of VOC replication in mice using the translation inhibitor plitidepsin. Conversely, modulation of the host inflammatory response was most divergent, where we found Alpha and Beta, but not Omicron BA.1, antagonized interferon stimulated genes (ISGs), a phenotype that correlated with differing levels of Orf6. Additionally, Delta more strongly upregulated proinflammatory genes compared to other VOCs. Systematic comparison of Omicron subvariants revealed BA.5 to have evolved enhanced ISG and proinflammatory gene suppression that similarly correlated with Orf6 expression, effects not seen in BA.4 due to a mutation that disrupts the Orf6-nuclear pore interaction. Our findings describe how VOCs have evolved to fine-tune viral protein expression and protein-protein interactions to evade both innate and adaptive immune responses, offering a likely explanation for increased transmission in humans.

**One sentence summary:** Systematic proteomic and genomic analyses of SARS-CoV-2 variants of concern reveal how variant-specific mutations alter viral gene expression, virus-host protein complexes, and the host response to infection with applications to therapy and future pandemic preparedness.

## MAIN TEXT

The emergence and rapid global spread of the novel coronavirus SARS-CoV-2, the causative agent of the COVID-19 (Coronavirus disease-2019) pandemic, continues to impact public health and global economies, becoming one of the deadliest outbreaks in modern history (*1*). SARS-CoV-2 variants of concern (VOCs) began to emerge in September 2020, starting with lineage B.1.1.7 (Alpha) (*2*–*4*), followed by P.1 (Gamma) (*5*), B.1.351 (Beta) (*6*) and B.1.617.2 (Delta) (*7*). Phylogenetic studies indicate that VOCs have independently evolved from early-lineage wave 1 (W1) viruses, with epidemiologically dominant VOCs Alpha, Delta and Omicron BA.1 spreading globally while Beta and Gamma remained geographically restricted (Fig. 1A). Omicron encodes a highly mutated Spike protein and represents the most significant antigenic shift to date, effectively facilitating evasion of adaptive immunity acquired from vaccines and prior infections (*8*–*11*). Omicron BA.1 and BA.2 have been rapidly replaced by BA.4 and BA.5, suggesting Omicron is the first VOC to give rise to globally dominant subvariants (*9, 12, 13*). The emergence and evolution of VOCs is likely driven by selective pressures to adapt to a new host (i.e. humans) as well as escape innate and adaptive immune responses, ultimately facilitating increased human-to-human transmission.

**Figure 1.**
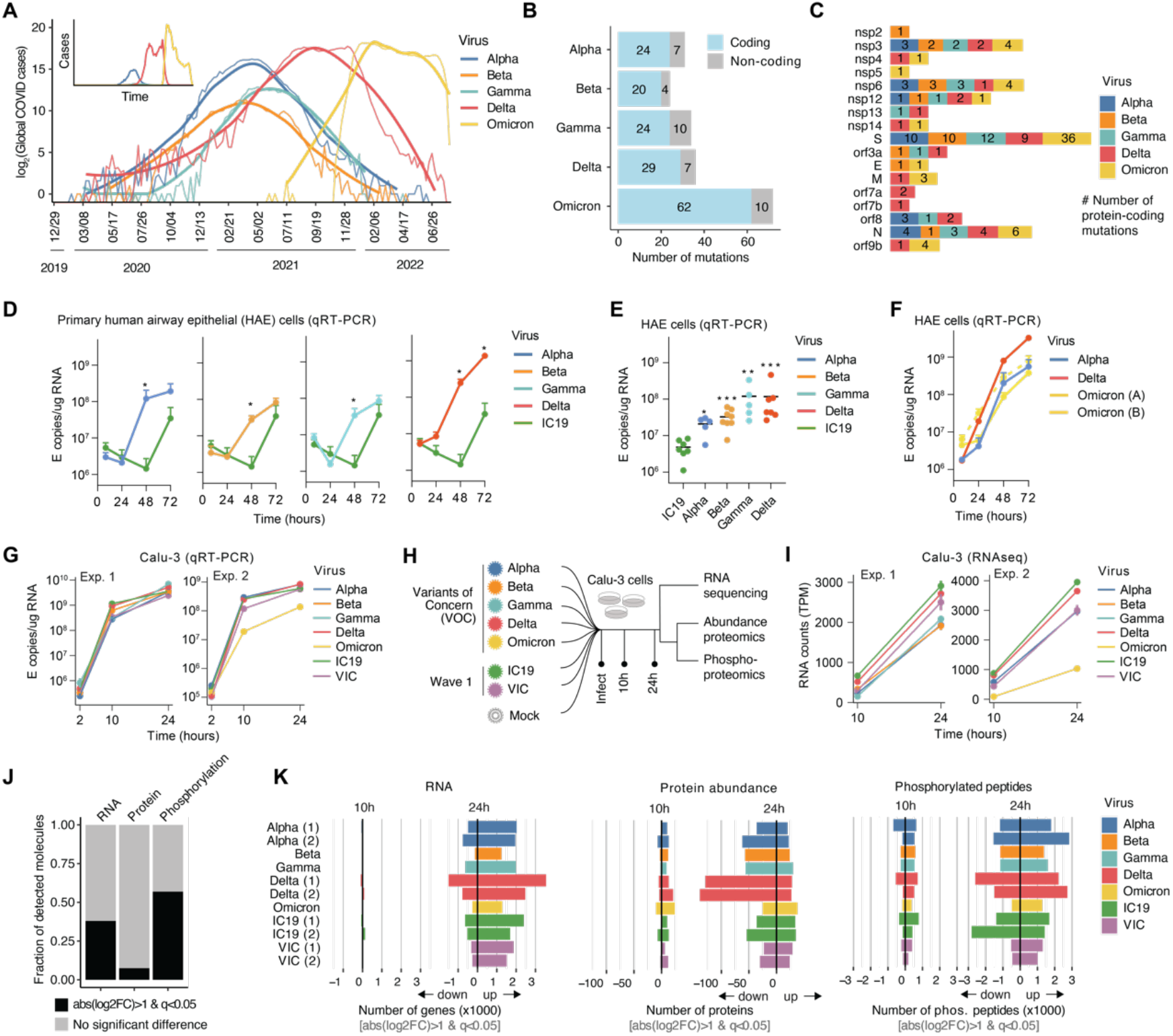
Emerged variants of concern impact RNA and protein landscape during infection. **(A)** Global COVID-19 case numbers (log_2_ transformed) over time, annotated for each SARS-CoV-2 variant of concern (VOC) based on sequences obtained from GSAID. Five VOCs are annotated: Alpha, Beta, Gamma, Delta, and Omicron BA.1. Thin lines reflect raw counts and thick lines represent a LOESS fit. Inset: y-axis is displayed in linear scale. **(B)** Number of protein coding and non-coding mutations for each VOC. Data were obtained from covariants.org on Jan 5,2022 and correspond to Alpha 20I V1, Beta 20H V2, Delta 21J, Gamma 20J V3, and Omicron 21K. **(C)** Number of protein coding mutations per SARS-CoV-2 viral protein in the VOCs (represented by colors). (**D-F**) Primary human airway epithelial (HAE) cells infected with 1500 E copies per cell of indicated VOCs. (D) Apical release of virus measured over time or (E) intracellular E levels at 72 hours post infection (hpi; n=5-7). (F) Apical release over time for Alpha, Delta and two independent Omicron BA.1 isolates. For D and E, VOCs were compared to IC19 by Mann-Whitney test at each time point. **(G)** Quantitative reverse-transcription PCR (qRT-PCR) for E viral transcript to quantify viral replication over time for each experiment. Calu-3 lung epithelial cells were infected with 2000 E copies of each SARS-CoV-2 VOC, two early-lineage wave 1 (W1) controls—IC19 (isolated from Europe in Spring 2020) and VIC (isolated from Victoria, Australia in Winter 2020)—and mock. **(H)** Experimental workflow. Infected Calu-3 cells were harvested at 10 and 24 hpi and processed for bulk mRNA sequencing and global mass spectrometry abundance proteomics and phosphoproteomics. Two independent experiments were performed with mostly overlapping conditions. Experiment 1 included Alpha, Beta, Gamma, Delta, IC19, VIC, and mock. Experiment 2 included Alpha, Delta, Omicron BA.1, IC19, VIC, and mock. Two sets of samples were generated in parallel for each experiment; one set of infected samples was processed for RNA and another set for both abundance proteomics and phosphoproteomics. **(I)** Viral replication over time in experiment 1 and 2 based on Orf1a leader sequence-containing counts from bulk mRNA sequencing. **(J)** Fraction of mRNA, protein, or phosphorylation sites that significantly change (black; abs[log2FC]>1 & q<0.05) in at least one virus infection, for at least one time point, in at least one experiment relative to mock. **(K)** Number of mRNA transcripts (left), proteins (middle), or phosphorylated peptides (right) that significantly increase or decrease for each condition and time point compared to mock. Numbers in parentheses next to viruses indicate the experiment number.

SARS-CoV-2 Spike mediates cell entry via host receptor ACE2 and is the most mutated VOC protein. However, SARS-CoV-2 VOCs have acquired additional non-synonymous mutations in non-structural, structural, and accessory proteins including Nsp3, Nsp6, Nsp9, Nsp12, Nsp13, nucleocapsid (N), membrane (M), envelope (E), Orf3a, Orf6, Orf7a, Orf7b, Orf8, and Orf9b. Each VOC has between 20-29 non-synonymous consensus mutations, with the exception of Omicron BA.1, which has 62 non-synonymous consensus mutations, largely localized within the Spike protein (Fig. 1B-C, S1A, Table S1). Each VOC also possesses between 4-10 non-coding mutations. We have previously shown adaptations outside of Spike can profoundly impact the host response to infection, reporting the Alpha variant evolved enhanced innate immune evasion by modulating specific viral protein expression levels (*14*), likely contributing to enhanced transmission. However, the impact of mutations outside of Spike in subsequent VOCs, including non-coding mutations, remains poorly understood. We propose that the development of prophylactic and therapeutic inhibitors, and effective management of the current and future coronavirus pandemics, will depend on a complete understanding of SARS-CoV-2 evolution and the host responses that variants provoke. Here, we report a comparative proteomic and genomic analysis of VOC phenotypes, host responses to VOC infections, and the impact of VOC evolution on virus-host interactions.

### Systems approaches reveal variants remodel the host landscape during infection

To understand the effect of VOC mutations on viral replication and cellular responses, we systematically compared SARS-CoV-2 variants during infection in human cells. Specifically, we compared the VOCs (Alpha, Beta, Gamma, Delta, and Omicron BA.1) and two early-lineage wave 1 (W1) isolates: (i) VIC, isolated from Victoria, Australia in Spring 2020 and (ii) IC19, isolated from Europe in Spring 2020, the latter harboring the universal Spike D614G mutation. We began by comparing viral replication in air-liquid-differentiated primary human airway epithelial (HAE) cells. Strikingly, we found that replication of each VOC outpaced W1 IC19, with Delta reaching the highest levels (Fig. 1D-E), consistent with VOC adaptive evolution. Furthermore, we found two independent Omicron isolates replicated comparably to Alpha in bronchial epithelial cells, but not effectively as Delta (Fig. 1F). Importantly, enhanced replication of VOCs was not observed in Calu-3 lung epithelial cells and quantitative reverse transcription PCR (qRT-PCR) detecting E RNA revealed equivalent replication kinetics for W1 virus and all VOCs, except for Omicron BA.1, which displayed a 5-fold reduction in replication (Fig. 1G). Calu-3 are therefore advantageous for comparing host responses (*14*), which can otherwise be confounded by large differences in replication, as observed in HAEs, as reduced infection typically diminishes the magnitude of the host response (*15*).

We prepared VOC-infected Calu-3 cells from two independent infections (termed experiment 1 and 2) harvested at 10 and 24 hours post infection for bulk mRNA sequencing (RNA-seq; Table S2-3), mass spectrometry abundance proteomics (Table S4-6), and phosphoproteomics (Table S7-9; Fig. 1H) analysis. Across both virus and host, our proteomics data captured 58,000-62,000 unique peptides (Fig. S2A) mapping to 4800-5100 proteins (Fig. S2B) per sample for abundance, and 23,000-35,000 unique phosphorylated peptides (Fig. S2C-D) mapping to 3700-4300 phosphorylated proteins (Fig. S2E), with strong reproducibility across biological replicates and experiments (Fig. S2F-G). We found viral replication kinetics to be reproduced at the RNA and protein level in the RNA-seq (Fig. 1I) and proteomics (Fig. S1B) datasets, as well as at single-cell resolution using flow cytometry (Fig. S1C). At the protein level, we noticed a slight decrease in protein production and/or viral spread for Beta in experiment 1. Of all detected molecules per dataset collected in Calu-3 cells, most changes during infection, relative to mock, were observed at the level of RNA (38%) and phosphorylation (57%) with fewer changes in protein abundance (7%; Fig. 1J). In general, we observed trends of RNA expression increases and protein expression decreases during infection relative to mock at 24 hours post infection, typically representing the induction of the inflammatory response to virus at the host RNA level and a virus-induced host translational blockade known to occur during SARS-CoV-2 infection at the protein level (*16*–*18*) (Fig. 1K). Regulation at the phosphorylation level was observed to be equivalently bi-directional, reflecting the complexity of phosphorylation signaling responses during viral infection (Fig. 1K).

### Variants evolved to modulate viral RNA and protein production

To understand the functional effect of VOC mutations, we first quantified differences in viral RNA and protein levels across the VOCs during infection in Calu-3 cells (Table S10). We normalized each viral RNA quantity to levels of positive-sense single-stranded RNA genome for each virus (e.g. “genomic”) and each viral protein quantity to the sum of all non-structural protein intensities, intended to control for differences in viral replication (Fig. 2A). Next, a log_2_ fold change (log2FC) was calculated to compare each virus to our W1 control VIC for both viral RNA and protein (see Methods). We observed large differences in the expression of subgenomic (sg) transcripts and proteins corresponding to structural and accessory genes during VOC infection (Fig. 2B-C, S3A). However, large differences in non-structural protein (Nsp) levels were not observed across the viruses (Fig. 2C), as expected given they are translated as a polyprotein and consistent with the notion that coronavirus evolution is associated with the diversification of genes encoded by sgRNAs (*19, 20*). Interestingly, viral RNA and protein expression did not always correlate (Fig. 2D, S3B), suggesting independent regulation of transcription and protein expression (translation and/or protein stability). This was especially evident for Orf6 expression by Omicron BA.1, which showed up-regulated sgRNA but down-regulated protein levels relative to VIC (Fig. 2B-C, S3B). This was also evident for Orf9b expression by Alpha and Delta, which both showed increased protein levels but only Alpha had upregulated Orf9b-specific sgRNA levels compared to VIC (Fig. 2E, S3A-B).

**Figure 2.**
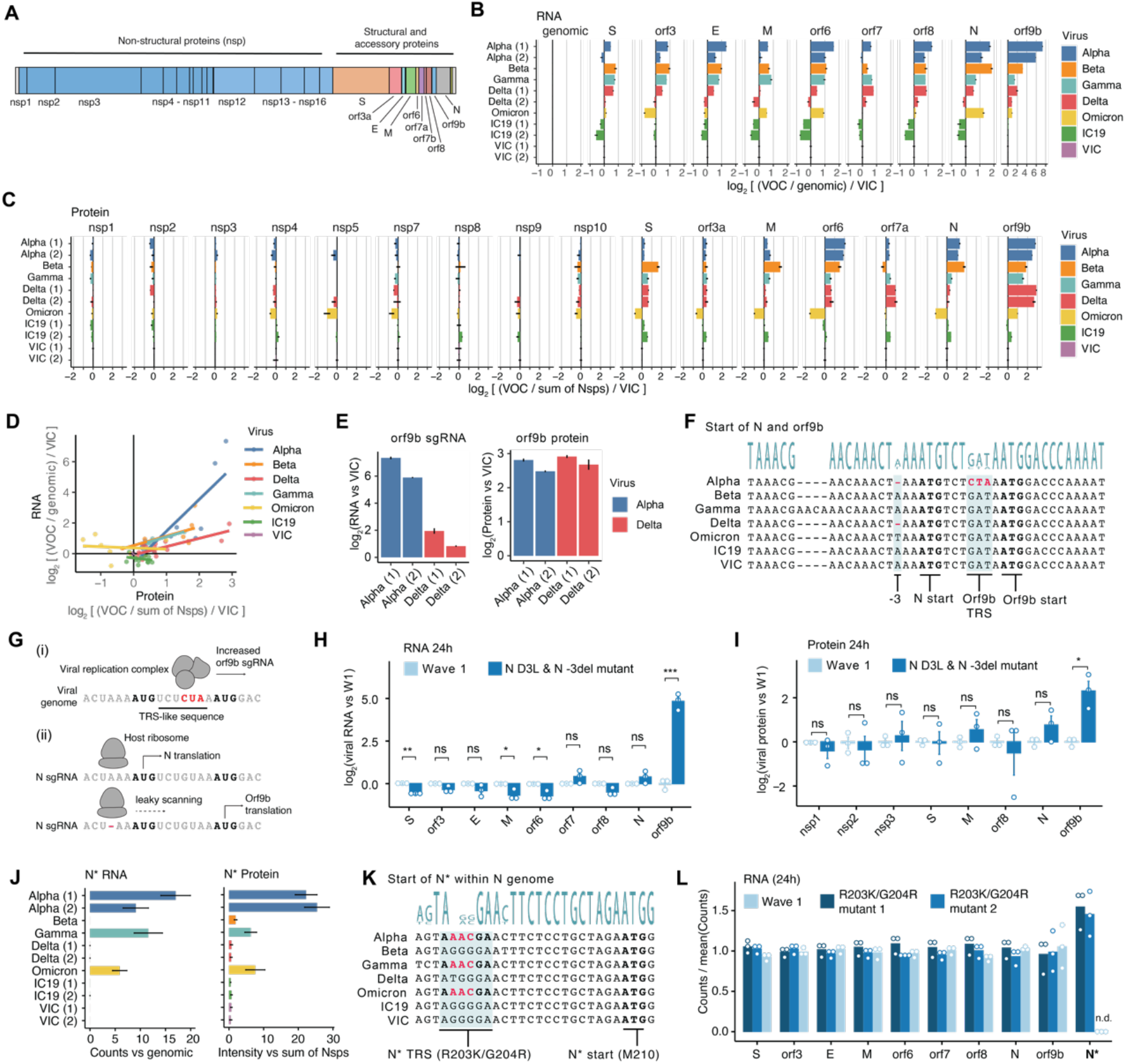
Variants of concern acquired mutations to modulate viral RNA and protein production. **(A)** Map of SARS-CoV-2 genome with each protein annotated. Blue represents the Orf1a/b gene, translated as either a shorter (Orf1a) or longer (Orf1ab) polyprotein and cleaved into 11 or 16 non-structural proteins (Nsps), respectively. Other colors represent the structural and accessory proteins, which are translated from viral subgenomic RNA, including Spike (S), Orf3a, envelope (E), membrane (M), Orf6, Orf7a, Orf7b, Orf8, nucleocapsid (N), and Orf9b. **(B)** Normalized read counts from bulk mRNA sequencing containing the leader sequence and mapping to a portion of the SARS-CoV-2 genome at 24 hours post infection (hpi) in Calu-3 cells. Each variant genome was reconstructed from bulk mRNA sequencing data obtained during infection. Counts for each viral gene were normalized to genomic (i.e. Orf1a transcript) counts for each virus, to account for differences in viral replication. This value was additionally normalized to VIC independently for each viral gene to depict changes in gene expression relative to a W1 isolate. Experiments 1 and 2 are annotated in parentheses where applicable. **(C)** Normalized protein intensities from global abundance proteomics. Similarly as for RNA, summarized protein intensities for each viral protein were first normalized to the sum of all detected Nsps per virus and then to VIC for each viral protein separately. Experiments 1 and 2 are annotated in parentheses where applicable. **(D)** Calculated VOC- and VIC-normalized viral RNA and protein quantities from (B) and (C) displayed as a scatter plot, fit with a linear regression line to capture the relationship between changes in RNA and protein expression across the VOCs relative to a W1 isolate (VIC). **(E)** The quantity of SARS-CoV-2 Orf9b viral RNA and protein for Alpha and Delta (across experiments 1 and 2) isolated from (B) and (C). **(F)** Genomic sequence of the region surrounding the start of the nucleocapsid (N) gene for viruses in this study. The “-3” indicates 3 nucleic acid positions upstream of the N translation start (“N start”). The Orf9b translation start (“Orf9b start”) is also indicated within the N coding region. Mutations present in Alpha (-3 deletion and N protein D3L) and Delta (-3 deletion only) are indicated in red. The Orf9b transcriptional regulator sequence (TRS) is also indicated and is thought to be enhanced by the GAU→CUA. **(G)** Cartoon depiction of how mutations colored red in (F) are thought to affect transcription and translation of Orf9b. Transcription of Orf9b sgRNA (i) occurs when the viral replication complex binds the Orf9b TRS. The GAU→CUA mutation enhances this TRS, increasing the production of Orf9b sgRNA. However, the majority of Orf9b translation (ii) is thought to occur from leaky scanning of the host ribosome along the N sgRNA. When the Kozak context for N is weakened by the -3 deletion (ii, bottom), ribosomal leaky scanning increases in frequency, driving increased Orf9b protein produced from the N sgRNA. **(H)** Expression of viral sgRNA from bulk mRNA sequencing at 24 hpi in A549-ACE2 cells with an early-lineage SARS-CoV-2 (wave 1) derived from Washington, USA (USA-WA1/2020), and one with two mutations, the D3L change in N (GAU→CUA) and a deletion at the -3 position upstream from the translation start site for N, generated by reverse genetics. Quantifications are normalized to genomic (leader + Orf1a) counts per virus to control for differences in viral replication and defined relative to wave 1 virus (log2(counts/genomic/W1)). **(I)** Expression of viral proteins from mass spectrometry abundance proteomics with the same conditions as in (H). Quantifications are normalized to the summed intensity of all Nsps per virus to control for differences in viral replication and defined relative to W1 virus (log2(intensity/sum of nsps/W1)). **(J)** Expression of SARS-CoV-2 N* (N-star) subgenomic RNA from bulk mRNA sequencing at 24 hpi in Calu-3 cells for all viruses in experiments 1 and 2. N* is a viral protein encoded within, and in-frame with, the N protein sequence, containing the C-terminal part of N from amino acid position 210 to the end of N. **(K)** Genomic sequence of viruses under study, highlighting the region surrounding the translation start site of N*. Mutations within the N* TRS in Alpha, Gamma, and Omicron (corresponding to the R203K/G204R mutations in N) are predicted to enhance the N* TRS leading to increased N* sgRNA production. **(L)** Expression of viral subgenomic RNA from bulk mRNA sequencing at 24 hpi in Calu-3 cells with an early-lineage SARS-CoV-2 (W1) from Wuhan and two individual attempts at constructing a virus with a GGG→AAC mutation (corresponding to the R203K/G204R in N), created using reverse genetics approaches. Quantifications are normalized to genomic (leader + Orf1a) counts per virus to control for differences in viral replication and defined relative to the average of wave 1, mutant 1, and mutant 2 (counts/mean(counts)) per viral gene. It was not possible to normalize or statistically compare to wildtype because zero N* counts were detected in the wildtype condition (n.d.). Statistical significance testing for (H) and (I) was performed using a two-tailed t-test assuming unequal variance. *, p<0.05; **, p<0.01; ***, p<0.001. ns=not significant; n.d.=not detected. Error bars represent the standard error (SE).

We and others previously reported increased Orf9b expression during Alpha infection (*14, 21*) and hypothesized that Orf9b sgRNA and protein expression could be controlled by distinct mutations. Orf9b is located in an alternative reading frame within N. Alpha encodes an N D3L mutation (Fig. 2F), predicted to enhance a transcriptional regulatory sequence (TRS) driving increased Orf9b-specific sgRNA expression (Fig. 2G). It also contains a single nucleotide deletion at position -3 upstream of N (Fig. 2F), which we predicted influences N Kozak context and therefore Orf9b translation by increasing leaky ribosomal scanning and Orf9b translation from the N sgRNA (Fig. 2G). Comparatively, Delta contains the -3 deletion but not N D3L (Fig. 2F), in line with increased Orf9b protein but not sgRNA (Fig. 2E). The similar increase in Orf9b protein levels in Alpha and Delta suggests Orf9b protein expression results predominantly from leaky scanning of N sgRNA to produce Orf9b protein driven by the -3 Kozak change, rather than from translation of Orf9b-specific sgRNA. This is in accordance with our previous observation that Alpha Orf9b transcript levels are in low abundance compared to N sgRNA levels (*14*). Notably, Omicron BA.1 contains an A-T mutation in the -3 N position (Fig. 2F), which we predict also weakens the N Kozak context, and as such results in modest upregulation of Orf9b protein (Fig. 2C). To probe regulation of Orf9b expression we generated a reverse genetics mutant virus with both N D3L and -3 A deletion in a W1 background (USA-WA1/2020 background, see Methods). Global mRNA and proteomic analyses during infection of lung epithelial A549-ACE2 cells with this virus revealed Orf9b to be specifically upregulated at both the sgRNA and protein level (Fig. 2H-I, S3C-E), as expected. Interestingly, Beta and Gamma also display upregulated Orf9b protein (Fig. 2C), though at lower levels compared to Alpha and Delta, but have neither of the mutations discussed herein (Fig. 2F), suggesting additional mechanisms regulating Orf9b expression.

In addition, we and others have reported the expression of a new sgRNA and protein of the Alpha VOC, called N* (“N-star”) (*14, 21*), a truncated version of N starting at amino acid position 210 and continuing until the end of N. Here, we confirmed N* sgRNA and protein to be specifically expressed by Alpha, Gamma, and Omicron (Fig. 2J). N* sgRNA was expressed at low abundance compared to N but similar to Orf6 and E (1-2% of viral transcripts; Fig. S3F). The R203K/G204R double substitution in N is found in the B.1.1 lineage, which includes Alpha, Gamma, and Omicron, and is hypothesized to regulate N* expression because the corresponding nucleotide changes (GGG→AAC), overlapping two codons, create a novel TRS (Fig. 2K). To test this hypothesis, we generated a reverse genetics mutant virus with the R203K and G204R mutations (GGG→AAC) in a W1 background from Wuhan. Global mRNA sequencing during infection in Calu-3 cells confirmed specific N* expression by the R203K/G204R mutant but not by the parental control virus (Fig 2L). However, N* expression was approximately half of that observed during Alpha infection (Fig. S3G), despite normalization to infection levels, suggesting additional mechanisms may contribute to N* expression or that Alpha upregulation of N sgRNA also results in increased N* transcripts.

We next used a SARS-CoV-2 virus-like particle (VLP) system (*22*), which contains viral structural proteins N, S, M, and E and a reporter genome encoding luciferase, to examine how alterations in structural proteins affected VLP production and infectivity (Fig. S3H, see Methods). We found W1 N produced the least infectious units when combined with W1 structural proteins (Fig. S3I). Omicron N produced >10x more infectious units than W1 N, as previously shown (*23*), while Alpha, Gamma, Delta and Beta N-bearing VLPs produced intermediate amounts (Fig. S3I). This suggests that all the VOCs evolved mutations in N protein to enhance viral packaging, likely affecting the production of infectious particles and infectivity. We also evaluated the effects of N* on packaging and infectivity using this assay. Surprisingly, we discovered that N* can package RNA and produce infectious VLPs (Fig. S3J) even though it does not contain the canonical N-terminal RNA binding domain, suggesting alternative mechanisms of viral packaging, perhaps related to phase separation (*24*).

### Host kinases modulate viral protein phosphorylation during infection

We have previously shown that phosphorylation of viral proteins can regulate their activity (e.g. Orf9b) (*14*). Therefore, we systematically compared viral protein phosphorylation across the VOCs during infection (Table S10). Across all viruses, we detected 53 viral protein phosphorylation sites, mapping to Nsp1, Nsp3, S, M, N, and Orf9b (Fig. 3A), similar to our previous report (*18*). We quantitatively compared phosphopeptide intensities, normalized to total protein abundance, between all possible virus pairs. Importantly, we only compared peptides with identical sequences between each virus pair to control for physicochemical peptide properties that may affect their quantitation using mass spectrometry (see Methods).

**Figure 3.**
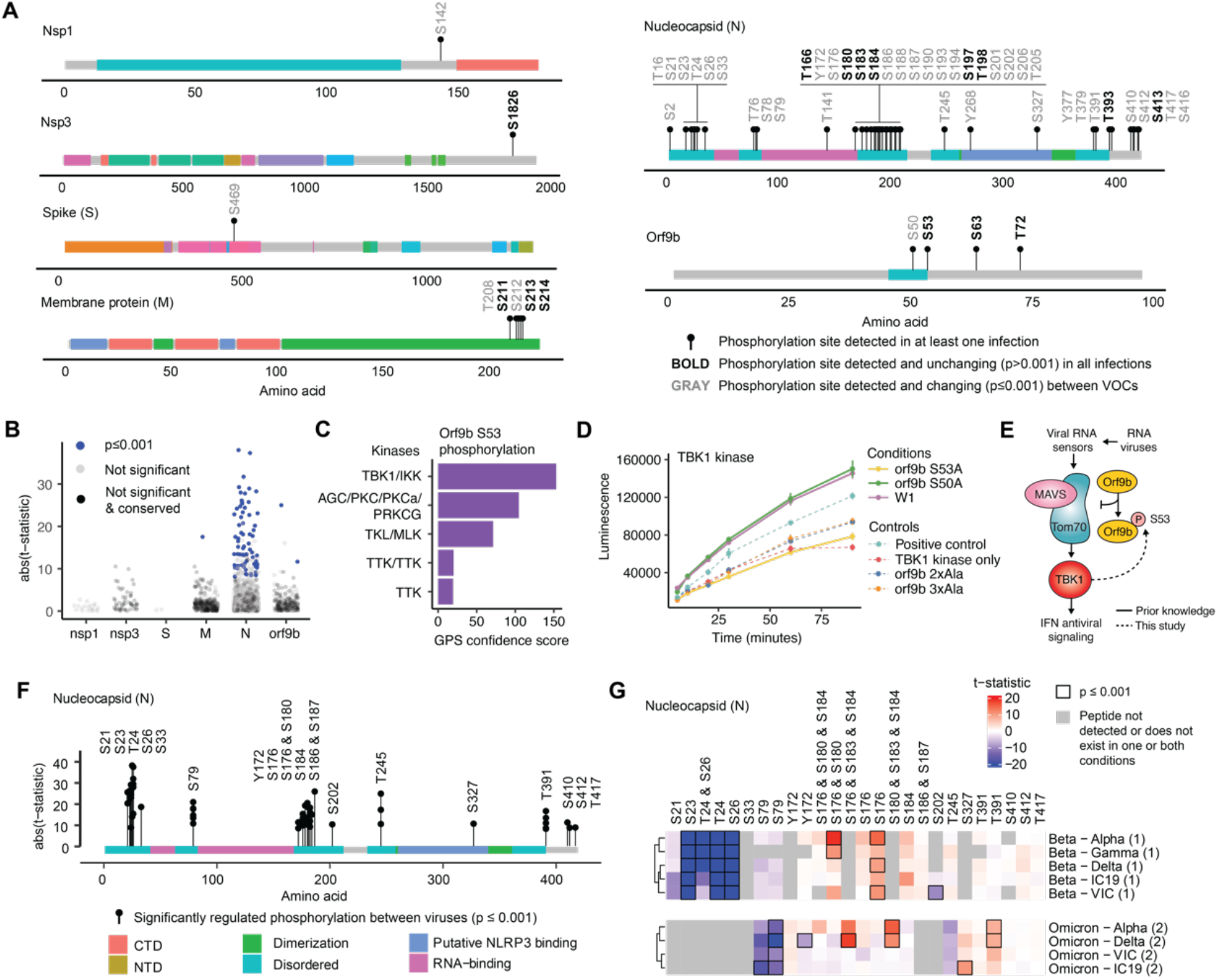
Host kinases modulate viral protein phosphorylation during infection. **(A)** Global phosphoproteomics map of viral protein phosphorylation sites detected during infection with at least one virus (Alpha, Beta, Gamma, Delta, Omicron BA.1, VIC, or IC19) in Calu-3 cells (black lollipops). Bold site labels indicate phosphorylation sites detected in all infections and with total protein abundance-normalized phosphorylation site intensities unchanging (p>0.001) between viruses. Gray site labels indicate phosphorylation sites detected and changing (p≤0.001) between at least one pair of VOCs (see Table S10 and Methods). Colors delineate known protein domains. **(B)** The absolute value t-statistic from t-tests comparing phosphorylated peptide intensities between pairs of viruses. Each dot represents one phosphopeptide compared between two viruses. Phosphorylated peptide intensities are normalized by the corresponding total protein abundance. Comparisons were restricted to viral peptides with identical sequences between virus pairs, given that peptide intensities of peptides with different sequences are not directly comparable using mass spectrometry. If p≤0.001, dots are colored dark blue, comparisons that are not significant are colored gray, comparisons that are not significant but are detected for all viruses are colored black (conserved, as in A). **(C)** Top five kinase groups predicted to phosphorylate S53 in Orf9b using GPS 5.0 (*25*), ranked by GPS confidence score. **(D)** TBK1-mediated phosphorylation of Orf9b measured using an *in vitro* ADP-Glo kinase time course. An increase in luminescence indicates increased peptide phosphorylation. Peptides were synthesized to reproduce the area surrounding Orf9b S53, including 7 amino acids downstream of S50 and 7 amino acids upstream of S53. Orf9b 2xAla indicates positions S50 and S53 were converted to non-phosphorylatable alanine. Orf9b 3xAla indicates positions S50, S53, and T70 were converted to alanine. Myelin basic protein was used as a positive control. TBK1 kinase only indicates the addition of kinase without any peptide substrate. **(E)** Cartoon schematic of proposed Orf9b-TBK1 signaling pathway. It was previously known that viral RNA is detected by host RNA sensors, which activate the MAVS-Tom70 complex and TBK1, leading to the upregulation of interferon stimulated genes. We previously reported (*14*) Orf9b to bind to Tom70 and antagonize innate immune activation, but only when not phosphorylated at S53 (black lines). Here, we report TBK1 is able to phosphorylate Orf9b S53, revealing a negative feedback loop between Orf9b and TBK1. **(F)** Significantly changed phosphorylation sites (p≤0.001) on N protein between pairs of viruses. Length of each lollipop depicts the abs(t-statistic) between the pairs of viruses. **(G)** Heatmap of significantly changed phosphorylated peptides (p≤0.001) mapping to N protein depicted in (F) relative to Beta and Omicron (see Fig. S4B for full heatmap). Rows indicate the comparison (first virus is numerator, second is denominator) and columns indicate the phosphorylated peptides, and residues, on N. Color represents the t-statistic and black bounding boxes indicate p≤0.001. Gray boxes indicate peptides that were either not detected in one or both viruses or did not possess identical sequences between viruses. Numbers next to viruses indicate experiment 1 or 2.

Twenty eight percent (15/53) of sites were conserved across all viruses, meaning they were detected during infection with every virus and were unchanging (p>0.001) at all time points between any pair of viruses (Fig. 3A-B). Conserved phosphorylation sites likely reflect regulatory interactions that are central to the SARS-CoV-2 life cycle. One conserved site was Orf9b S53, which we additionally confirmed to be unchanging across viruses using targeted proteomics (Fig. S4A). We previously reported that Orf9b S53 phosphorylation can inhibit its binding to human TOM70, disrupting Orf9b-mediated innate immune antagonism (*14*). Using the GPS computational algorithm (*25*), we found TBK1, a kinase known to activate the innate immune response downstream of TOM70 and MAVS, to be the top candidate kinase for Orf9b S53 phosphorylation (Fig. 3C). Using an *in vitro* ADP-Glo kinase assay, we confirmed TBK1 can robustly phosphorylate Orf9b exclusively at S53 and not S50 or T60 (Fig 3D). Interestingly, this finding reveals a negative feedback loop between Orf9b and TBK1 (Fig. 3E); while Orf9b can inhibit TBK1 activation through TOM70, TBK1 can also block the interaction between Orf9b and TOM70 by phosphorylating Orf9b S53. These findings suggest that activation of TBK1 during infection may provide a mechanism to reverse Orf9b mediated innate immune antagonism, promoting inflammatory responses after viral sensing. Orf9b may therefore sense TBK1 activity and act as a molecular switch that responds to innate immune activation, regulating it in both directions depending on timing. However, more work is needed to understand exactly how cellular phosphorylation signaling is manipulated by SARS-CoV-2 infection and how this regulates infection.

Forty three percent (23/53) of detected sites were differentially regulated between viruses (p≤0.001; see Methods), mostly on N protein (Fig. 3B, 3F). Differential regulation was observed across the N protein surface, with clusters localized at the N-terminal region (S21-S33) and linker region after the RNA binding domain (Y172-S202; Fig. 3F). Interestingly, we detected a strong mutant-specific decrease in N-terminal (S23-S26) phosphorylation on N protein for Beta compared to all other viruses (Fig. 3G, S4B; Omicron BA.1 was excluded due to mutations in this region). Sequence-based kinase-substrate motif analysis suggested Polo-Like Kinase (PLK) or Casein Kinase could phosphorylate these sites (Fig. S4C). Interestingly, our APMS data (described later) identified a complete loss of interaction between Beta N protein and CSNK2B, the regulatory subunit of Casein Kinase II (see Fig. 4D, S6). We also observed increased phosphorylation of the Beta and Omicron N linker region (S176, S183, and S184) and decreased phosphorylation of Omicron N S79 compared to other viruses (Fig. 3G, S4B). Previous studies have shown that phosphorylation of N protein impacts RNA binding, packaging, and viral assembly (*24, 26, 27*), suggesting that differential regulation of N phosphorylation by VOCs could affect these stages of the viral life cycle, perhaps even influencing viral genome sensing.

**Figure 4.**
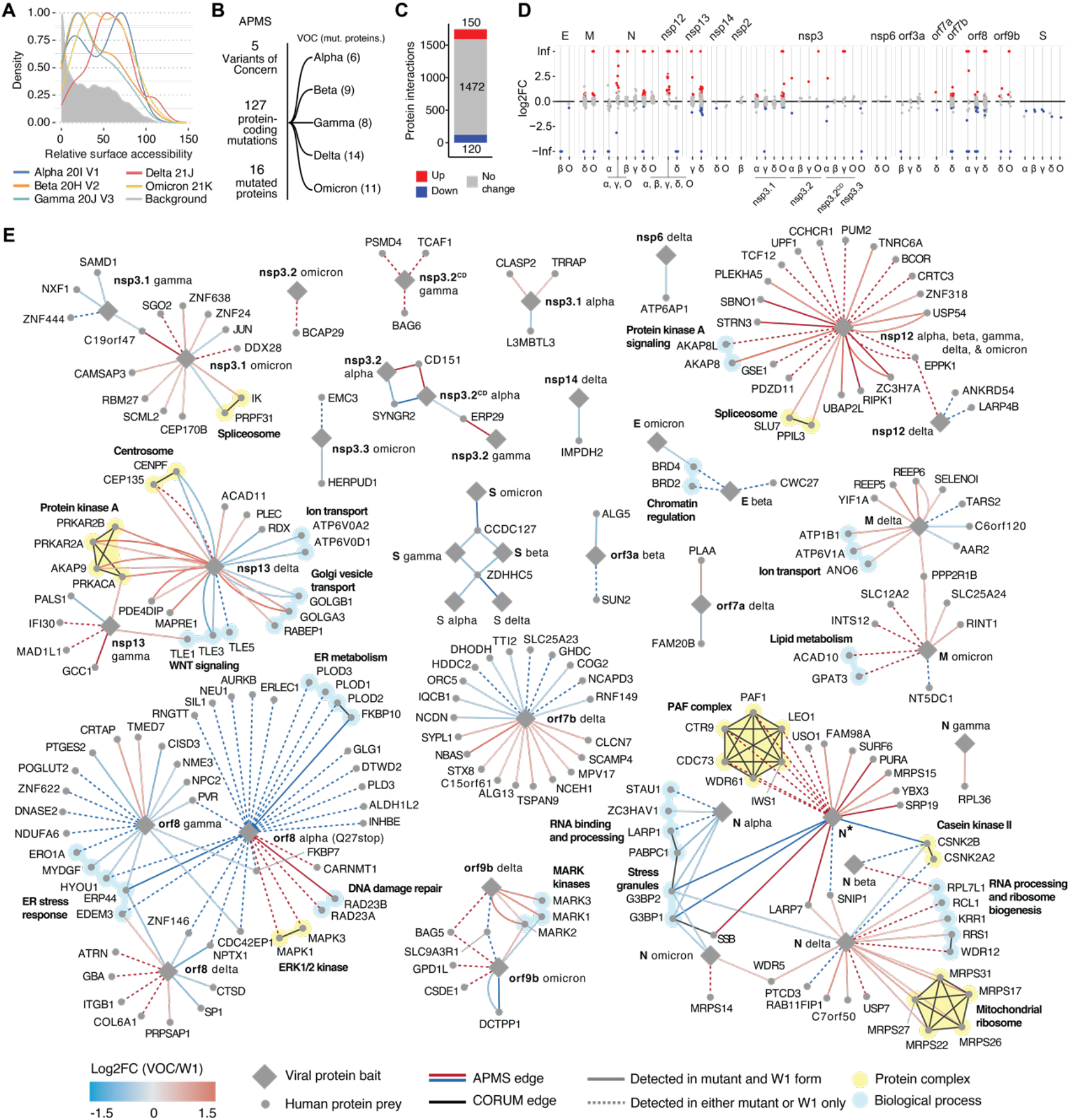
Mutations in variant proteins rewire virus-host protein interactions. **(A)** Density distribution of naccess surface accessibility calculations for mutations within the VOCs (colored lines) compared to all possible mutations (background, gray shaded region). Only viral proteins with solved structures were included. Shift to the right indicates increased surface accessibility for mutations in VOCs compared to the background. **(B)** Schematic depicting numbers of VOCs (5), protein-coding mutations (127), total mutated viral proteins (16), and the breakdown of those mutations among the individual VOCs. **(C)** Number of high confidence virus-human interactions for all mutant VOC and wave 1 (W1) viral proteins using affinity purification mass spectrometry (APMS). Of 1746 total interactions, 1473 were unchanged between mutant and wildtype (gray), 150 increased in binding affinity with at least one mutant (red), and 123 decreased in binding affinity with at least one mutant (blue). Significant increase in mutant binding affinity (red) is defined as log_2_ fold change>0.5 & p<0.05. Significant decrease in mutant binding affinity (blue) is defined as log_2_ fold change<-0.5 & p<0.05. **(D)** Jitter plot showing log_2_ fold changes (log2FC) of interactions conserved (gray) between VOC and W1, increased in binding to mutant (red) and decreased in binding to mutant (blue), for each viral “bait” protein. Positive and negative infinity log2FC values indicate interactions detected in mutant or W1 only, respectively. Red and blue annotation is as defined in (C). **(E)** Protein-protein interaction (PPI) map for significantly changing interactions (absolute value log_2_ fold change>0.5 & p<0.05) comparing VOC to W1 forms. When multiple mutations occur in a VOC viral protein, they are merged into one construct (see Table S11). Diamonds and circles indicate viral “bait” and human “prey” proteins, respectively. Color of edge represents log_2_ fold change in the abundance of each human prey protein in the affinity purification as determined by mass spectrometry, comparing VOC and W1 forms. Human preys detected in both VOC and W1 are indicated with solid edges whereas preys detected in either VOC or W1 only are indicated with dashed edges. Multiple edges are displayed when the same affinity purification was performed multiple times and both results were significantly differentially interacting. Black edges indicate human-human protein complexes annotated by CORUM (*28*), also highlighted and annotated using yellow shading. Biological processes are indicated using cyan shading.

### Protein-coding mutations in variants rewire virus-host protein interactions

Global replacement of one SARS-CoV-2 VOC by another is presumably due to enhanced human-to-human transmission driven by the unique constellation of adaptive mutations within each VOC. To comparatively assess the functional consequences of VOC adaptation, we performed an integrative sequence and structural analysis of thousands of mutations cataloged by GISAID (Fig. 4A, S5A). Across the five VOCs, there are a total of 127 non-synonymous mutations spanning 16 viral proteins across Alpha (6 viral proteins), Beta (9), Gamma (8), Delta (14), and Omicron BA.1 (11) (Fig. 4B). Our analysis predicts mutations found within the variants to be structurally well-tolerated at the level of protein folding and unlikely to dramatically disrupt viral protein structure (Fig. S5A), which is expected given a core disruption would likely be disadvantageous to the virus. Conversely, we found VOC mutations were more likely to be surface accessible (Fig. 4A), suggesting they may play a role in modulating virus-host protein-protein interactions (PPIs).

To assess the impact of protein-coding mutations on virus-host PPIs, we systematically performed affinity purification mass spectrometry (APMS) on all viral proteins mutated in the VOCs compared to Wuhan reference virus (W1; Table S11). Briefly, we overexpressed and affinity purified VOC and W1 viral proteins with 2x strep-tags from HEK293T cells and performed mass spectrometry analysis, as previously described (*29, 30*). SAINTexpress (*31*) and MiST (*32*) scoring algorithms were used to define high-confidence interactions in either the VOC or W1 counterpart (Fig. S5B-C). Next, MSstats (*33*) was used to quantitatively compare the abundance of each human protein “prey” within each affinity purification between VOC and W1 “baits”, generating a log_2_ fold-change and p-value for each interaction. Of 1742 high-confidence interactions, 270 (16%) were found to be significantly differentially interacting between VOC and W1 forms (abs(log2FC)>0.5 & p<0.05), 150 with increased (red) and 123 with decreased (blue) affinity for the VOC form (Fig. 4C). Differential interactions were observed for most viral proteins, except for Nsp2 and Nsp6 (Fig. 4D). We additionally profiled a variety of individual mutations or mutation subsets across the VOCs to help pinpoint how specific mutations impacted the observed changes (Fig. S6). We visualized interactions defined as significantly differentially interacting (abs(log2FC)>0.5 & p<0.05) as a network, annotated with known host protein complexes (yellow) and biological processes (cyan; Fig. 4E).

Given that N* is lowly expressed, accounting for only 1-2% of total viral sgRNA reads compared to 15-55% for full length N (Fig. 3F), we hypothesized it must possess an additional novel function other than packaging viral genome. Indeed, the differential PPI map revealed a novel and specific association between N* and the RNAPII-associated factor (PAF) complex, specifically complex members PAF1, LEO1, IWS1, WRD61, CDC73, and CTR9 (Fig. 4E) (*34*–*36*), an interaction we confirmed using AP-western blot analysis (Fig. S5D). Strikingly, PAF interactions were completely absent from any of the full-length VOC or W1 N affinity purifications, consistent with this interaction representing a function not present in the zoonotic W1 virus. Specifically, this interaction suggests a novel function in the regulation of host gene expression during infection of N*-expressing VOCs (Alpha, Gamma, and Omicron), perhaps similar to a feature of influenza NS1 protein (*37*).

Other interactions of interest include several VOC N proteins (Alpha, Beta, Delta, and Omicron) showing decreased affinity for G3BP1/2 and Casein Kinase II (Fig. 4E). Additionally, the interaction between Delta Orf9b and all three MARK kinases (MARK1-3) was increased, which we validated by AP-western blot analysis for MARK2 (Fig. S5E). Other differential interactions of note include a strongly increased interaction between Alpha Nsp3 and the tetraspanin CD151, a host factor that could promote coronavirus infection (*38, 39*), and an increased interaction between Delta and Gamma Nsp13 and protein kinase A. A decreased interaction between Beta and Omicron E and BRD2/4 was also discovered, which may suggest differences in E protein acetylation (*40*). Finally, Alpha Orf8 lost the majority of its interaction partners, which is expected given that a mutation introduces a stop codon at position 27. However, it also gained several interactions including with RAD23A/B and MAPK3/1 (ERK1/2), suggesting new functionality (Fig. 4E). This analysis provides a holistic view of how VOCs are adapting to host by altering their host-virus interactions.

### Systems analysis reveals conservation and divergence of host response to variants

We next sought to understand how the VOCs impact host cellular biology during infection by developing an integrative computational analysis of our systems level datasets and ranking the modulation of host cellular processes across the VOCs to (i) reveal therapeutic targets for pan-variant antivirals by identifying similarly-hijacked cellular processes and (ii) uncover how viral evolution has led to divergent host responses. Compiling differentially regulated host genes and proteins (as compared to mock) from our mRNA sequencing, abundance proteomics, and phosphoproteomics datasets, resulted in 5284 genes regulated during infection (Fig. 5A, see Methods). We overlaid regulated genes onto the STRING network (*41*) and used measures of network proximity to cluster genes into 85 separate pathway modules (Fig. 5B, S7A, Table S12; see Methods). The magnitude of gene regulation per module was quantified as the average absolute value log_2_ fold change of genes within each module, separately for RNA, protein abundance, and phosphorylation, generating a single metric per module, per virus (Fig. 5C). Interestingly, we noticed most modules followed a general trend, which we defined as the Average Host Response (AHR; average regulation across modules; Fig. 5C). For example, the AHR was more pronounced during Delta compared to VIC infection. As expected, the AHR correlated well with the amount of viral RNA (genomic reads; R=0.59) and protein (sum of non-structural proteins; R=0.77) produced for each virus, suggesting the AHR to represent a generalized cellular response to infection, proportional to the amount viral replication (Fig. 5D).

**Figure 5.**
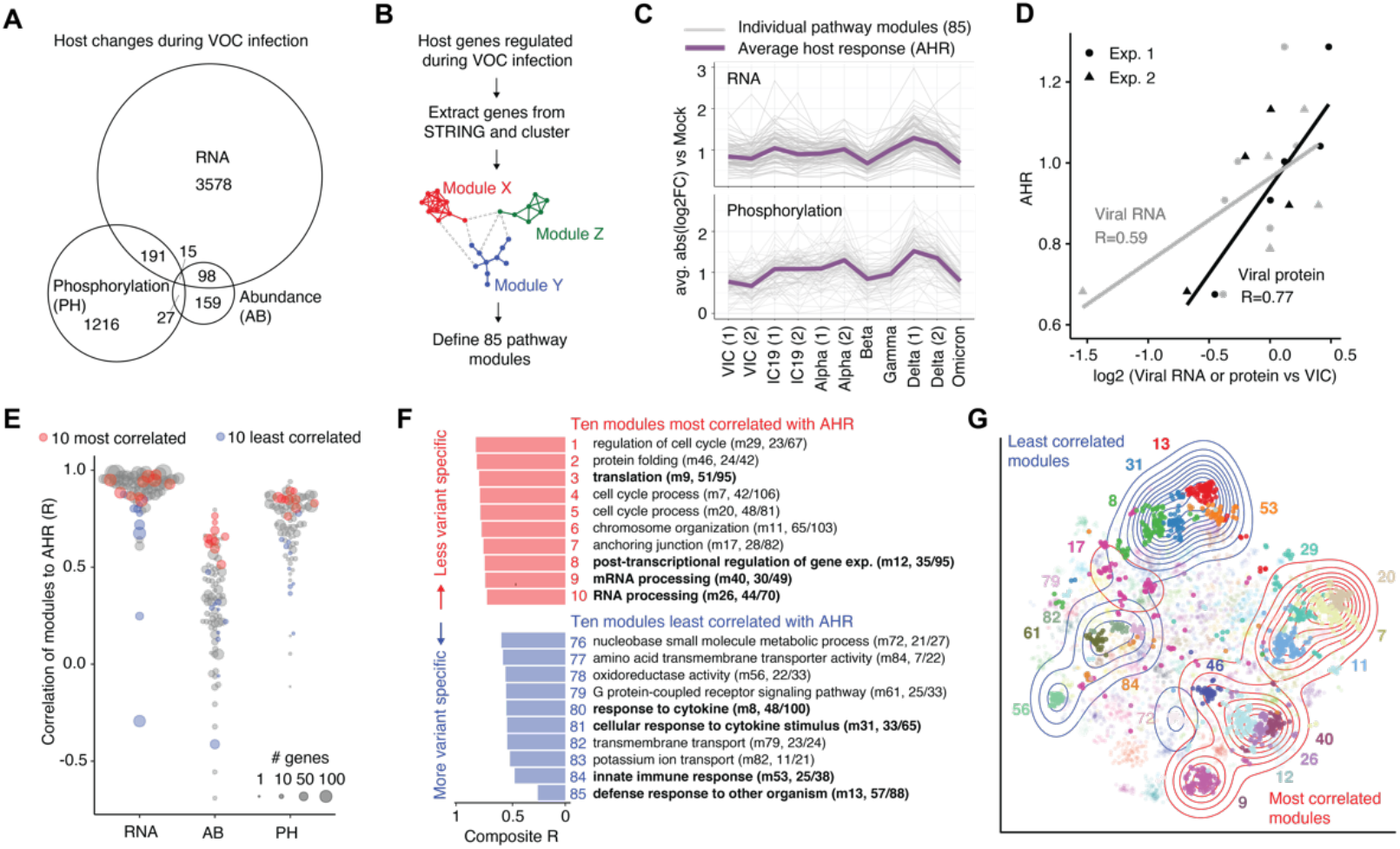
Integrative computational analysis reveals conservation and divergence of host response to variants. **(A)** Number of RNA transcripts, proteins, or phosphorylation sites that significantly changed during VOC infection compared to mock at 24 hpi. For transcriptomics and phosphoproteomics, we required absolute value log_2_ fold change (log2FC)>1 and p<0.001. For abundance proteomics, we required abs(log2FC)>log_2_(1.5) & q<0.05. For each dataset, a molecule had to pass the threshold twice at either different time points, viruses, or experiments. **(B)** Flowchart of computational pipeline. Host genes regulated during VOC infection from (A) were extracted from the STRING network and clustered into 85 pathway modules based on a diffusion measure of network node proximity (see Methods). **(C)** The average absolute value log_2_ fold change versus mock for each module (gray lines) using RNA or phosphorylation data. Virus conditions are ordered according to their timeline of emergence around the world. The purple line defines the average across the module intensities to define the Average Host Response (AHR). **(D)** Pearson’s R correlation between viral genomic (leader + Orf1a) RNA counts (R=0.59) or sum of non-structural protein (Nsps) intensities (R=0.77) versus VIC plotted against the AHR defined in (C). Each dot represents one virus condition at 24 hpi. **(E)** Pearson’s R correlation between the average log_2_ fold change of each module, per dataset, and the RNA-derived AHR. Red colored dots indicate the 10 modules most correlated with the AHR based on a geometric mean across RNA, abundance proteomics, and phosphoproteomics datasets which we term the “composite R”. Similarly, blue dots indicate the 10 modules least correlated with the AHR. **(F)** The 10 most (red) and 10 least (blue) correlated modules with the AHR, annotated by the most prevalent GO Biological Process term, module number, and number of genes within the module that connect to the top GO term. The x-axis depicts the composite R value (defined in E). Colored numbers indicate ranking of modules based on composite R. Terms in bold highlight prevalent biological categories: translation related terms in top 10 and innate immune/inflammation related terms in the bottom 10. Red modules represent pathways similarly regulated by all variants (“less variant specific”). Blue modules represent pathways differently regulated across the variants (“more variant specific”). **(G)** A t-Distributed Stochastic Neighbor Embedding (t-SNE) plot representing the STRING network proximity between genes, colored according to the module annotation. The top (red) and bottom (blue) 10 modules are bolded and their locations are annotated using contours, revealing the two groups to be well separated in multidimensional space.

We utilized the AHR to identify similarities and differences between VOCs. Systematic correlation of each module versus the AHR revealed most processes to be well correlated (Fig. 5C, 5E), highlighting the array of host pathways modulated in concert with a general cellular response to viral infection. Some modules correlated poorly with the AHR, representing VOC-specific differences, above-and-beyond a generalized cellular response, allowing us to hone in on key differences between the variants. The ten most correlated modules were related to cell cycle, protein folding, RNA translation and processing, chromosome organization, and the cytoskeleton (Fig. 5F, top). The ten least correlated modules included defense response to foreign organisms, innate immune and cytokine response, ion transport, GPCR signaling, and metabolic pathways (Fig. 5F, bottom). The functional distinction between the most and least correlated modules was corroborated by visualizing them on a t-distributed stochastic neighbor embedding (tSNE) (Fig. 5G) and a principal components analysis (PCA) plot (Fig. S7B), showing them to be well separated in multidimensional space. Because restricting our analysis to significantly regulated genes could bias our results, we reperformed our analysis using network propagation to embed the entirety of our quantitative data per gene within their pathway network neighborhoods, which resulted in highly similar results (Fig. S7C, see Methods).

### Host translation inhibitor plitidepsin provides pan-variant therapeutic efficacy *in vivo*

Viruses are known to hijack host translational machinery to preferentially produce viral proteins. Analysis of host pathway responses after VOC infection revealed the translation module to be the third most correlated with the AHR, alongside three other translation-related modules, representing a conserved host response across the viruses (Fig. 6A). Previously, we and others found inhibition of eukaryotic translation elongation factor 1A (eEF1A) with plitidepsin to exert potent antiviral activity against SARS-CoV-2 within the nanomolar concentration range (*42*). Plitidepsin has since entered phase III clinical trials for the treatment of COVID-19 (*43*). We also demonstrated plitidepsin to retain antiviral efficacy against all five VOCs in a cell culture model (*44*). Here, we assessed whether plitidepsin could maintain antiviral efficacy against all five VOCs *in vivo*. We intranasally infected K18-hACE2 mice with each SARS-CoV-2 VOC, or W1 control, and administered 0.3 mg/kg of plitidepsin via intraperitoneal injection once daily for three days (Fig. 6B). On the fourth day, viral titers were quantified from the lungs via TCID50. In line with cell culture results, we observed decreases in viral titers for all VOCs, all significant (p<0.05) except for Gamma, which showed a trend of reduction that did not reach significance (p=0.08; Fig. 6C). These results support plitidepsin as a therapeutic strategy to treat SARS-CoV-2 infection and highlight its potential as a broadly effective therapy against future SARS-CoV-2 variants.

**Figure 6.**
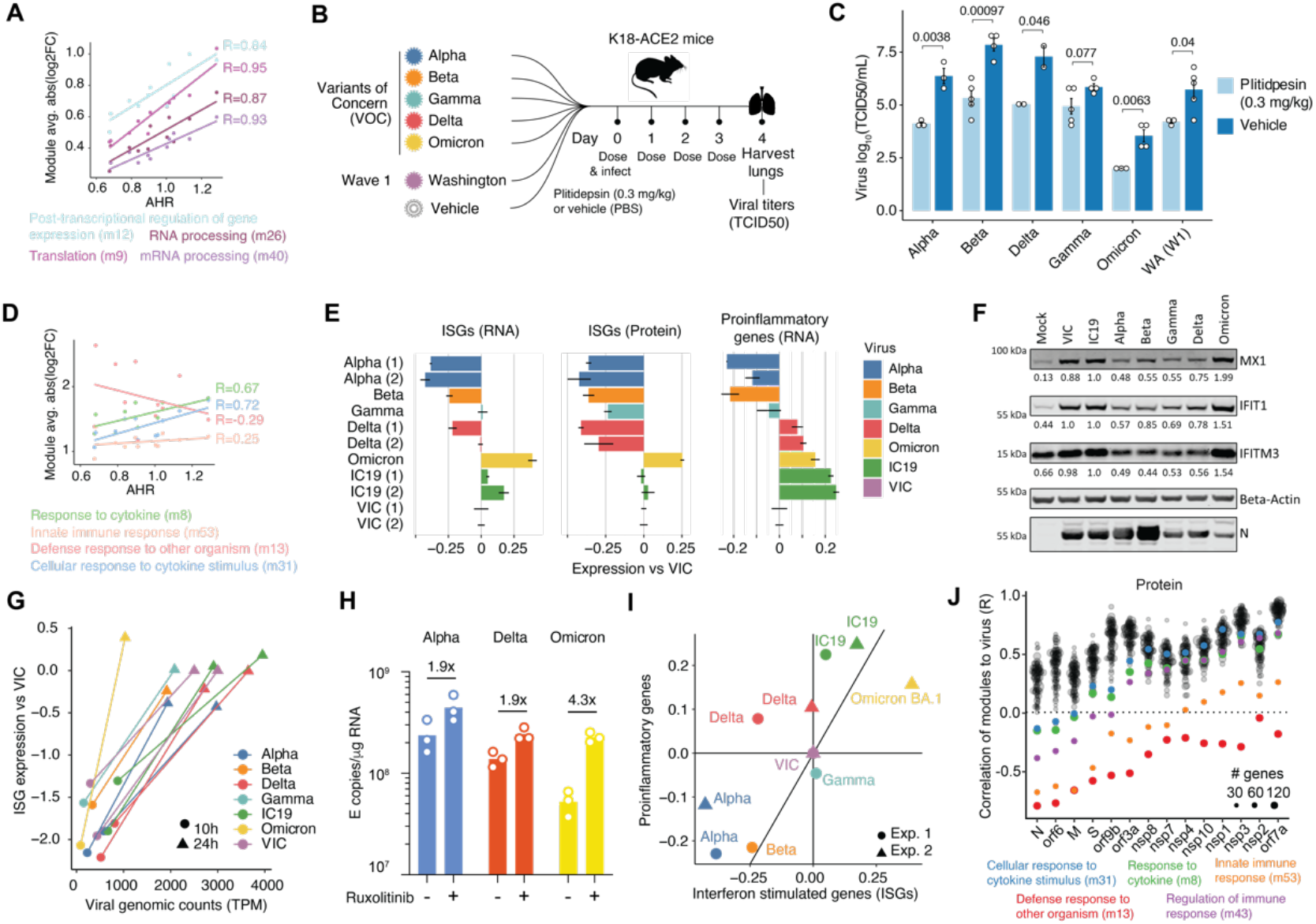
All variants depend on host translation but differentially regulate inflammatory responses. **(A)** Translation related modules within the top 10 most correlated with the AHR (from Fig. 5F). Scatter plot, linear regression lines, and Pearson’s R correlation coefficients are shown for each module from the RNA dataset. **(B)** Experimental workflow for *in vivo* pharmacological study. K19-ACE2 mice were treated with a VOC, wave 1 control (Washington strain or WA), or vehicle (PBS) on day 0 and dosed with 0.3mg/kg plitidepsin once a day for three days. Mouse lungs were harvested on the fourth day and viral titers were assessed by TCID50. Five mice were included per group. Mice with low infection were excluded from subsequent analyses. **(C)** Comparison of viral titers (log_10_ TCID50/mL) in K18-ACE2 mice from each virus, comparing plitidepsin treatment to vehicle. Statistical significance was determined using a two-tailed t-test, p values are annotated. Error bars represent the standard error (SE). **(D)** Innate immune and inflammation related modules within the 10 least correlated with the AHR, as in (A). **(E)** RNA and protein expression of interferon stimulated genes (ISGs) and RNA expression of pro-inflammatory genes at 24 hpi in Calu-3 cells. Expression is defined as the average log2FC of ISGs or proinflammatory genes (see Table S12 for list of genes) for each virus compared to VIC. Error bars depict the standard error (SE). Proinflammatory genes were sparsely detected at the protein level and thus excluded. **(F)** Western blot of MX1, IFIT1, IFITM3, and SARS-CoV-2 nucleocapsid (N) protein expression in Calu-3 cells infected with 2000 E copies/cell at 24 hours post infection. Protein quantification is noted and normalized to IC19 levels. **(G)** Correlation of viral genomic (leader + Orf1a) counts and ISG expression (versus VIC) as in (E) over time. Viral counts are represented as transcripts per million (TPM). Time points 10 and 24 hpi are represented by circles and triangles, respectively. **(H)** Viral replication of Alpha, Delta, or Omicron BA.1 in Calu-3 cells at 48 hpi in the presence or absence of ruxolitinib, a JAK/STAT inhibitor. Fold change between conditions is noted. **(I)** Relationship between ISG and proinflammatory gene expression for each virus at 24 hpi in Calu-3 cells relative to VIC for RNA as in (E). **(J)** Pearson’s R correlation between average log2FC for each module and levels of each viral protein across the viruses. Innate immune and inflammatory modules in the 10 least correlated category, including one additional innate immune term ranked 11th (m43, regulation of immune response), are colored. Viral proteins are ranked from left-to-right according to the average R values across the five inflammation-related terms.

### Variants differ in their modulation of the host inflammatory response

Our integrative computational analysis pinpointed innate immune and inflammatory processes as least correlated with the AHR (Fig. 6D, 5F), suggesting VOCs have evolved distinct relationships with the host antiviral inflammatory response, a phenomenon that is in line with the long-studied host-virus evolutionary arms race (*45*). To probe this further, we merged genes from the five inflammation modules and separated them into two groups: interferon-stimulated genes (ISGs) and proinflammatory genes (Table S12). We defined ISGs as directly induced upon interferon production after infection (containing an ISRE promoter) and pro-inflammatory genes as contributors to systemic inflammation which are driven by other transcription factors (i.e. NF-kB and AP-1), many of which have been associated with severe COVID-19 (*46*–*51*). We calculated a log_2_ fold change of VOCs versus VIC based on the average expression change for genes within each group to compare ISG and pro-inflammatory gene regulation across the VOCs.

We found all VOCs, except Omicron BA.1, uniformly antagonized ISG production at the protein level (Fig. 6E-F, S8A, S8C, S9). Alpha and Beta potently suppressed ISG responses at the RNA and protein level (Fig. 6E-F), which was also reflected in the level of IRF3 nuclear translocation (Fig. S9H). Gamma and Delta infection induced similar levels of ISG transcription relative to W1 viruses (Fig. 6E), but greater suppression was observed for particular genes, especially for Gamma (Fig. S8, S9). More robust suppression of ISG responses at the protein level, but not always at the RNA level, suggests the evolution of more effective and selective translational inhibition of host responses whilst leaving viral translation intact. Conversely, Omicron BA.1 triggered a greater type-I interferon response and accompanying ISG expression at the RNA and protein level compared to W1 (Fig. 6E-F, S8, S9), despite lower levels of replication compared to all other viruses (Fig. 6G). These observations were consistent using independent isolates of the same VOCs (Fig. S9D-G). Interestingly, we noticed greater rescue of Omicron replication, compared to Alpha and Delta, upon administration of the JAK/STAT signaling inhibitor ruxolitinib, underscoring how Omicron induces an antiviral interferon response that subsequently suppresses its replication (Fig. 6H) (*52*).

To further compare VOC immunomodulation, we plotted the averaged ISG vs pro-inflammatory gene induction signatures (mRNA) for each VOC, relative to VIC (Fig. 6I). Alpha and Beta were in the lower left quadrant, representing robust antagonism of both ISG and pro-inflammatory gene expression. By contrast, Gamma regulated inflammatory gene expression similar to W1 VIC. Interestingly, Delta shifted into the upper left quadrant, displaying increased induction of pro-inflammatory genes compared to other VOCs (Fig. 6I), despite similar replication kinetics (Fig. 6G). Strikingly, Omicron BA.1 did not cluster with any of the other variants due to increased induction of both ISGs and pro-inflammatory genes compared to other VOCs and W1 VIC (Fig. 6I).

To investigate if inflammatory pathway regulation by the VOCs was associated with viral protein expression, as we found previously for Alpha (*14*), we correlated viral protein and RNA levels with pathway module regulation from our integrative computational analysis and ranked viral genes according to their correlation with inflammatory response modules (modules 8, 13, 31, 43, and 53). Expression of N and Orf6, both well-studied innate immune antagonist proteins (*53*–*55*), were the most anticorrelated with inflammatory response modules, consistent with the notion that their enhanced expression contributes to the suppression of the host innate immune response across the VOCs (Fig. 6J, S7D). In conclusion, each virus evoked surprisingly different host inflammatory responses during infection; early VOCs Alpha and Beta tend to induce muted responses whereas later VOCs Delta and Omicron tend to drive greater inflammatory responses. These data suggest step-wise evolution of an increasingly sophisticated and more nuanced manipulation of host responses to promote VOC replication and transmission.

As mentioned earlier, Omicron BA.1 did not display enhanced innate immune antagonism compared to W1. This was despite a slightly higher expression of the innate antagonist, Orf9b (Fig. 2B-C, 2J). To probe Orf9b function, we generated a reverse genetics Orf9b deletion virus in the Alpha background (Fig. S8E) and observed it to trigger slightly enhanced innate immune responses during infection in Calu-3 cells relative to wild-type Alpha infection (Fig. S8F). This confirmed Orf9b contribution to innate immune suppression. By contrast, we have previously shown, using an Alpha Orf6 KO virus in Calu-3, that Orf6 is a potent innate immune antagonist (*52*). The fact that Omicron BA.1 down-regulates Orf6 expression and only slightly upregulates Orf9b, a weaker innate antagonist, suggests Omicron BA.1 may not have enhanced expression of the full complement of antagonists required for the most effective innate immune suppression. It is also possible that mutations in Omicron and Delta Orf9b affect their functional potency. Intriguingly, we found that the Alpha Orf9b KO virus had a defect in replication in primary HAEs, which was not rescued by inhibiting interferon signaling with ruxolitinib, suggesting that Orf9b may have additional roles in promoting viral replication (Fig. S8G).

### Omicron subvariants evolved innate immune antagonism by modulating Orf6

Phylogenetic studies indicate that the five VOCs (Alpha, Beta, Gamma, Delta, and Omicron BA.1) evolved independently from W1 virus. However, Omicron subvariants (BA.2, BA.4, and BA.5) are thought to have evolved from a common ancestor in a complex and incompletely understood way (*12, 56*). Omicron subvariants have co-circulated, becoming globally or locally dominant with evidence for adaptation to evade innate immune responses (*12, 52, 57*). To probe the molecular mechanisms underlying their differences, we globally quantified mRNA and protein levels in Calu-3 cells infected with equal doses of Omicron subvariants BA.1, BA.2, BA.4, or BA.5 alongside Alpha, Delta, and W1 IC19 (Fig. 7A). Assessment of viral replication by viral genome counts in RNAseq data demonstrated that in this experiment BA.1 and BA.5 replicated comparably, and more efficiently than BA.2 and BA.4 (Fig. 7B). Because Omicron subvariant replication was significantly reduced relative to prior VOCs (Fig. 7B), we compared them to each other and focused on the late (48 hpi) time point to better capture the viral and host response to infection.

**Figure 7.**
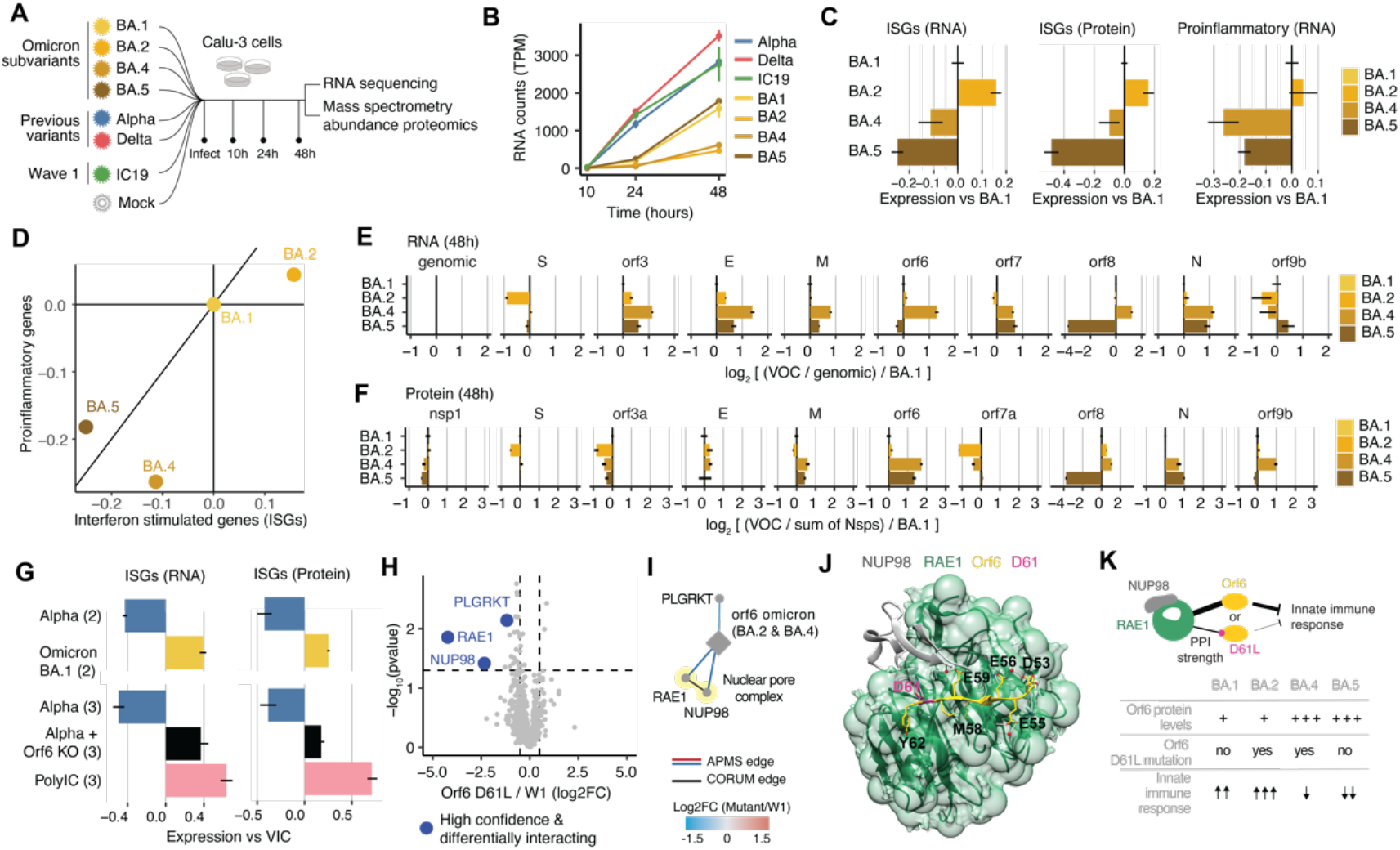
Omicron subvariants evolved innate immune antagonism by modulating Orf6. **(A)** Experimental workflow. Calu-3 cells were infected with Omicron BA.1, BA.2, BA.4, or BA.5 alongside Alpha, Delta, and W1 control IC19. Cells were harvested at 10, 24, and 48 hpi for bulk mRNA sequencing. Cells were harvested at 24 and 48 hpi for abundance mass spectrometry proteomics analysis. **(B)** Counts of viral genomic (leader + Orf1a) RNA over time for each virus. **(C)** ISG (RNA and protein) and proinflammatory gene (RNA) expression at 48 hpi relative to BA.1. Proinflammatory genes were sparsely detected at the protein level and thus excluded. Error bars represent the standard error (SE). **(D)** Relationship between ISG and proinflammatory gene expression for each virus at 48 hpi in Calu-3 cells relative to BA.1 for RNA as in (C). **(E)** Expression of viral RNA for Omicron subvariants, normalized to viral genomic (leader + orf1a) counts for each virus and set relative to BA.1. Error bars represent the standard error (SE). **(F)** Expression of viral protein for Omicron subvariants, normalized to the sum non-structural protein (Nsps) intensities for each virus and set relative to BA.1. Error bars represent the standard error (SE). **(G)** ISG RNA (left) and protein (right) expression versus VIC for Alpha (blue), Omicron BA.1 (yellow), an Orf6 knock-out version of Alpha created using reverse genetics (black), and poly I:C (pink). **(H)** Affinity purification mass spectrometry (APMS) of Orf6 D61L (occuring in BA.2 and BA.4 but not BA.1 or BA.5) compared to wave 1 (W1) Orf6 in HEK293T cells. All detected proteins are plotted with high confidence interactions that are also significantly differentially interacting (abs(log2FC)>0.5 & p<0.05) highlighted in blue. **(I)** Network representation of differentially interacting proteins from (H). Diamonds indicate the viral protein and circles indicate the human prey proteins. Colored edges represent the log2FC of human protein intensity between the mutant and W1 (log_2_(mutant/W1)). Known protein complexes are shaded yellow and connected by black edges from CORUM (nuclear pore) (*28*). **(J)** X-ray crystallography structure (PDB 7VPG) of RAE1 (green), NUP98 (gray), and Orf6 (yellow). The Orf6 D61 residue is indicated in pink and is shown to form a hydrogen bond (gray sticks) with RAE1. Other Orf6 residues (E59, E56, D53, and E55) that participate in interactions with RAE1 are also shown. M58 inserts into a RAE1 hydrophobic pocket. **(K)** Model of the effects of Orf6 levels and Orf6 D61L mutant status on the innate immune response. The nuclear pore (RAE1 and NUP98) physically interacts with Orf6, which suppresses the nuclear translocation of ISG-inducing transcription factors (e.g. STAT1) and export of ISG mRNAs during infection. This interaction is weakened, but not abolished, when the D61L mutation is present, resulting in diminished innate immune antagonism by Orf6. BA.1 and BA.2 downregulate Orf6 relative to early-lineage SARS-CoV-2 (IC19), which results in an increased innate immune response during infection, exacerbated by the presence of the D61L mutation in BA.2. Conversely, BA.4 and BA.5 upregulate Orf6 protein to similar levels; however, BA.5 more strongly antagonizes the innate immune response, which we speculate is due to the absence of the D61L mutation.

Strikingly, compared to BA.1, BA.2 stimulated a stronger innate immune response despite reduced replication. Host responses to BA.4 infection were decreased when compared to BA.1, particularly for pro-inflammatory gene induction (Fig. 7C-D). Thus, poor replication (BA.2 and BA.4) does not necessarily correlate with reduced host inflammatory responses, suggesting differences in the regulation of host innate immunity between these viruses. BA.5 displayed the greatest suppression of ISG and pro-inflammatory gene induction (Fig. 7C-D, S8B, S8D). Importantly, comparing host responses to Omicron subvariants with those to Alpha and Delta indicates that BA.5 is closest to Alpha (Fig. S10A-E), the best overall innate immune suppressor, suggesting that BA.5 could evolve further Alpha-like host adaptations in the future to even better antagonize innate immune activation.

Based on these data and our previous observations (*52*), we next hypothesized that Omicron subvariants evolved enhanced innate immune suppression by modulating the expression of specific viral proteins known to inhibit interferon responses. Interrogation of global viral RNA and protein expression during infection revealed differences in the production of structural and accessory proteins, but not non-structural proteins, across the Omicron subvariants (Fig. 7E-F, S10C-E). At the protein level, we observed decreased Orf3a and Orf7a expression in BA.2, increased Orf9b expression in BA.4, and a dramatic loss of Orf8 expression in BA.5. Interestingly, we observed an increase in Orf6 and N protein expression in BA.4 and BA.5, both known innate immune antagonists. Although Orf6 viral sgRNA was increased in BA.4, it was not for BA.5, suggesting the existence of a selective translational or post-translational control mechanism underlying Orf6 protein production for BA.5. Similarly, we previously observed increased viral sgRNA but decreased Orf6 protein production for BA.1 (Fig. 2B-C). Furthermore, our prior integrative computational analysis found N and Orf6 to be the most anticorrelated with the innate immune response during infection (Fig. 6J), underscoring their role in innate immune regulation. To validate the impact of Orf6 on the innate immune response, we used an Orf6 deletion virus in the Alpha background. The Orf6-deleted virus strongly induced ISG production compared to the Alpha control, reaching similar levels as Omicron BA.1 (Fig. 7G, S10F), consistent with the fact that BA.1 showed decreased Orf6 protein expression compared to the other VOCs (Fig. 2C). These results suggest that Orf6 is an important regulator of the innate immune response during SARS-CoV-2 infection and could underpin the evolved immunomodulatory profile of Omicron subvariants.

Given similar Orf6 protein levels, we were surprised that BA.4 and BA.5 induced different degrees of innate immune responses, with BA.5 seemingly suppressing ISG expression more strongly than BA.4 (Fig. 7C). However, BA.4 and BA.2 both contain a protein-coding mutation in Orf6 (D61L) and comparative APMS studies revealed the D61L mutation to strongly reduce the interaction between Orf6 and the mRNA transport factor RAE1 and nucleoporin NUP98 (Fig. 7H-I). Orf6 is known to bind to the nuclear pore to prevent the nuclear translocation of ISG-associated transcription factors (e.g. STAT1), as well as modulate the nuclear export of ISG mRNAs for cytoplasmic translation (*53, 58*). Thus, a decreased interaction between Orf6 and RAE1/NUP98 could explain the reduced suppression of ISGs seen for BA.4 and BA.2. Furthermore, using the RAE/NUP98/Orf6 crystal structure (PDB 7VPG) (*59*) and the SSIPe program (*60*), we predicted a model of the mutated complex and computed the difference of free energy change between mutant and W1 forms (ΔΔG = ΔG_bind,D61L_ - ΔG_bind,wt_). This analysis confirmed the D61L substitution unfavorably affects the RAE/Orf6 binding stability (ΔΔG = 1.2 kcal mol^−1^). Further interpretation of the structure revealed D61 to form a hydrogen bond with RAE1 R239. Other Orf6 residues, including E59, E56, D53, and E55, also formed similar interactions with RAE1 (Fig. 7J). In addition, M58 is inserted into a hydrophobic pocket within RAE1, providing a tight binding interface. Given the number of other residues regulating the Orf6/RAE1 interaction, we predict the D61L mutation to reduce, but not completely abolish, the interaction, in accordance with our APMS results (Fig. 7H-I). This mutation appears to also decrease the ability of Orf6 to inhibit the nuclear translocation of IRF3 and the export of host mRNAs (Fig. S8B, S8D; data shown in an accompanying paper).

In summary, we hypothesize the gain of the D61L mutation by BA.2 may be responsible for poor interferon antagonism and increased ISG levels compared to BA.1, in addition to low overall Orf6 protein levels. BA.4 upregulated Orf6 protein levels, which likely contributed to a boost in innate immune suppression, but due to the presence of the D61L mutation, it did not result in a potent inhibition of ISG expression. In contrast, BA.5 upregulates Orf6 and does not contain any Orf6 mutation, which likely explains why BA.5 is the strongest suppressor of innate immune activation among the Omicron subvariants (Fig. 7K).

## DISCUSSION

In this study, we sought to understand the biology and selective forces underlying the evolution of SARS-CoV-2 variants of concern. We applied an unbiased global approach to understand VOC infection biology and the impact of mutational adaptations on viral replication and host cellular responses. Although most work comparing VOCs has focused on Spike, we found that mutations beyond Spike critically impact VOC host-virus interactions.

Global comparison of viral RNA and protein expression across the viruses revealed VOCs tend to enhance expression levels of key innate immune antagonists to manipulate defensive host responses. Given VOCs are each independently derived from an early lineage wave 1 virus, our observation of different VOCs enhancing the same set of proteins (e.g. N, Orf6, and Orf9b), as well as evolution of N* expression (Fig. 2), strongly suggests convergent evolution of related strategies to subvert host defenses and improve transmission. Here, we were able to map several of the mutations responsible for altered protein expression, particularly for N, N* and Orf9b (Fig. 2), but the changes that regulate Orf6 remain elusive. We were also able to link adaptation of host-virus PPIs to specific viral mutations using APMS (Fig. 3), discovering an array of intriguing VOC-specific modifications to the host-virus protein-protein interaction landscape. For example, the novel VOC-specific protein N* interacts with the host RNAPII-associated factor complex (PAFc), suggesting VOCs have evolved to influence host gene expression through N*. Overall, our results highlight the plasticity of viral protein evolution and of host-virus interactions, showing that viral point mutations, including non-coding changes, can have significant functional impact.

One of the most striking features of SARS-CoV-2 VOCs is their sequential replacement of the single dominant VOC, with Alpha replacing wave 1, followed by Delta replacing Alpha, and then Omicron replacing Delta, rather than continuous global co-circulation. A wealth of data implicates Spike adaptation to escape adaptive antibody responses, but our data suggests that VOC evolution is more complex. Our integrative computational analysis revealed the most divergently regulated cellular pathways across the VOCs were those connected to innate immune and cytokine responses. This suggests that improving the capacity to regulate host innate immune responses makes a significant contribution to sequential VOC dominance and improved transmission. This also likely reflects a strong selection imposed by the human innate immune system on the virus, whose ancestor was originally adapted to evade bat innate immunity. However, our data do not explain why some VOC lineages in circulation, here Beta and Gamma, did not become dominant worldwide, but they do link viral genetics to host innate immune activation and its suppression. Nonetheless, we propose that a major force in shaping virus-host adaptations is related to evasion of innate and adaptive responses (*45*), as is consistent with our recent work demonstrating that evasion of innate immune responses defines pandemic HIV-1 (M) contributing to enhanced human-to-human transmission (*61*).

A significant event in SARS-CoV-2 evolution was represented by the emergence of the Omicron lineage. Omicron represents the biggest antigenic shift in SARS-CoV-2 evasion of the host adaptive immune responses, presumably driven by vaccination and/or prior infection, significantly reducing neutralization sensitivity. Omicron Spike mutations not only shifted antigenicity effectively to a new serotype, but also altered the biology of viral entry, reducing dependence on TMPRSS2 and surface fusion and instead moving towards dependence on cathepsin and endosomal fusion, and likely shifting *in vivo* tropism to the upper airway (*11, 62, 63*). Our data shows that Omicron is less effective in suppressing innate immune responses than Alpha or Delta. However, subsequent Omicron lineages, particularly BA.5, have enhanced their capacity to express key innate immune antagonist Orf6 and effectively suppress inflammatory responses during infection. We propose that having achieved the antigenic shift required to escape widespread adaptive immune responses to Spike, Omicron, like previous VOCs, subsequently experienced the next strongest selective pressure, resulting in the enhancement of innate immune evasion via upregulation of viral protein antagonists. A simple model of enhanced innate immune evasion by increased expression of Orf6 protein is somewhat confounded by the appearance of the Orf6 D61L mutation in the Omicron sublineages BA.2 and BA.4. However, APMS studies demonstrated this mutation to weaken the interaction with RAE1 and NUP98, suggesting a weakened inhibition of nuclear transport for mediators of the inflammatory response. Strikingly, D61L is encoded by the same 3 nucleotide substitutions in BA.2 and BA.4, suggesting this mutation occurred only once, perhaps in BA.2 and then BA.4 by recombination.

Surprisingly, we also detected differences in the expression of pro-inflammatory cytokines between cells infected with different VOCs. Cytokines including IL-6, CCL5, IL-8, TNF, and IL-1β have been associated with increased COVID-19 severity and poor prognosis (*46*–*51*). Whether such differences could account for VOC-specific pathogenicity, for example reports of more severe disease with Delta (*64*–*67*) or late Omicron sublineages (*68*–*72*), is hard to determine due to confounding pre-existing immunity. Thus, whilst VOCs, particularly Omicron with its modified Spike antigenicity, altered tropism, and enhanced innate immune evasion in later subvariants, may drive less severe disease than its predecessor Delta, which appears to be particularly inflammatory, dissecting the differences in the mechanisms driving pathogenicity and disease severity remains a necessary challenge. Although no model is perfect, a combination of cell lines, primary human cell models, and *in vivo* rodent models may produce the richest understanding of functional VOC evolution and its links to disease severity.

Importantly, our analysis pinpointed cellular pathways, including translation, cell cycle, and mRNA processing, that are similarly modulated across the VOCs during infection and thus represent putative targets for pan-coronavirus antivirals. For example, our previous work describing the SARS-CoV-2-human PPI network in conjunction with pharmacological testing uncovered the translation inhibitor plitidepsin to possess strong antiviral activity against both early SARS-CoV-2 strains *in vivo* and the VOCs *in vitro (29, 42, 44*). In the current study, we expanded upon these data by showing plitidepsin is also effective against VOCs *in vivo*. This demonstrates that targeting a host factor essential for viral replication, in this case eEF1A, provides a suitable strategy for the development of new antivirals.

Overall, our integrative systems multiomics platform enables the rapid evaluation of emerging viral mutations and their mechanistic consequences on viral replication and pathogenicity. Our analyses pinpoint conserved pathways that are central to the SARS-CoV-2 life cycle, paving the way to uncover broad antivirals for existing and future viral strains. Additionally, we uncover how viral evolution has led to divergent host responses, with implications for disease pathology, severity, and transmission in humans.

## SUPPLEMENTARY FIGURES

**Supplementary Figure 1.**
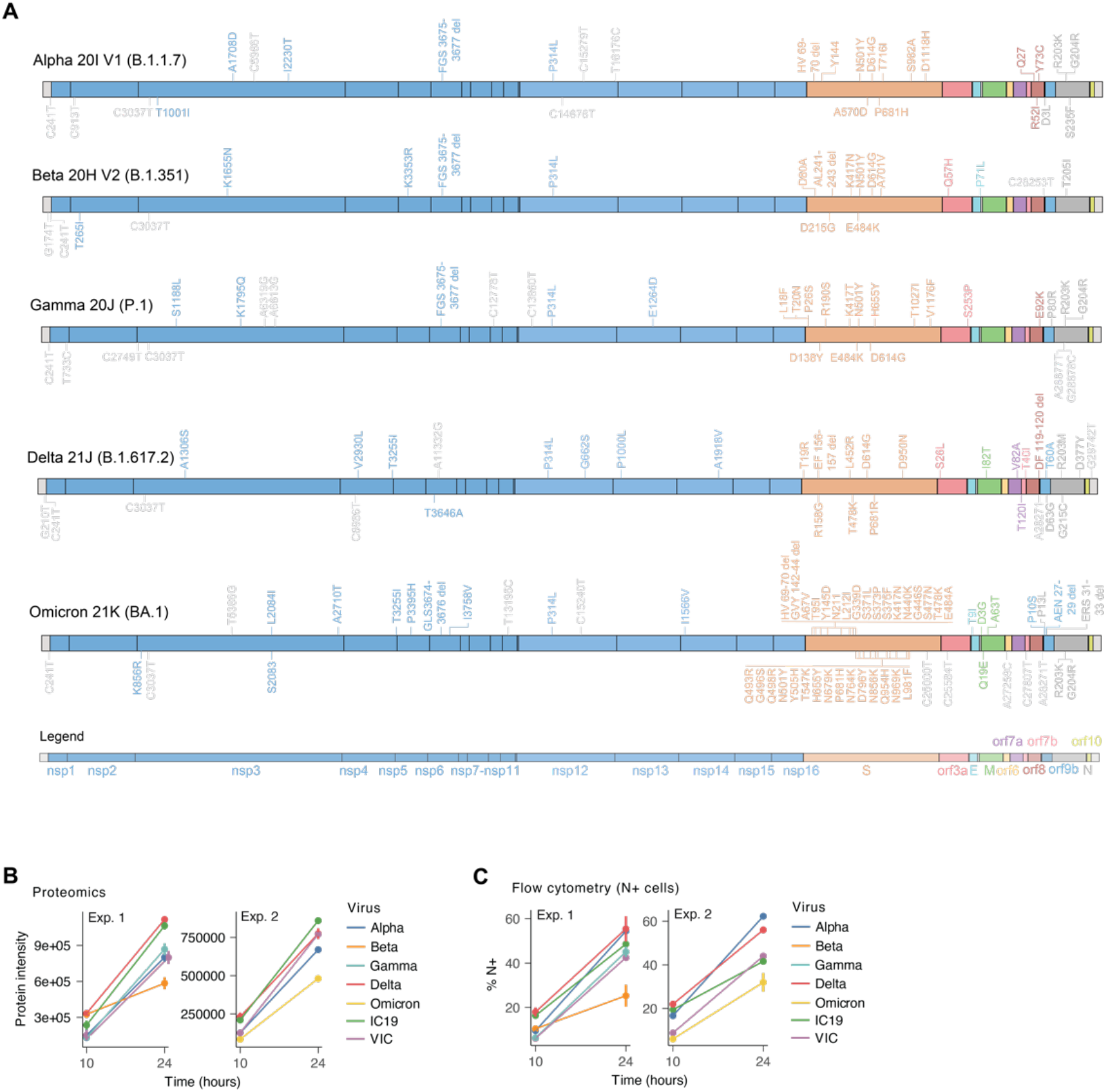
Genomic mutation map and replication kinetics of the SARS-CoV-2 variants of concern. **(A)** Annotated map of protein-coding (colored) and non-coding (light gray) mutations for each SARS-CoV-2 variant of concern. Color corresponds to the viral gene as depicted in the legend (bottom). **(B)** Quantification of the sum of non-structural protein intensities from abundance proteomics for each virus in each experiment (1 or 2) at 10 and 24 hours post infection in Calu-3 cells. Error bars depict the standard error (SE). **(C)** Flow cytometry assessing the percentage of cells positive for nucleocapsid (N) staining for each virus at 10 and 24 hours post infection in Calu-3 cells. Error bars represent the standard error (SE).

**Supplementary Figure 2.**
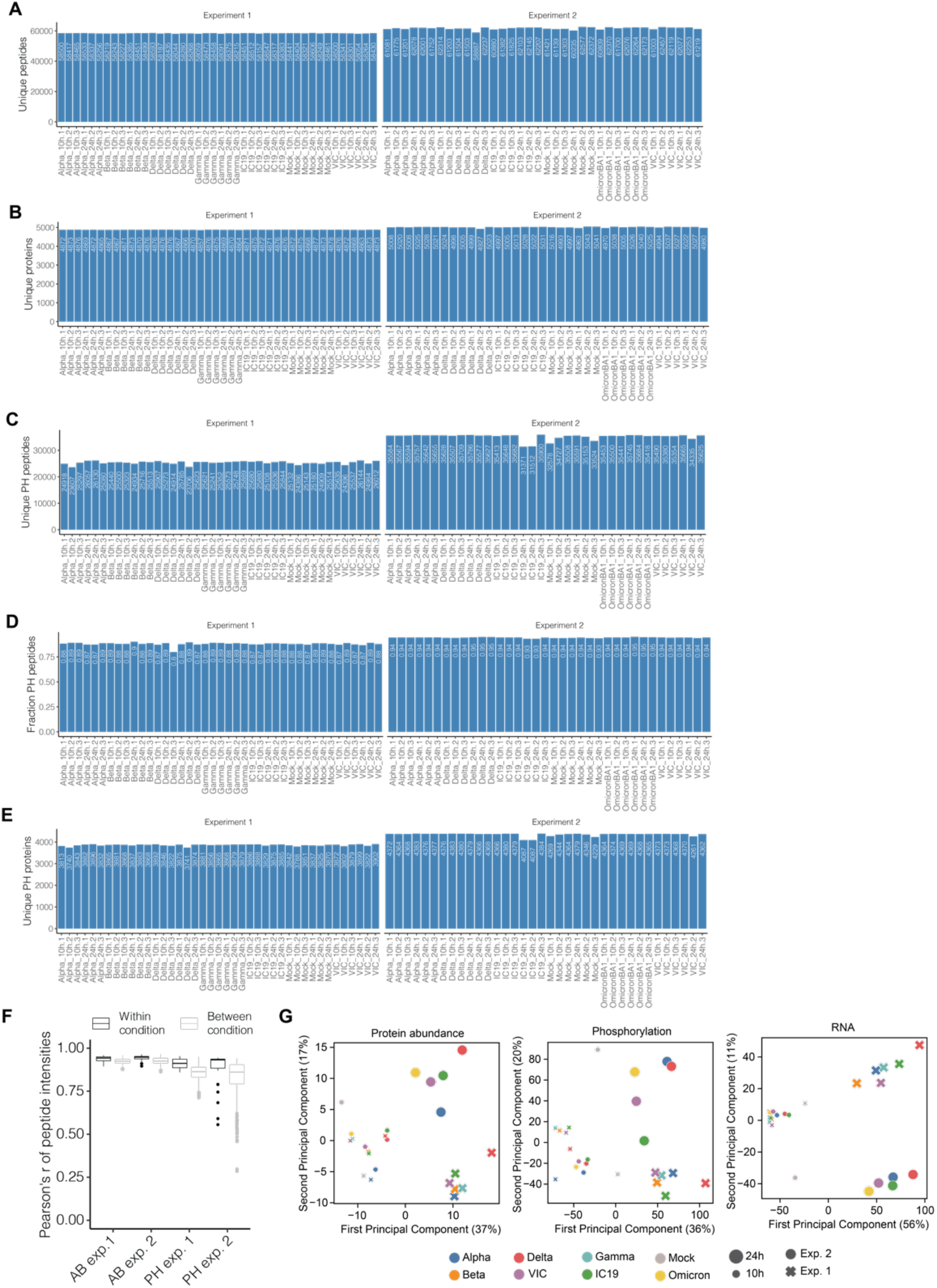
Proteomics detections and quality control. **(A)** Number of unique peptides (intensity > 2E5) detected in each sample from data independent acquisition (DIA) mass spectrometry abundance proteomics analysis of Calu-3 cells at 10 and 24 hours post infection for experiment 1 (left) and 2 (right). **(B)** Number of unique proteins mapping to peptides in (A). **(C)** Number of unique phosphorylated (PH) peptides (intensity > 2E5) detected in each sample from DIA mass spectrometry phosphoproteomics analysis of Calu-3 cells at 10 and 24 hours post infection for experiment 1 (left) and 2 (right). **(D)** Fraction of peptides with a phosphate modification in phospho-enriched samples, corresponding to peptides from (C). **(E)** Number of unique phosphorylated (PH) proteins mapping to peptides in (C). **(F)** Pairwise correlation of peptide intensities within condition (black) and between conditions (gray) for abundance proteomics (left) and phosphoproteomics (right) analysis in experiments 1 and 2. **(G)** Principal components analysis (PCA) of peptide intensities (protein abundance and phosphorylation, left and middle) or transcript counts (RNA, right) for virus infections in Calu-3 cells at 10 and 24 hours post infection. Experiment number is annotated by shape and time point by shape size. The percent of explained variance is indicated in parentheses for each principal component. Although the second principal component often separates according to the experiment, the rank ordering of the same viruses across different experiments is well preserved along the first principal component, which explains the greatest variance in the data.

**Supplementary Figure 3.**
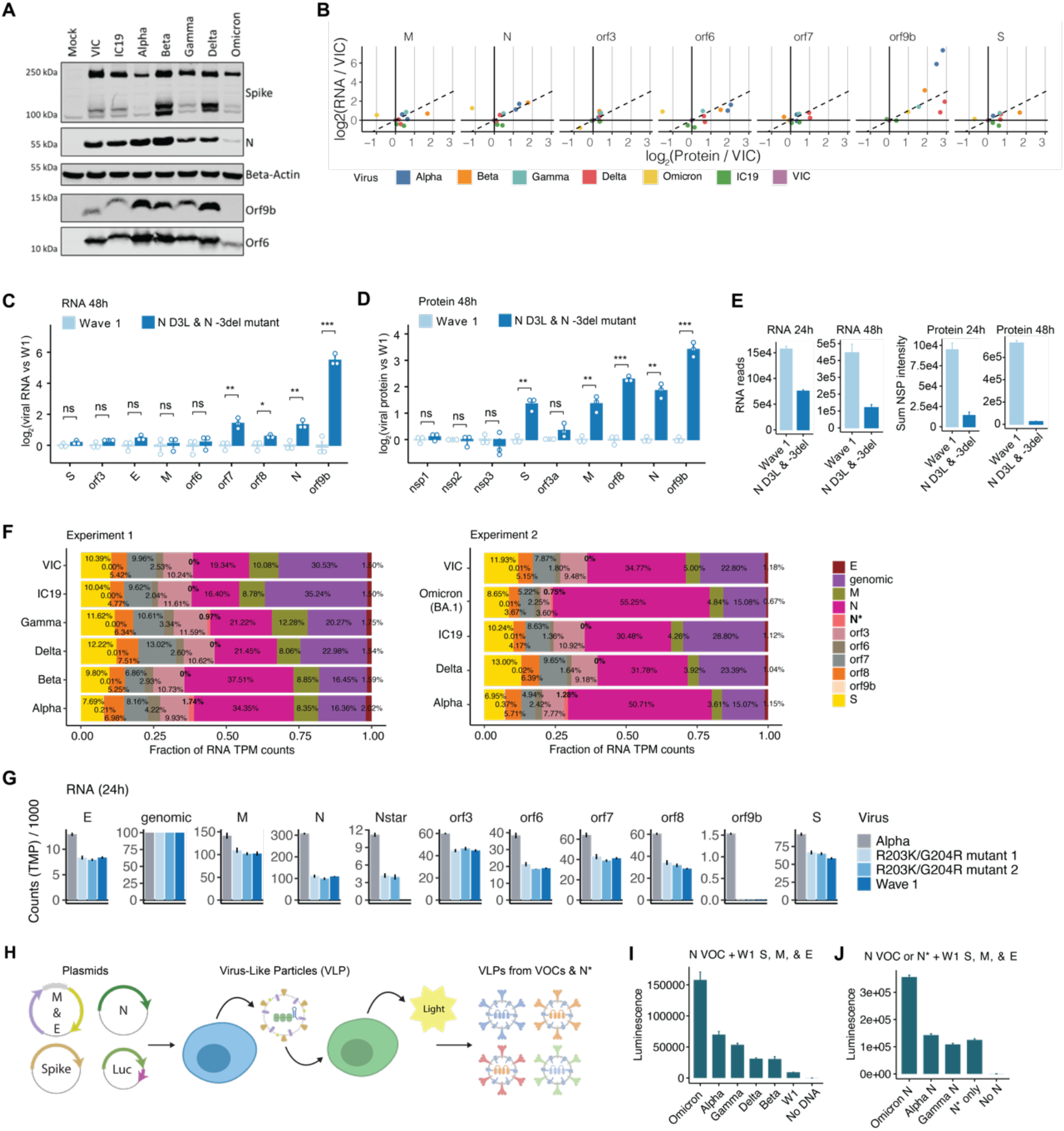
Viral RNA and protein expression, reverse genetics, and virus-like particles. **(A)** Western blot of Calu-3 cells infected with 2000 E copies per cell of VIC, IC19, Alpha, Beta, Gamma, Delta and Omicron BA.1 at 24 hours post infection (hpi). Of note, the lack of Orf9b detection in the Omicron infection samples may be due to mutations in Omicron Orf9b affecting the epitope for the antibody. **(B)** Correlation between viral RNA and protein expression for each virus relative to VIC, corresponding to Fig. 2D but separated by viral protein. Dashed line indicates x=y line. **(C)** Corresponding to Fig. 2H, expression of viral RNA from bulk mRNA sequencing at 48 hpi in A549-ACE2 cells infected with an wave 1 SARS-CoV-2 (W1) and one with two mutations, the D3L change in N (GAU→CUA) and a deletion at the -3 position upstream from the translation start site for N, created using reverse genetics approaches. Quantifications are normalized to genomic (leader + orf1a) counts per virus to control for differences in viral replication and defined relative to wildtype virus (log2(counts/genomic/W1)). Statistical significance testing is performed using a two-tailed t-test assuming unequal variance. *, p<0.05; **, p<0.01; ***, p<0.001. ns=not significant. **(D)** Corresponding to Fig. 2I, expression of viral proteins from abundance mass spectrometry proteomics at 48 hpi as in (C). Quantifications are normalized to the sum of non-structural protein intensities per virus to control for differences in viral replication and defined relative to W1 virus (log2(protein intensity/sum of nsp intensity/W1)). Statistical significance testing is performed as described in (C). **(E)** Baseline transcripts per million (TPM) RNA counts (left) and sum of non-structural proteins (right) per virus in at 24 and 48 hours post infection to reveal differences in viral replication. **(F)** Percentage of RNA reads (TPM) mapping to each SARS-CoV-2 viral gene. Genomic refers to reads mapping to Orf1a. Bold indicates N*. **(G)** Expression of viral RNA from bulk mRNA sequencing at 24 hpi for Alpha, a Wuhan early-lineage control, and two attempts at creating R203K/G204R mutants within the Wuhan background (mutant 1 and 2). All TMP counts are normalized to genomic (leader + orf1a) per virus to account for differences in viral replication. **(H)** Cartoon depiction of virus-like particle (VLP) system (*22*). Plasmids encoding SARS-CoV-2 structural proteins and luciferase are transfected into HEK293T, which produce virus particles containing the luciferase RNA as a genome. These VLPs were collected from the supernatant and administered to fresh HEK293T cells expressing ACE2 and TMPRSS2 from which the production of luciferase luminescence is quantified. **(I)** Results from VLP experiment described in (H). N proteins encoding mutations from each VOC were transfected alongside W1 forms of the other structural proteins (S, E, and M) and the resulting luminescence of VLP infection quantified. Error bars depict standard error (SE) of 4 biological replicates. **(J)** Results from VLP experiment described in (H). N proteins encoding mutations from Alpha, Gamma, and Omicron or N* were transfected alongside W1 forms of the other structural proteins (S, E, and M) and the resulting luminescence quantified. Interestingly, N* alone (no full length N) is able to package RNA and produce VLPs. All error bars in this figure represent the standard error (SE).

**Supplementary Figure 4.**
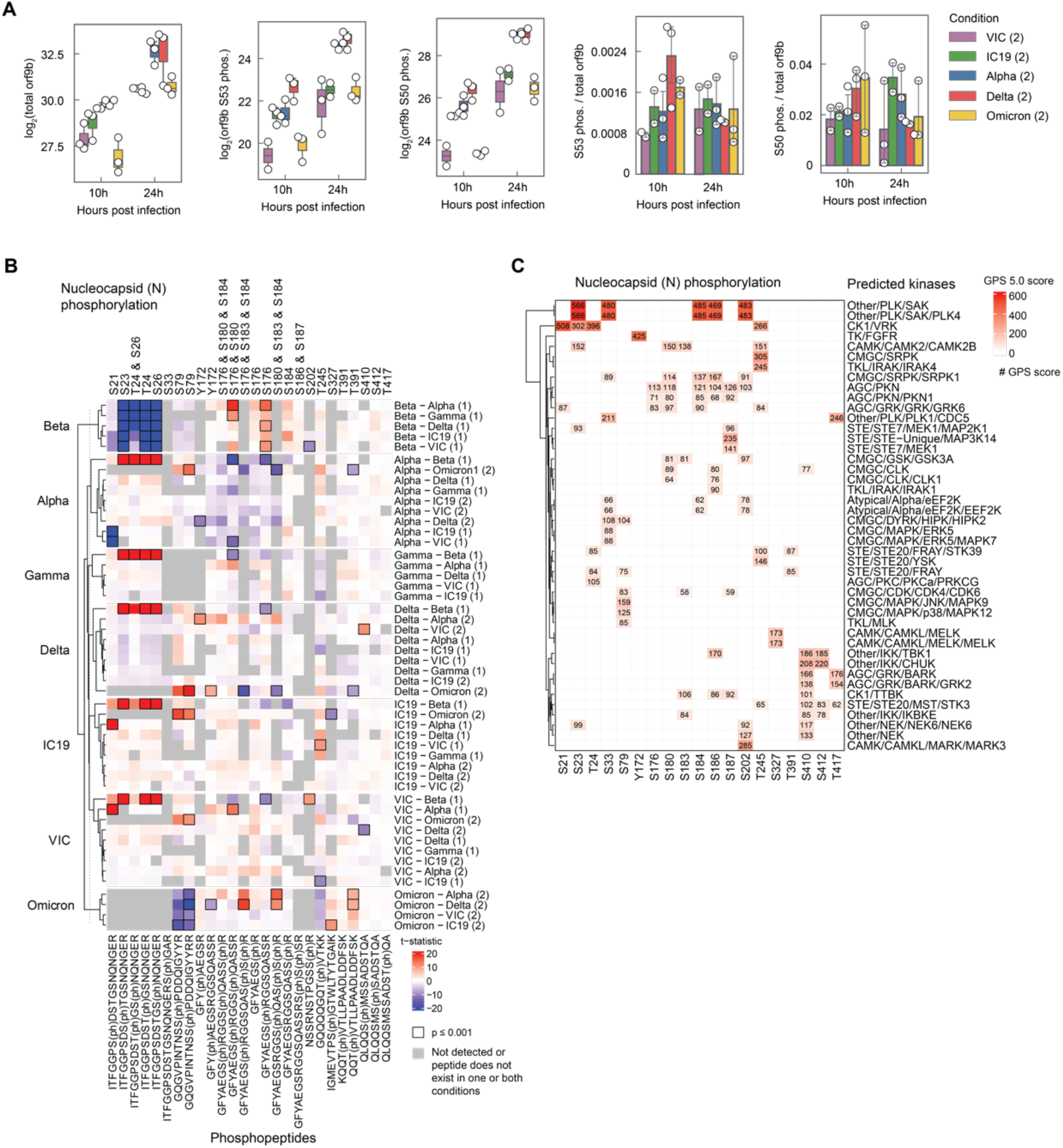
Phosphorylation of viral proteins during infection with SARS-CoV-2 variants. **(A)** Targeted proteomics (PRM) of total Orf9b abundance, phosphorylated S53 and S50 peptide abundances, and the ratio of phosphorylated Orf9b S53 and S50 to total Orf9b for experiment 2 (noted in parentheses). **(B)** Heatmap of significantly changed phosphorylated peptides mapping to N protein. Rows indicate the comparison (first virus is numerator, second is denominator) and columns indicate the phosphorylated peptides. Color represents the t-statistic and black bounding boxes indicate if p≤0.001. Gray boxes indicate peptides that were either not detected in one or both viruses or did not possess identical sequences between viruses. **(C)** Top kinases predicted to phosphorylate S, T, or Y residues on N protein using GPS 5.0 (*25*). To simplify the diagram, only kinases with at least one score greater than 80 were included as rows.

**Supplementary Figure 5.**
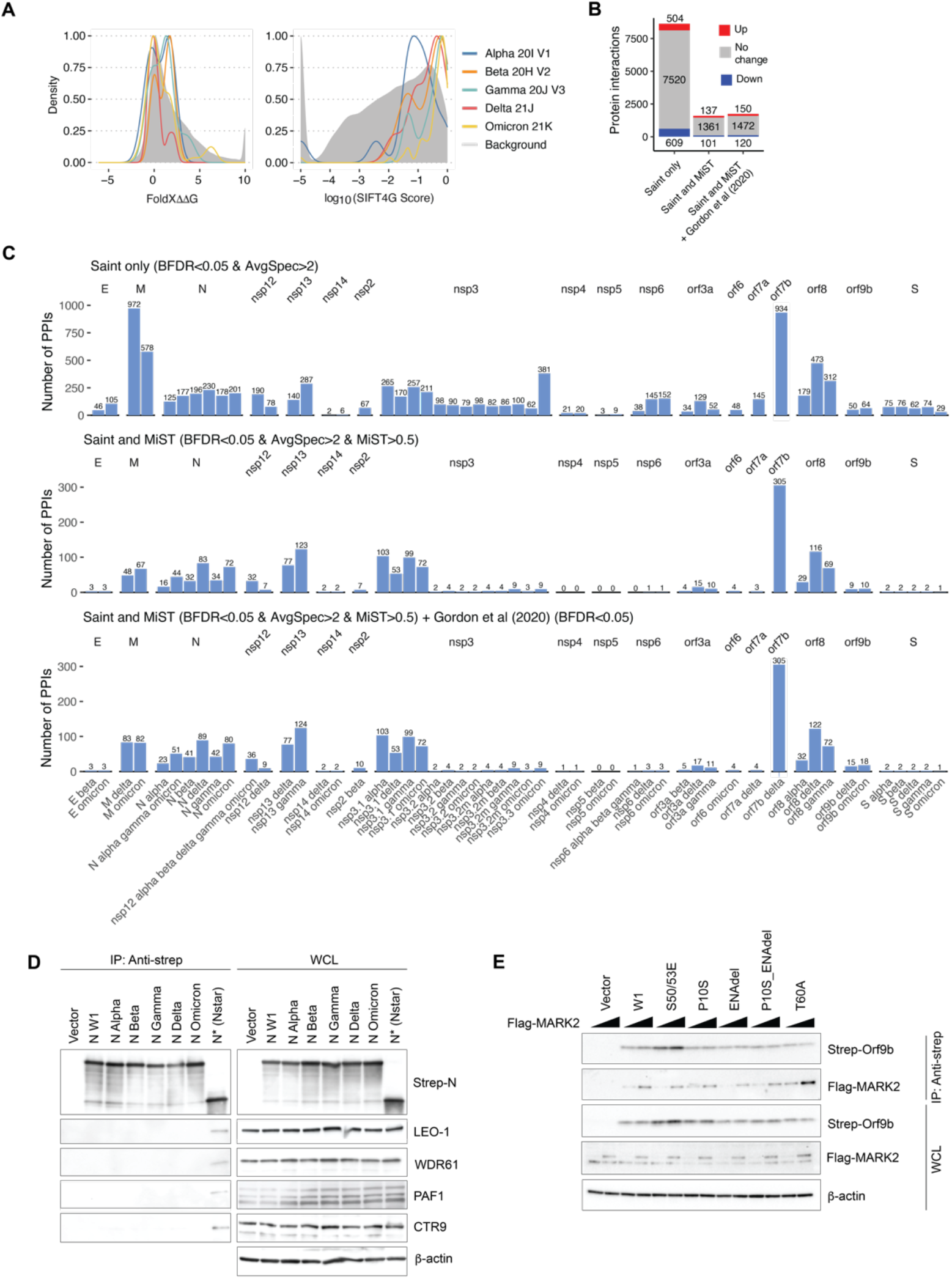
Structural analysis of mutations and virus-host protein-protein interactions. **(A)** Distribution of FoldX ΔΔG predictions (left) and evolutionary conservation SIFT4G scores (right) for mutations in SARS-CoV-2 variants of concern (VOCs) compared to the background distribution of all possible mutations. Interpretation is that VOC mutations are generally less structurally destabilizing and more evolutionarily tolerated than background. FoldX was run based on the highest quality structure available in the SARS-CoV-2 SWISS-model repository (*73*) containing 19 of the 28 viral proteins and SIFT4G was run against a clustered database of coronaviridae sequences. **(B)** Three scoring thresholds were generated, all included the requirement of average spectral counts (AvgSpec) to be greater than two: (i) SAINT only requiring SAINT BFDR<0.05, (ii) SAINT and MiST requiring BFDR<0.05 & MiST>0.5, and our final selected threshold (iii) SAINT, MiST, and Gordon et al. (2020) requiring [BFDR<0.05 & MiST>0.5] | [BFDR<0.05 & present in Gordon et al. (2020)]. These thresholds had to be satisfied in either the mutant or wave 1, but not both, for every mutant and wave 1 pair. The numbers of unchanging (i.e., conserved) interactions are indicated in gray, numbers of interactions increased in binding to the mutant in red, and numbers of interactions decreased in binding to mutant in blue. **(C)** The numbers of high-confidence interactions recovered for each bait under each scoring scheme described in (B). **(D)** Affinity purification western blot of strep-tagged VOC N or N*, blotting for endogenous PAF complex members: LEO1, WDR61, PAF1, and CTR9. This confirms APMS results showing N* specific interaction with the human PAF complex. **(E)** Affinity purification western blot of strep-tagged Orf9b (coding changes from Omicron (P10S and ENAdel) and Delta (T60A)) and flag-tagged MARK2 to confirm the enhanced interaction between Delta orf9b and MARK2 observed by APMS.

**Supplementary Figure 6.**
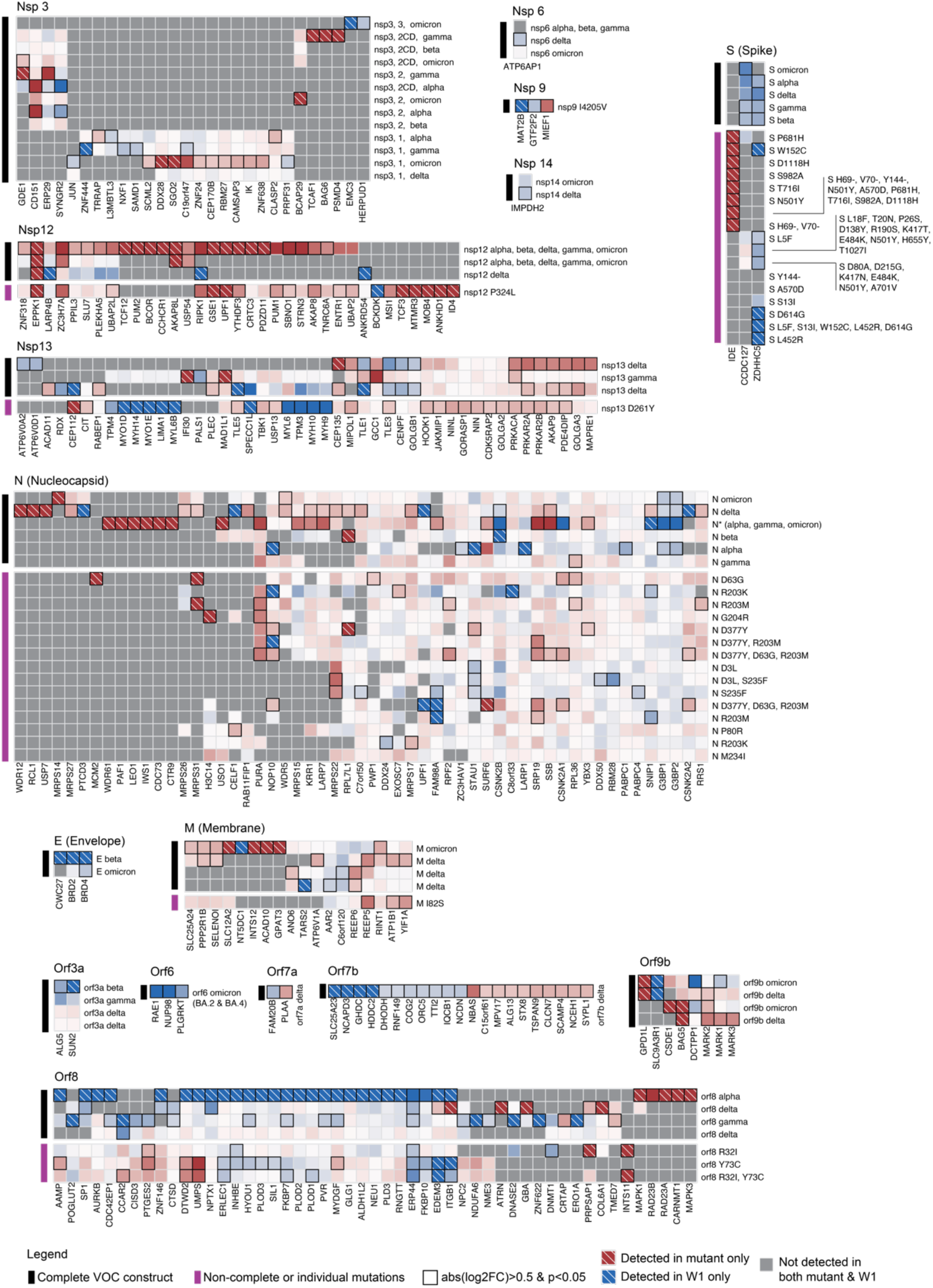
Heatmap of affinity purification mass spectrometry data. Each block represents a single annotated viral protein. Black lines denote viral bait proteins that are complete with all consensus mutations present in each variant of concern (“complete VOC construct”). If multiple mutations exist in a single VOC protein, all are combined together in the same construct. Purple lines depict other constructs possessing individual mutations or other combinations of mutations to assess the impact of single mutations on protein interactions (“non-complete or individual mutations”). Color indicates the log_2_ fold change of the abundance of each prey protein in the affinity purification between the mutant and W1 form. Red indicates increased binding to mutant and blue indicates decreased binding to mutant. Black bounding boxes indicate a significant differential interaction (p<0.05). White-dashed boxes indicate a prey detected in only the mutant (red) or W1 (blue). To simplify the diagram, only human prey proteins that were significantly differentially interacting (abs(log2FC)>0.5 & p<0.05) for one of the “complete variant constructs” are visualized for the remaining constructs. See Table S11 for the complete list of interactions.

**Supplementary Figure 7.**
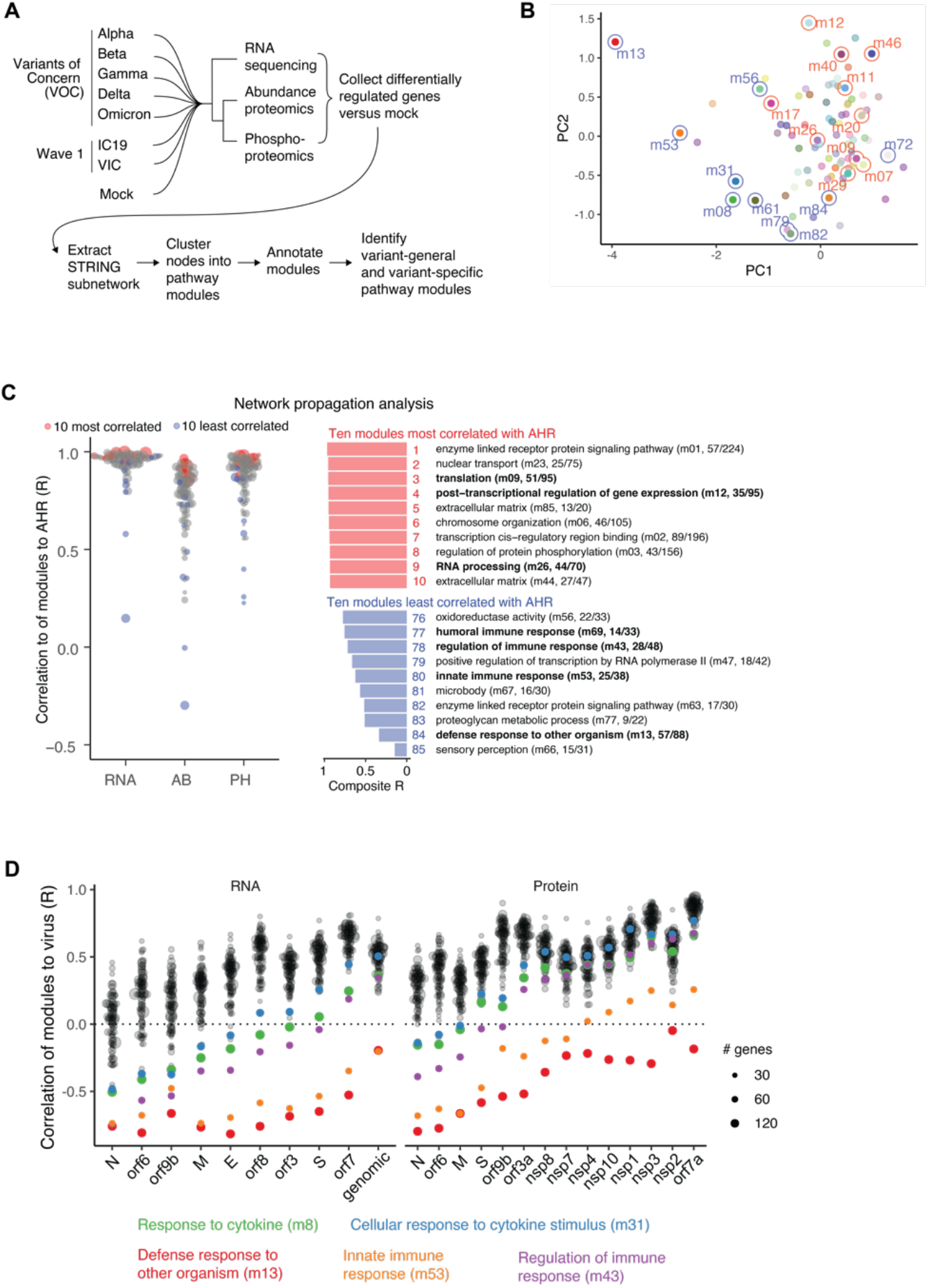
Integrative computational analysis and network propagation analysis of host response to infection. **(A)** Flowchart representation of integrative computational analysis depicted in Figure 5. Differentially regulated genes during VOC infection (versus mock) were extracted from the STRING network, clustered into 85 pathway modules based on network proximity, and annotated based on the most prevalent GO Biological Process term (see Methods). This analysis was used to identify conservation and divergence of host response to variants. **(B)** In Figure 5, we present a Pearson’s R correlation analysis to rank pathway modules. Here, we present a principal components analysis (PCA) of the underlying network proximity information for each module showing that the modules most (red) and least (blue) correlated with the AHR are separated in multidimensional space. This is intended to complement the tSNE plot presented in Figure 5G. **(C)** Network propagation analysis of multiomics datasets as done for integrative computational analysis to confirm findings using an orthogonal approach. **(D)** Pearson’s R correlation of viral RNA (left) and protein (right) levels and the Average Host Response (AHR) for each VOC, ranked by the average R across the annotated innate immune and inflammatory modules.

**Supplementary Figure 8.**
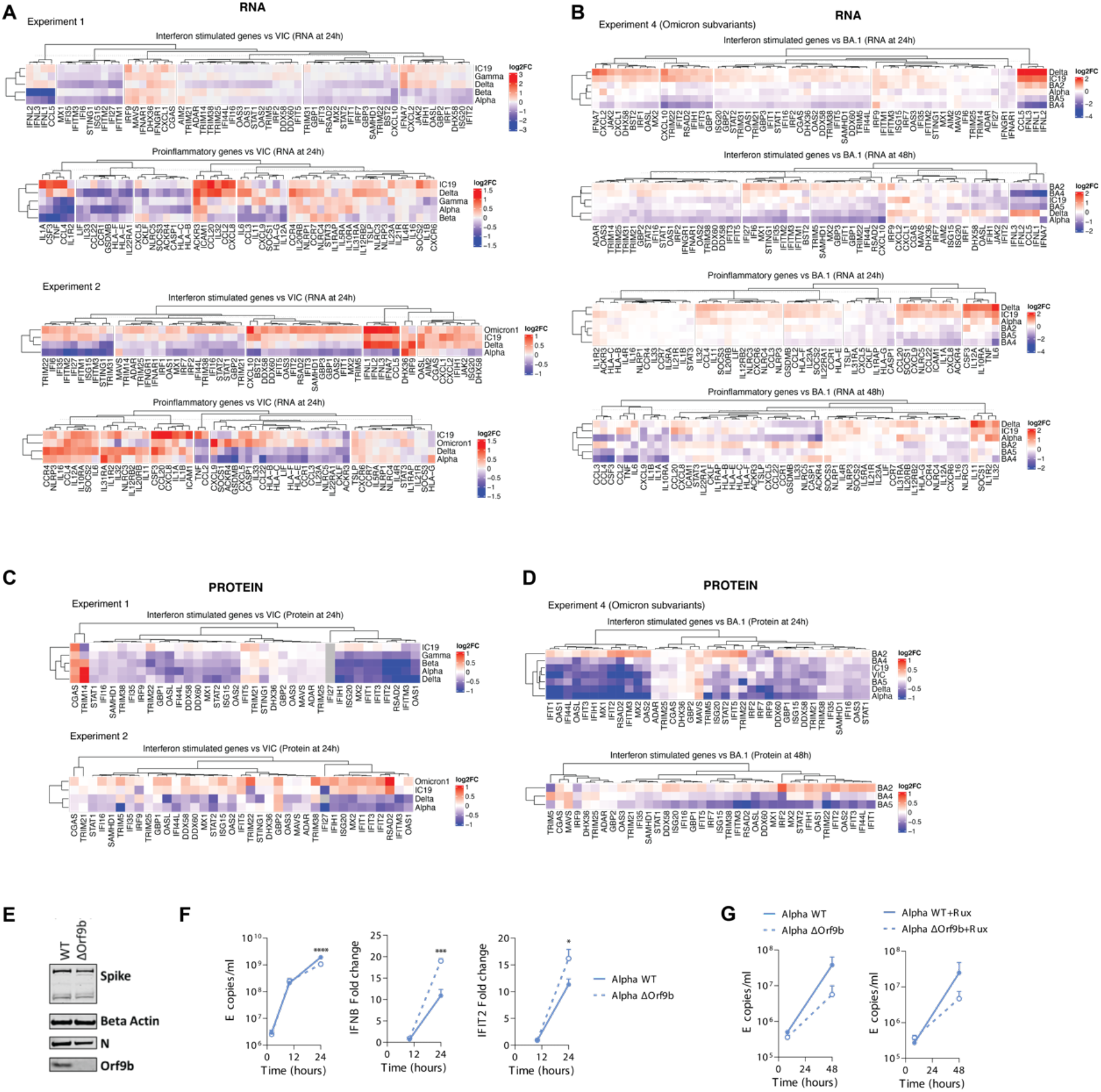
Heatmaps of inflammatory genes and Orf9b knock-out virus. **(A)** Heatmap of interferon stimulated genes (ISGs) and pro-inflammatory genes (RNA) for experiment 1 (top) and 2 (bottom). Color indicates the log_2_ fold change in expression relative to VIC. **(B)** Same as in (A) but for the Omicron subvariant experiment (RNA). **(C)** Same as in (A) but for protein. Only ISGs are included because pro-inflammatory genes were sparsely detected at the protein level, perhaps because they are typically excreted from the cell. **(D)** Same as in (B) but for protein. Only ISGs are included because pro-inflammatory genes were sparsely detected. **(E)** Western blot for Orf9b expression levels in Calu-3 cells infected with Alpha wild-type (WT) or Alpha Orf9b knock-out (ΔOrf9b) at 24hpi. **(F)** Replication and *IFNB* or *IFIT2* expression over time in Calu-3 cells infected with Alpha WT or Alpha ΔOrf9b (n=3). **(G)** Replication of Alpha WT or Alpha ΔOrf9b reverse genetics mutant virus in primary human airway epithelial (HAE) cells over time in the absence (left) or presence (right) of ruxolitinib (Rux) (n=3). For (F), pairwise statistical comparison at each timepoint was performed using One-Way ANOVA. p<0.05, *; p<0.001, ***; p<0.0001,****.

**Supplementary Figure 9.**
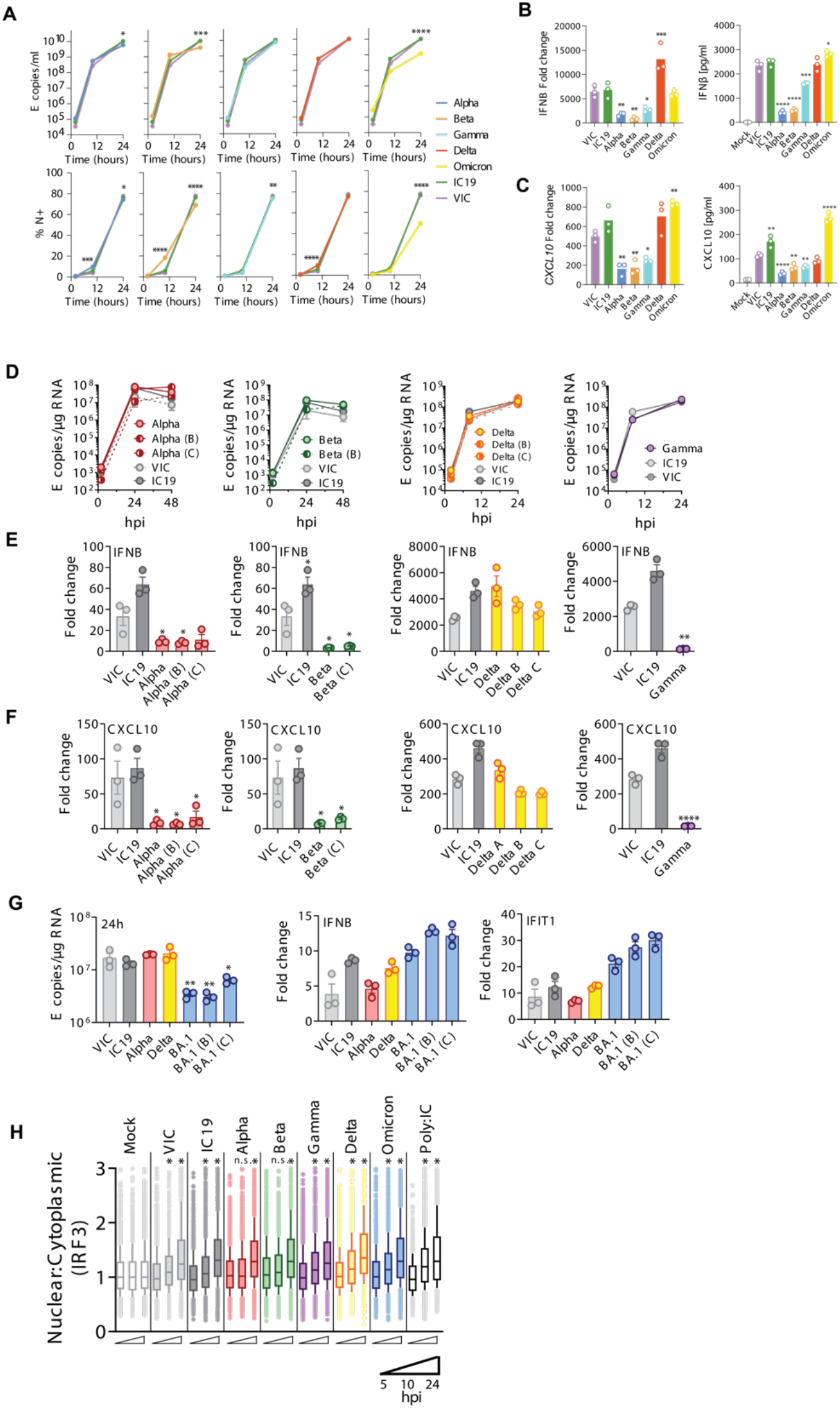
Targeted quantification of inflammatory response genes during VOC infection. (**A**) Replication of indicated isolates measured by E copies (top) or N-expression (bottom) in Calu-3 cells (n=3). (**B-C**) Expression of IFNβ (B) or CXCL10 (C) in infected Calu-3 cells at 24 hours post infection (hpi) for RNA (left) and protein (right) (n=3). Replication for this experiment is depicted in (A). (**D-F**) Infection of Calu-3 cells with 2000 E copies per cell while comparing multiple SARS-CoV-2 isolates of the same VOC. Replication over time (D), fold change of IFNB expression (E), and CXCL10 expression (F) are shown (n=3). **(G)** Viral replication and expression of *IFNB* or *IFIT1* of 3 independent Omicron BA.1 isolates in comparison to VIC, IC19, Alpha and Delta are shown at 24hpi (n=3). **(H)** Nuclear-to-cytoplasmic IRF3 ratio measured by single-cell immunofluorescence at 24 hours post infection in Calu-3 cells infected at 2000 E copies per cell. Per condition, 1500 randomly sampled cells are shown. For (A)-(G), pairwise comparisons against W1 viruses were performed using One-Way ANOVA. p<0.5, *; p<0.01, **; p<0.001, ***; p<0.0001, ****. For (H) pairwise comparisons against Mock at each time point were performed using Mann-Whitney test. Significance is indicated by *. ns=not significant.

**Supplementary Figure 10.**
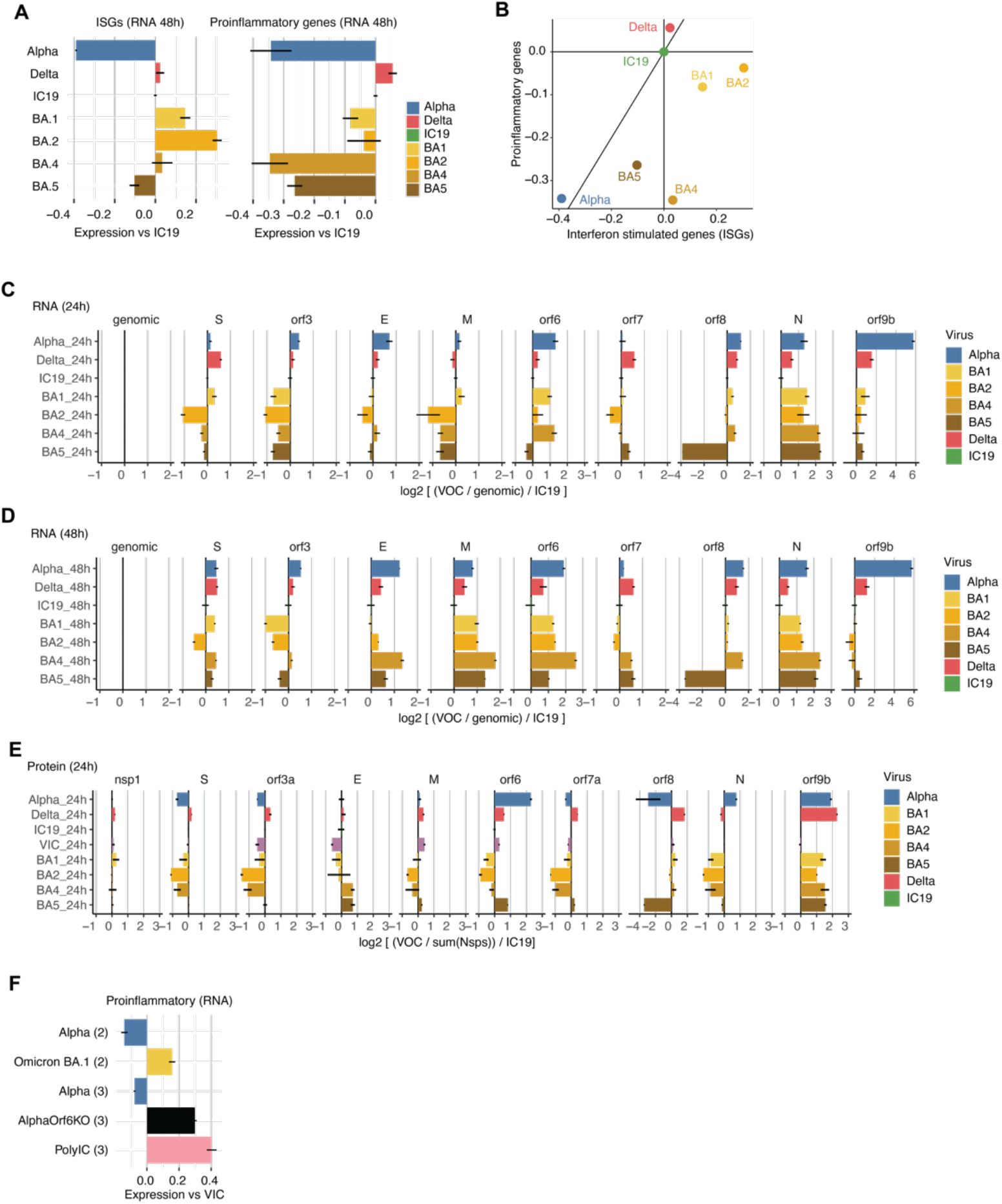
Inflammatory response and viral gene expression for Omicron subvariants and Orf6 knock-out virus. **(A)** RNA expression of interferon stimulated genes (ISG; left) and pro-inflammatory genes (right) at 48 hours post infection (hpi) for all viruses used in the Omicron subvariant experiment (Alpha, Delta, IC19, BA.1, BA.2, BA.4, and BA.5). Values are the average log_2_ fold change in expression relative to VIC. **(B)** Relationship between ISG and pro-inflammatory gene expression across all viruses listed in (A). **(C)** Viral RNA expression for viruses listed in (A) at 24 hpi. Quantity on the x-axis is the log_2_ of RNA counts normalized to genomic (Orf1a + leader) counts, then normalized to IC19. **(D)** Same as in (C) but at 48 hours post infection. **(E)** Same as in (C) but for viral protein. Quantity on the x-axis is the log_2_ of protein intensity normalized to the sum of non-structural protein intensities, then normalized to IC19. **(F)** Pro-inflammatory gene regulation by Alpha, Omicron BA.1, an Orf6 knock-out (KO) virus in the Alpha background generated using reverse genetics, and poly I:C, compared to VIC. All error bars in this figure represent the standard error (SE).

## SUPPLEMENTARY TABLES

Table S1. Consensus mutations in the VOCs from CoVariants.org.

Table S2. Quantification of host RNA transcript counts in transcripts per million (TPM) during infection.

Table S3. Differential host RNA expression analysis comparing conditions during infection.

Table S4. Quantification of host and viral proteins using abundance proteomics at the feature level.

Table S5. Quantification of host and viral proteins using abundance proteomics at the protein level.

Table S6. Differential host protein expression analysis comparing conditions during infection.

Table S7. Quantification of host and viral phosphorylated peptides using phosphoproteomics at the feature level.

Table S8. Quantification of host and viral phosphorylation sites using phosphoproteomics summarized at the site level.

Table S9. Differential host phosphorylation site intensity analysis comparing conditions during infection.

Table S10. Viral RNA, protein, and phosphorylation site abundances during infection, including analysis comparing viral phosphorylated peptide intensities across VOCs.

Table S11. Affinity purification mass spectrometry (APMS) metadata and results. Table S12. Integrative computational analysis metadata and results.

## ACKNOWLEDGEMENTS

We thank Wendy Barclay (Imperial College London, UK), and Alex Sigal and Khadija Khan (Africa Health Research Institute, Durban, South Africa) for the generous gifts of VOC isolates. SARS-CoV-2 cross-reactive antibody was a gift from Laura McCoy (University College London, UK), and Caco-2 cells were a gift from D. Bailey. We thank Dr. Randy Albrecht for support with the BSL3 facility and procedures at the ISMMS as well as Richard Cadagan for excellent technical assistance. We thank Tom Peacock at Imperial College London for helpful discussions.

## FUNDING

This work has been funded by:

National Institutes of Health grant U19AI171110 (NJK)

National Institutes of Health grant U19AI135990 (NJK)

National Institutes of Health grant U19AI135972 (NJK, AG-S)

National Institutes of Health grant R21AI159666 (JAD)

National Institutes of Health grant K99AI163868 (MB)

Defense Advanced Research Projects Agency (DARPA) Cooperative Agreement #HR0011-19-2-0020 (NJK, AG-S). The views, opinions, and/or findings contained in this material are those of the authors and should not be interpreted as representing the official views or policies of the Department of Defense or the U.S. Government.

Laboratory for Genomics Research (LGR) Excellence in Research Award #133122P (NJK)

F. Hoffmann-La Roche (NJK)

Vir Biotechnology (NJK)

Gifts from QCRG philanthropic donors (NJK)

Pharmamar (AG-S, KW)

This work was also partly supported by CRIPT (Center for Research on Influenza Pathogenesis and Transmission), a NIAID funded Center of Excellence for Influenza Research and Response (CEIRR, contract # 75N93021C00014) and by NIAID grant U19AI142733, NCI Seronet grant U54CA260560, the JPB and OPP foundations and an anonymous philanthropic donor to AG-S. Department of Defense W81XWH2110103 (LMS)

Department of Defense W81XWH2110095 (LMS)

National Institutes of Health grant 1R43AI165089-01 (LMS)

National Institutes of Health grant 1R01AI161363-01 (LMS)

National Institutes of Health grant 1R01AI161175-01A1 (LMS)

National Institutes of Health contract number 75N93021C00014 (LMS)

San Antonio Partnership for Precision Therapeutics (LMS)

San Antonio Medical Foundation (LMS)

Marie Skłodowska-Curie Individual Fellowships no. 896014. (RR)

European Research Council (ERC-Stg no. 639429) (PB)

The Rosetrees Trust (M362-F1; M553) (PB) NIHR GOSH BRC (PB)

CF Trust (SRC006; SRC020) (PB)

Wellcome Investigator Award 223065 (CJ)

Wellcome Investigator Award 220863 and Wellcome Collaborative Award 214344(GJT) MRC/UKRI G2P-UK National Virology consortium (MR/W005611/1) (GJT, CJ)

## AUTHOR CONTRIBUTIONS

Conceptualization: MB, AKR, BJP, MU, LGT, AG-S, CJ, LZA, GJT, NJK

Investigation: MB, BJP, AKR, MU, LGT, CY, RRR, AP, JB, GMJ, JX, JM, YZ, BH, ES, AR, RR, MW, WF, GDL, AP, VC, AMS, AC, ND, DP, GD, DM, JT, ALR, AD, XZ, RMK, ME, NL, TK, AC, MR, NM, SA, AH, HF, KO, JMF, MS, RH, IJ, MK, IE, KV, PB, RS, JAD, LMS, AP, MP, LM, KW, DLS, AG-S, CJ, LZA, GJT, NJK

Visualization: MB, AKR, BJP, MU, LGT, AG-S, CJ, LZA, GJT, NJK

Funding acquisition: AG-S, CJ, LZA, GJT, NJK

Supervision: KV, PB, RS, JAD, LMS, AP, MP, LM, KW, DLS, AG-S, CJ, LZA, GJT, NJK Writing: MB, AKR, BJP, MU, LGT, AG-S, CJ, LZA, GJT, NJK

Resources: MS

## DATA AVAILABILITY

Further information and requests for resources and reagents should be directed to and will be fulfilled by NJK. The mass spectrometry proteomics infection proteomics and affinity purification mass spectrometry (APMS) data have been deposited to the ProteomeXchange Consortium (http://proteomecentral.proteomexchange.org) via the PRIDE partner repository (*74*) with the dataset identifiers PXD036968 ( and password OoQuRtk5) and PXD036798 (reviewer username and password feVx3D2k), respectively. All mRNA sequencing data have been deposited in NCBI’s Gene Expression Omnibus (*75*) and are accessible through GEO Series accession number GSE213759 (reviewer token cxshmmmaxpqtxuj). All other data are available in the main text or the supplementary materials.

## METHODSs

### Description of proteomics and transcriptomics experiments

Global proteomics and transcriptomics studies were collected across seven (7) separate experiments (see Supplemental Tables). Experiment 1 contains Alpha, Beta, Gamma, Delta, IC19, VIC, and mock infections in Calu-3 cells harvested at 10 and 24 hpi and was processed for mRNA sequencing, abundance proteomics, and phosphoproteomics. Experiment 2 contains Alpha, Delta, Omicron BA.1, IC19, VIC, and mock infections in Calu-3 cells harvested at 10 and 24 hpi and was processed for mRNA sequencing, abundance proteomics, and phosphoproteomics. Experiment 3 contains Alpha Orf6 KO, Alpha wild-type, IC19, VIC, mock infections and polyIC treatment in Calu-3 cells harvested at 10 and 24 hpi and was processed for mRNA sequencing, abundance proteomics, and phosphoproteomics. Experiment 4 contains Omicron BA.1, BA.2, BA.4, BA.5, Alpha, Delta, IC19, VIC, and mock infection in Calu-3 cells harvested at 24 hpi and Omicron BA.1, BA.2, BA.4, and BA.5 in Calu-3 cells harvested at 48hpi and processed for abundance proteomics only. Experiment 5 contains Omicron BA.1, BA.2, BA.4, BA.5, Alpha, Delta, IC19, VIC, and mock infection in Calu-3 cells harvested at 24 and 48 hpi and processed for mRNA sequencing only. Experiment 6 contains viral RNA and protein counts from global mRNA sequencing and abundance proteomics, respectively, for Washington wild-type and N D3L + -3 deletion mutant virus in Washington background infection in A549-ACE2 cells at 24 and 48hpi. Experiment 7 contains viral RNA counts from global mRNA sequencing for R203K/G204R mutant virus in Wuhan virus background infection in Calu-3 cells at 24hpi. The purpose of experiment 7 was to assess the expression of N*.

### Infections in Calu-3 cells with SARS-CoV-2 variants

#### Cell culture

Calu-3 cells were purchased from ATCC (HTB-55) or AddexBio (C0016001) and Caco-2 cells were a gift from D. Bailey. Cells were cultured in Dulbecco’s modified Eagle’s medium (DMEM) supplemented with 10% heat-inactivated FBS (Labtech) and 100 U ml−1 penicillin–streptomycin, with the addition of 1% sodium pyruvate (Gibco) and 1% Glutamax. All cells were passaged at 80% confluence and were frequently monitored for mycoplasma contamination. For infections, adherent cells were trypsinized, washed once in fresh medium and passed through a 70-µm cell strainer before seeding at 0.2 × 10^6^ cells per ml into tissue-culture plates. Calu-3 cells were grown to 60–80% confluence before infection as described previously (*14*). For stimulation with poly:IC (Peprotech), 250ng of poly:IC were transfected using lipofectamine 2000 (ThermoFisher).

#### Viruses and reverse genetics

SARS-CoV-2 lineages Alpha (B.1.1.7) (*14, 15*), Beta (B.1.351), Gamma (P.1), Delta (B.1.617.2) (*11*) and Omicron (lineage 375 B.1.1.529.1/BA.1 and lineage B.1.1.529.2/BA.2) isolates were a gift from Wendy Barclay (Imperial 376 College London, UK). Beta (B.1.351), Omicron BA.4 (lineage B.1.1.529.4) and BA.5 (lineage B.1.1.529.5) were a gift from Alex Sigal and Khadija Khan (Africa Health Research Institute, Durban, South Africa) (*12*). Early-lineage isolate VIC was provided by NISBC and IC19 a gift from Wendy Barclay (*14*). Alpha Orf6 deletion virus (Alpha ΔOrf6) was achieved by mutation of the first two methionines: M1L (A27216T) and M19L (A27200T). Alpha Orf9b deletion virus (Alpha ΔOrf9b) was achieved by introducing the following synonymous mutations into N: I15I (T28318A) and N29N (T28360C), resulting in Orf9b L12Stop and M26T. Wuhan-Hu-1-D614G with R203K/G204R was achieved by introducing mutations R203K (G28881A, G28882A) and G204R (G28883C). Reverse genetics (RG) derived viruses were generated essentially as previously described (*76, 77*). Rescued RG SARS-CoV-2 viruses were sequenced using Oxford Nanopore as previously described (*78*). Viruses were propagated by infecting Caco-2 cells in DMEM culture medium supplemented with 10% FBS and 100U/ml penicillin/streptomycin at 37 °C as previously described (*14, 15*). Virus stocks used in experiments comparing Omicron subvariants, virus stocks were prepared in DMEM culture medium supplemented with 1% FBS and 100U/ml penicillin/streptomycin, which was maintained in Calu-3 infections. Virus was collected at 72 hpi and clarified by centrifugation at 2,100xg for 15 min at 4°C to remove any cellular debris. Virus stocks were aliquoted and stored at −80 °C. Virus stocks were quantified by extracting RNA from 100 µl of supernatant with 1 µg/ml carrier RNA using Qiagen RNeasy clean-up protocol, before measuring viral E RNA copies per ml by RT-qPCR (*14, 15*).

#### Infections

For infections, inoculi were calculated using E copies per cell as indicated. Cells were inoculated with diluted virus stocks for 2 h at 37 °C, subsequently washed once with PBS and fresh culture medium was added. At the indicated time points, cells were collected for analysis as previously described (*14*).

#### RT-qPCR of host gene expression in infected cells

cDNA was synthesized from RNA using SuperScript IV (Thermo) with random hexamer primers (Thermo). RT-qPCR was performed using Fast SYBR Green Master Mix (Thermo) for host gene expression and subgenomic RNA expression or TaqMan Master mix (Thermo Fisher Scientific) for viral RNA quantification. Reactions were performed on the QuantStudio 5 Real-Time PCR systems (Thermo Fisher Scientific). Viral E RNA copies were determined as described previously (*15*). Host gene expression was determined using the 2−ΔΔCt method and normalized to GAPDH expression. The following probes and primers were used: GAPDH fw: 5’-ACATCGCTCAGACACCATG-3’, rv: 5’-TGTAGTTGAGGTCAATGAAGGG-3’; IFNB fw: 5’-GCTTGGATTCCTACAAAGAAGCA-3’, rv: 5’-ATAGATGGTCAATGCGGCGTC-3’; CXCL10 fw: 5’-TGGCATTCAAGGAGTACCTC-3’, rv: 5’-TTGTAGCAATGATCTCAACACG-3’; IFIT1 fw: 5’-CCTCCTTGGGTTCGTCTACA-3’, rv: 5’-GGCTGATATCTGGGTGCCTA-3’, IFIT2 fw: 5′-CAGCTGAGAATTGCACTGCAA-3′, rv: 5′-CGTAGGCTGCTCTCCAAGGA-3′.

#### Detection of secreted cytokines by ELISA

Cytokine release by infected Calu-3 cells was measured in culture supernatants at 48 hpi. IFNβ and CXCL10 were measured using Human IFN-beta Quantikine ELISA Kit or Human CXCL10/IP-10 DuoSet ELISA reagents (biotechne R&D systems) according to the manufacturer’s instructions.

#### Western blotting

For western blotting, whole-cell lysates were separated by SDS-PAGE, transferred onto nitrocellulose and blocked in PBS with 0.05% Tween 20 (v/v) and 5% skimmed milk (w/v). For detection of Spike, N, Orf6, Orf9b, spike, MX1, IFIT1 and β-actin expression membranes were probed with rabbit-anti-IFIT1 (CST, #14769, clone D2×9Z), rabbit-anti-MX1 (CST, #37849, clone D3W7I) rabbit-anti-SARS spike (Invitrogen, PA1-411-1165), rabbit-anti-Orf6 (Abnova, PAB31757), rabbit-anti-Orf9b (ProSci, 9191), Cr3009 SARS-CoV cross-reactive human-anti-N antibody (a gift from Laura McCoy, UCL) and rabbit-anti-beta-actin (SIGMA), followed by IRDye 800CW or 680RD secondary antibodies (Abcam, goat anti-rabbit, goat anti-mouse or goat anti-human). Blots were imaged using an Odyssey Infrared Imager (LI-COR Biosciences) and analyzed with Image Studio Lite software.

#### Flow cytometry of infected cells

Adherent cells were recovered by trypsinization and washed in PBS with 2 mM EDTA (PBS/EDTA). Cells were stained with fixable Zombie UV Live/Dead dye (BioLegend) for 6 min at room temperature. Excess stain was quenched with FBS-complemented DMEM. Unbound antibody was washed off thoroughly and cells were fixed in 4% PFA before intracellular staining. For intracellular detection of SARS-CoV-2 nucleoprotein, cells were permeabilized for 15 min with intracellular staining perm wash buffer (BioLegend). Cells were then incubated with 1 μg ml−1 CR3009 SARS-CoV-2 cross-reactive antibody (a gift from L. McCoy) in permeabilization buffer for 30 min at room temperature, washed once and incubated with secondary Alexa Fluor 488-donkey-anti-human IgG (Jackson Labs). All samples were acquired on a BD Fortessa X20 using BD FACSDiva software. Data were analyzed using FlowJo v.10 (Tree Star).

### Immunofluorescence staining and image analysis

Infected cells were fixed using 4% PFA/formaldehyde for 1 hour (h) at room temperature and subsequently washed with PBS. A blocking step was carried out for 35h at room temperature with 10% goat serum/1%BSA/0.001 Triton-TX100 in PBS. IRF3 and dsRNA staining to was performed by primary incubation with rabbit-anti-IRF3 antibody (sc-33641, Santa Cruz), mouse-anti-dsRNA (MABE1134, Millipore) antibody for 18h and washed thoroughly in PBS. Primary antibody detection occurred using secondary anti-rabbit-AlexaFluor-488 and anti-mouse-AlexaFluor-568 conjugates (Jackson ImmunoResearch) for 1h. All cells were labeled with Hoechst33342 (H3570, Thermo Fisher). Images were acquired using the WiScan® Hermes 7-Colour High-Content Imaging System (IDEA Bio-Medical, Rehovot, Israel) at magnification 10X/0.4NA. Four channel automated acquisition was carried out sequentially. Images were acquired across a well area density resulting in 31 FOV/well and ∼20,000 cells. Images were pre-processed by applying a batch rolling ball background correction in FIJI ImageJ software package (*79*) prior to quantification. IRF3 translocation analysis was carried out using the Athena Image analysis software (IDEA Bio-Medical, Rehovot, Israel) and data post-processed in Python. Infected cell populations were determined by thresholding of populations with greater than 2 segmented dsRNA punctae.

### Culture and infection of primary differentiated human airway epithelial (HAE) cells

#### Culture

Primary normal (healthy) bronchial epithelial (NHBE-A) cells were cultured for five to seven passages and differentiated at an air-liquid interface as previously described (*14, 52*). After 21-24 days of differentiation, cells were used in infection experiments.

#### Infections

For primary HAE infections, cells were washed once with PBS on the apical side before addition of diluted virus stocks for 2-3 h at 37°C. Supernatant was then removed and cells were washed twice with PBS. All liquid was removed from the apical side and basal medium was replaced with fresh Pneumacult ALI medium for the duration of the experiment. Virus release was measured at the indicated time points by extracting viral RNA from apical PBS washes. To this end, cells were incubated with PBS for 30 minutes at 37°C at the indicated time points. Cellular RNA was collected by lysing cells at the experimental endpoint in RLT (Qiagen) supplemented with 0.1% beta-mercaptoethanol (Sigma) and extracted as previously described (*14*).

### Infection of A549-ACE2 with mutant rSARS-CoV-2 N D3L and -3 deletion

#### Cell culture

Vero E6 (ATCC, CRL-1586) and Vero TMPRSS2 cells (BPS Bioscience Cat# 78081) were maintained in Dulbecco’s modified Eagle’s medium (Corning) supplemented with 10% fetal bovine serum (Peak Serum), non-essential amino acids (Gibco), HEPES (Gibco) and penicillin/streptomycin (Corning) at 37 °C and 5% CO_2_. A549-ACE2 (*80*) were maintained in Dulbecco’s modified Eagle’s medium (Corning) supplemented with 10% fetal bovine serum (Peak Serum) and penicillin/streptomycin (Corning) at 37 °C and 5% CO_2_. All cell lines used in this study were regularly screened for Mycoplasma contamination, using the Universal Mycoplasma Detection Kit (ATCC, 30-1012K).

#### Viruses and reverse genetics

The bacterial artificial chromosome (BAC) harboring the entire viral genome of SARS-CoV-2 USA-WA1/2020 strain (Accession No. MN985325) was described previously (*81*). The mutations of D3L/-3A, D3L, and -3A were achieved in viral fragment 1 by site-directed mutagenesis. After confirming the mutations by Sanger sequencing, fragments containing the mutations were released and reassembled into the BAC using BamHI and RsrII restriction endonucleases. Then, the BAC containing the mutations were maxi prepared and transfected into Vero AT (ACE2/TMPRSS2) cells using Lipofectamine 2000 (ThermoFisher Scientific) according to the manufacturer’s instruction. At 12 hours post-transfection, the supernatant was replaced by post-infection media (DMEM+1% FBS+1% PSG), and the supernatant was collected at 72 hours post-transfection, aliquoted, labeled as P0, and stored at -80 °C. The P1 stocks were generated by infecting monolayer of Vero AT cells with the P0 stocks (MOI 0.001). At 48 hours post-infection, the cell culture supernatant was collected, clarified, aliquoted, labeled as P1, and stored at -80 °C. After confirmation by deep sequencing, the P1 stocks were titrated and used for experiments.

Primers used for generation of mutant rSARS-CoV-2

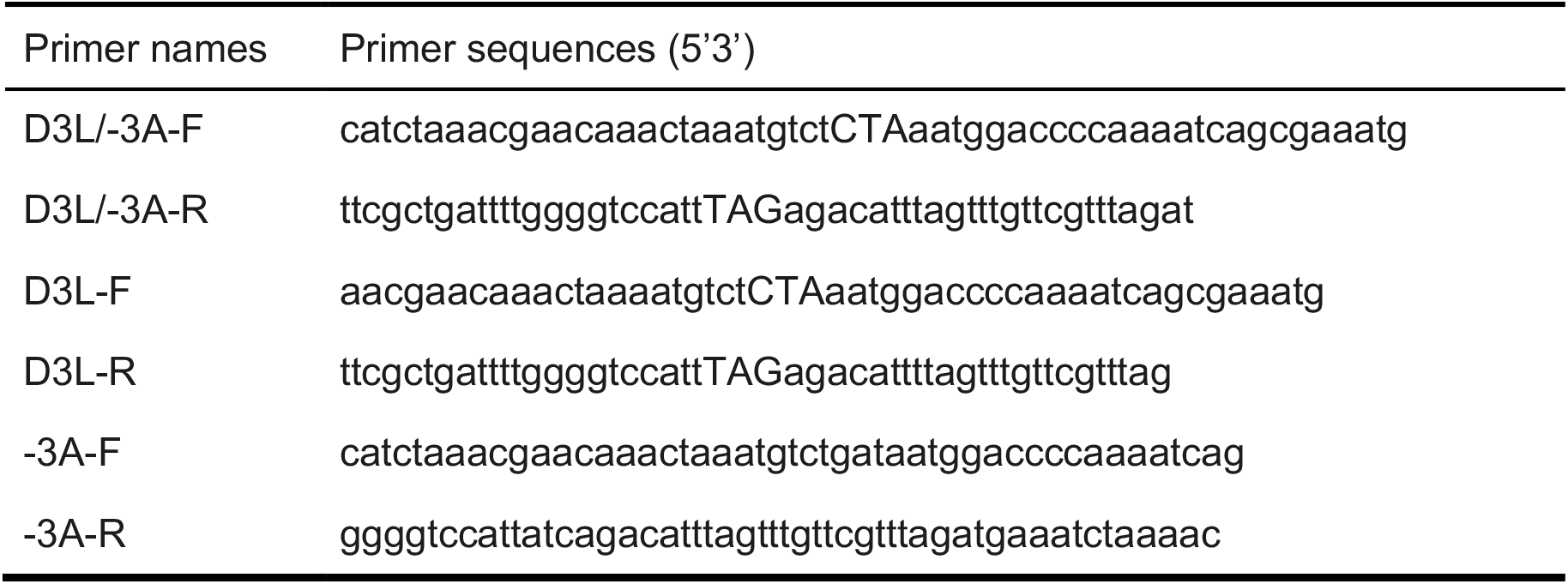

#### Infections

Unless otherwise specified, infections were performed in viral growth media (VGM): Dulbecco’s modified Eagle’s medium (Corning) supplemented with 2% fetal bovine serum (Peak Serum), non-essential amino acids (Gibco), HEPES (Gibco) and penicillin/streptomycin (Corning) at 37°C and 5% CO_2_ as previously described (*82*). For proteomic samples, one day before infection, 8×10^5^ A549-ACE2 cells per biological replicate were seeded in 6-well plates in complete media. Cells were then infected with the indicated viruses under BSL3 containment in accordance with the biosafety protocols developed by the Icahn School of Medicine at Mount Sinai. At the appropriate time post infection, cells were washed three times in ice cold 1x PBS and lysed in 500uL/well of 6M guanidine hydrochloride (Sigma) in 100mM Tris-HCl (pH 8.0). Samples were then boiled for 5 minutes at 95C to inactivate proteases, phosphatases, and virus. Samples were frozen at -80°C until further processing. For RNAseq analysis, one day before infection, 4×10^5^ A549-ACE2 cells per biological replicate were seeded in 6-well plates in complete media. Cells were then infected with the indicated viruses under BSL3 containment in accordance with the biosafety protocols developed by the Icahn School of Medicine at Mount Sinai. At the appropriate time post infection, cells were washed three times in ice cold 1x PBS and lysed by adding 1mL/well of Trizol reagent (Thermo Fisher).

### mRNA sequencing acquisition and analysis

#### RNA extraction, library preparation, and sequencing

RNA was extracted using the RNeasy Micro Kits (Qiagen) following manufacturers instructions or TRIzol (Invitrogen). Library preparation and sequencing were performed by Azenta Life Science using the following conditions: rRNA depletion for mRNA and long noncoding species, standard RNAseq run in Illumina® HiSeq 4000™ with a depth of 20-30 million reads per sample.

#### mRNA quantitative analysis

All reads were mapped to the human host genome (ensembl 101) using HISAT2 aligner (*83*). Host transcript abundances were estimated using human annotations (ensembl 101) using StringTie (*84*). Differential gene expression was calculated on the basis of read counts extracted for each protein-coding gene using featureCount and significance was determined by the DESeq2 R package (*85*). On average, we quantified ∼16,000–17,000 mRNA transcripts above 1 count per million reads.

#### SARS-CoV-2 genome reconstruction and evaluation from mRNA sequencing data

Forward and reverse reads were merged into one file and were not treated as paired-end for denovo assembly. Reads were normalized such that the maximum genome coverage would be no more than 200x using bbnorm from the bbmap package (https://jgi.doe.gov/data-and-tools/software-tools/bbtools/bb-tools-user-guide/bbmap-guide/). This aided assembly of genomes with high variability in coverage. Normalized reads were then assembled with SPAdes (*86*) (v3.15.3) with various k-mer sets ranging from 27 to 97. Assemblies were compared to the reference SARS-CoV-2 genome and complete assemblies of the entire genome were selected for further analysis.

### Infection proteomics in Calu-3 and A549-ACE2 cells acquisition and analysis

#### Mass spectrometry sample preparation

Samples were lysed in 6M guanidine hydrochloride (Sigma), boiled at 95°C for 5 minutes, and stored on dry ice. Lysed samples were thawed and sonicated using a probe sonicator 1x for 15 seconds at 20% amplitude. Insoluble material was pelleted by spinning samples at max speed (∼13,000 rpm) for 10 minutes. Supernatant was transferred to a new protein lo-bind tube and protein was quantified using a Bradford assay. Approximately 500ug of protein sample was used for further processing, starting with reduction and alkylation using a 1:10 sample volume of tris-(2-carboxyethyl) (TCEP) (10mM final) and 2-chloroacetamide (4.4mM final) for 5 minutes at 45°C with shaking. Prior to protein digestion, the 6M guanidine hydrochloride was diluted 1:6 with 100mM Tris-HCl pH8 to increase the activity of trypsin and LysC proteolytic enzymes, which were subsequently added at a 1:100 (wt/wt) enzyme-substrate ratio and placed in a 37°C water bath overnight (∼16-20 hours). Following digestion, 10% trifluoroacetic acid (TFA) was added to each sample to a final pH of ∼2. Samples were desalted using a vacuum manifold with 50mg Sep Pak C18 cartridges (Waters). Each cartridge was activated with 1 mL 80% acetonitrile (ACN)/0.1% TFA, then equilibrated with 3 × 1 mL of 0.1% TFA. Following sample loading, cartridges were washed with 3 × 1 mL of 0.1% TFA, and samples were eluted with 1 × 0.8 mL 50% ACN/0.25% formic acid (FA). Ten percent (10%) of the resulting volume (∼50μg) was reserved for protein abundance measurements, and the remainder was used for phosphopeptide enrichment (thus, the same starting material was used for the abundance proteomics and phosphoproteomics analysis). All samples were dried by vacuum centrifugation. For phosphopeptide enrichment of samples for phosphoproteomics, IMAC beads (Ni-NTA from Qiagen) were prepared by washing 3x with HPLC water, incubating for 30 minutes with 50mM EDTA pH 8.0 to strip the Ni, washing 3x with HPLC water, incubating with 50mM FeCl3 dissolved in 10% TFA for 30 minutes at room temperature with shaking, washing 3x with and resuspending in 0.1% TFA in 80% acetonitrile. Peptides were enriched for phosphorylated peptides using a King Fisher Flex (KFF). For a detailed KFF protocol, please contact the authors.

#### Mass spectrometry data acquisition

Digested samples were analyzed on an Orbitrap Exploris 480 mass spectrometry system (Thermo Fisher Scientific) equipped with an Easy nLC 1200 ultra-high pressure liquid chromatography system (Thermo Fisher Scientific) interfaced via a Nanospray Flex nanoelectrospray source. For all analyses, samples were injected on a C18 reverse phase column (25 cm × 75 μm packed with ReprosilPur 1.9-μm particles). Analytical columns were equilibrated with 6 μl of mobile phase A with a max pressure of 650 bar. For abundance proteomics (phosphoproteomics same unless indicated in parentheses) mobile phase A consisted of 0.1% FA, and mobile phase B consisted of 0.1% FA / 80% ACN. Peptides were separated by an organic gradient from 4% (2%) to 30% (25%) mobile phase B over 62 min followed by an increase to 45% (40%) B over 10 min, then held at 95% B for 8 min at a flow rate of 300 nl min−1. Data-independent analysis (DIA) was performed on abundance and phosphoproteomics samples using an 80 minute gradient. An MS scan at 60,000 resolving power over a scan range of 350–1100 *m*/*z*, a normalized AGC target of 300%, and an RF lens setting of 40%. This was followed by DIA scans at 15000 resolving power, using 20 *m*/*z* isolation windows over 350–1100 *m*/*z* at a normalized HCD collision energy of 30%. Loop control was set to All. To build a spectral library, one sample from each set of biological replicates was acquired in a data-dependent manner. Data-dependent analysis (DDA) was performed by acquiring a full scan over a *m*/*z* range of 350–1100 in the Orbitrap at 60,000 resolving power with a normalized AGC target of 300% and an RF lens setting of 40%. Dynamic exclusion was set to 45s, with a 10-ppm exclusion width setting. Peptides with charge states 2–6 were selected for MS/MS interrogation using higher-energy collisional dissociation (HCD), with 20 MS/MS scans per cycle. For phosphopeptide-enriched samples, MS/MS scans were analyzed in the Orbitrap using isolation width of 1.6 *m*/*z*, normalized HCD collision energy of 30%, and normalized AGC of 200% at a resolving power of 15,000 with a 22ms (40ms for phosphoproteomics) maximum ion injection time.

#### Proteomics data search

Mass spectra from each DDA dataset were used to build experiment-specific libraries for DIA searches using the Pulsar search engine integrated into Spectronaut v. 16.0.220606.53000 (Hawking) by searching against a database of Uniprot *Homo sapiens* sequences (downloaded 22 March 2022) and a SARS-CoV-2 proteome of 149 protein sequences spanning five variants of concern (Alpha 20I V1, Beta 20H V2, Gamma 20J V3, Delta 21J, and Omicron 21K) and 32 viral proteins, including N-star (N*). For Experiment 3, directDIA search settings were used. For protein abundance samples, data were searched using the default Biognosys (BGS) settings, variable modification of methionine oxidation, static modification of carbamidomethyl cysteine, and filtering to a final 1% false discovery rate (FDR) at the peptide, peptide spectrum match (PSM) and protein level. For phosphopeptide-enriched samples, BGS settings were modified to include phosphorylation of S, T and Y as a variable modification. The generated search libraries were used to search the DIA data. For protein abundance samples, default BGS settings were used, with no data normalization performed. For phosphopeptide-enriched samples, the PTM site localization score in Spectronaut was applied. Imputation was removed for all analyses.

#### Proteomics quantitative analysis

Quantitative analysis was performed in the R statistical programming language (v.4.1.3). Initial quality control analyses, including inter-run clusterings, correlations, principal component analysis (PCA), peptide and protein counts and intensities were completed with the R package artMS (v. 1.12.1). On the basis of obvious outliers in intensities, correlations and clusterings in PCA analysis, one run was discarded from the phosphoproteomics dataset: Experiment 1, IC10_10h.1. Statistical analysis of phosphorylation and protein abundance changes between mock and infected runs, as well as between infected runs from different variants were computed using peptide ion fragment data output from Spectronaut and processed using artMS. Quantification of phosphorylation differences was performed using artMS as a wrapper around MSstats, via functions artMS::doSiteConversion and artMS::artmsQuantification with default settings. All peptides containing the same set of phosphorylated sites were grouped and quantified together into phosphorylation site groups. For both phosphopeptide and protein abundance MSstats pipelines, MSstats performs normalization by median equalization, no imputation of missing values, and median smoothing to combine intensities for multiple peptide ions or fragments into a single intensity for their protein or phosphorylation site group. Lastly, statistical tests of differences in intensity between infected and control time points were performed. When not explicitly indicated, we used defaults for MSstats for adjusted *P* values, even in cases of *n* = 2. By default, MSstats uses the Student’s *t*-test for *P* value calculation and the Benjamini–Hochberg method of FDR estimation to adjust *P* values.

### Quantifying viral RNA and protein

#### Quantifying viral RNA

Coronavirus RNA transcripts were characterized by the junction of the leader with the downstream subgenomic sequence. Reads containing possible junctions were extracted by filtering for exact matches to the 3’ end of the leader sequence “CTTTCGATCTCTTGTAGATCTGTTCTC” or the mutated form “CTTT<u>T</u>GATCTCTTGTAGATCTGTTCTC” (mutation underlined) as appropriate for BA.2, BA.4 and BA.5 using the tool bbduk from the BBTools package (version 38.96; Bushnell B. - sourceforge.net/projects/bbmap/). This subset of leader-containing reads were then left-trimmed to remove the leader, also using bbduk. The filtered and trimmed reads were then matched against a full length genomic sequence with the bbmap tool from BBtools using settings (maxindel=100, strictmaxindel=t, local=t). The left-most mapped position in the reference was used as the junction site. Full length genome sequences were assembled de novo from all reads from a late time point sample of the same virus in the same batch using SPAdes (*86*), as described above. The position numbers between viral genomes were standardized to a reference SARS-CoV-2 sequence (accession NC_045512.2) using global pairwise alignments of full length sequences. Junction sites were labeled according to canonical locations of TRS sequences, or other known site with a +/-5 base pair window as follows (genomic = 67, S = 21553, orf3 = 25382, E = 26237, M = 26470, orf6 = 27041, orf7 = 27385, orf8 = 27885, N = 28257, orf9b = 28280, N* = 28878).

#### Quantifying viral proteins

Peptide ions or “features” (each charge state treated separately) that were likely false identifications based on an unexpected observation within the mock-infected samples were excluded. This exclusion was based on a categorical linear regression of log2 intensity values at 24 hours post infection using the R function lm per peptide ion, encoding category names to treat all mock-infected samples as the intercept, and requiring a significant coefficient on each and all of the virus samples (log2FC > 1, p-value < 0.05, two sided t-test). This exclusion was done separately per experiment. For the rSARS-CoV-2 N D3L and -3 deletion mutant virus experiment (exp. 6), we removed falsely identified peptide ions by required peptide ion intensities to be at least 10x greater in the average of infection conditions relative to the mock conditions. Once final features were selected, viral protein intensities were calculated as the sum of feature intensities that mapped to each viral protein (as determined by the Spectronaut search, above).

### Phosphorylation of viral proteins analysis

We started by systematically determining the amino acid mutations present within each VOC (Table S1) relative to the Wuhan reference, annotated by its position within each viral protein. Deletions or insertions were annotated in relation to the Wuhan reference. Phosphorylated peptide intensities were defined as the sum of phosphorylated peptide ion (i.e., same peptide could have different charge states or other modifications such as oxidation) intensities from the phosphoproteomics data (per biological replicate), requiring that summed peptides had identical phosphorylation site modification(s). For example, a peptide with two forms, phosphorylated on one site or two, were quantified separately. Importantly, only phosphorylated peptides that possessed a conserved sequence across each pair of viruses being compared were used for quantification of phosphorylated peptide intensities to control for physicochemical peptide properties that may affect their quantitation using mass spectrometry. Viral protein intensities were defined as the sum of peptide feature intensities from the abundance proteomics data that mapped to each viral protein (as determined by the Spectronaut search, above). We then calculated the ratio between each corresponding phosphorylated peptide intensity and viral protein intensity per biological replicate (phosphorylated peptide intensity / viral protein intensity). Next, we performed a two-tailed t-test, assuming unequal variance, generating a t-statistic and p-value per phosphorylated peptide. We required at least two biological replicates per condition to perform the t-test (see Table S10 for full set of results).

### Targeted proteomics for Orf9b phosphorylation

A spectral library was constructed from the DIA data to obtain Orf9b-specific transitions for abundance and phosphorylation measurements. Four Orf9b peptides were included to determine Orf9b abundance. To determine site-specific Orf9b phosphorylation, we included peptides representing Ser50 (LGS(+80)PLSLNMAR) and Ser53 (LGSPLS(+80)LNMAR). Unrelated heat shock peptides were included and used for normalization when appropriate. All samples were loaded on a 15 cm PepSep column (Bruker) packed with 1.5 μm, 150 Å ReproSil C18 particles connected to a nanoLC easy 1200 (Thermo Fisher Scientific). For abundance measurements, peptides were separated with the following 50 minute gradient at 0.6 μl min−1: 2% to 6% B (0.1% FA in 80% ACN) in 1 minute to 25% B in 35 minutes, 34% B over 4 minutes, 90% B over 1 minute, followed by a 10 minute wash at 90% B. Phosphorylated peptides were separated with the following 62 minute gradient at 0.6 μl min−1: 2% to 5% B in 2 minutes to 27% B at 32 minutes, 33% B over 15 minutes, 90% B over 1 minute, followed by a 12 minute wash at 90% B. All samples were acquired on an Orbitrap Tribrid Lumos (Thermo Fisher Scientific) operated in positive ion mode. For abundance samples, each MS1 scan was acquired at a Fourier transform (FT) resolution at 120,000 (200 m/z) from 350 to 1,100 m/z. Peptide ions were accumulated for 50 ms or until reaching an AGC of 5e5. Four peptides were placed on an inclusion list and fragmented using stepped HCD at an NCE of 33 and a spread of ±3%. Peptides were isolated with a 1.4 Da window, and the resultant fragments were analyzed in the Orbitrap at 60,000 resolution (200 m/z). For analysis of phosphorylated peptides, the mass spectrometer was operated in targeted mode (PRM). One FT MS1 scan at 120,000 resolution (200 m/z) from 500 to 800 m/z was acquired for 256 ms or until reaching an AGC of 1e6, followed by four targeted scans per cycle. Precursor isolation width was set to 1.6 Da, list and fragmented using stepped HCD at an NCE of 33 and a spread of ±3%. MS2 scans were acquired in the Orbitrap at a resolution of 60,000 (200 m/z), and ions were accumulated for 118 ms or until reaching an AGC of 5e5. Following MS acquisition, each experiment was separately analyzed in Skyline. For all analyses, the MS1 filter was set to count and three precursors were used at 10 ppm mass error. The MS2 filtering was set to Orbitrap and the resolution was set to 60,000 (200 m/z). For the phosphorylation site experiments both b/y and a/z ions were used, whereas for the abundance experiments b and y ions were included. Peaks were manually inspected for integration and boundaries refined if necessary. For Orf9b Ser50/Ser53 the presence of the proline in the peptide sequence resulted in a split chromatographic peak between the two isomers and the second peak was used for integration for all samples. For both phosphoisomers, only phophosite-specific ions were used for quantification (y5-y9/b6-b10 for Ser53 and y9-y5/b2-b6 for Ser50).

### *In vitro* kinase activity assay

ADP-Glo kinase reactions (Promega #V6930) were carried out per the manufacturer’s instructions using 10 nM TBK1 kinase (Promega #V3991) and 10 μM peptide substrate with 100 μM ATP. The assay reaction buffer consisted of 40 mM Tris pH 7.5, 20 mM MgCl_2_, 0.1 mg/mL BSA, and 10 μM DTT. A standard curve from 0 to 100 μM ADP was used for validation. Orf9b peptides were synthesized by GenScript using the WT sequence PIILRLGsPLsLNMARKt, with Ala substituted at the corresponding Ser and Thr sites. To account for TBK1 auto-phosphorylation a no substrate control was included. Myelin Basic Protein was included as a positive control.

### Mutation and structure analysis

#### Conservation

SARS-CoV-2 protein sequences were downloaded from Uniprot (*87*) and the orf1ab polyprotein split into sub-sequences based on the Uniprot annotation. A custom reference database was generated based on the NCBI virus coronavirus genomes dataset <u>(NCBI Resource Coordinators, 2018)</u>, which includes sequences from a large range of coronaviruses. SARS-CoV-2, SARS and MERS sequences were filtered to only contain sequences from the Wuhan-Hu-1 strain, the Urbani strain and the HCoV-EMC/2012 strain respectively. Without this the dataset contains very large numbers of almost identical sequences from patient samples, which are not informative since SIFT4G looks to compare across species. The remaining sequences were clustered using MMseqs2 (*88*) with an overlap threshold of 0.8 and a sequence identity threshold of 0.95, which grouped other duplicate sequences into representative clusters. SIFT4G Scores were generated for all possible variants to the SARS-CoV-2 sequences based on this database. A modified copy of SIFT4G was used, which reports scores to 5 decimal places instead of the usual 2.

#### Structural Destabilization

Structures were sourced from the SWISS-Model (*73, 89*) SARS-CoV-2 repository (https://swissmodel.expasy.org/repository/species/2697049), which contains experimental structures and homology models. Models were required to have greater than 30% sequence identity and a QMean score (*90*) greater than -4, as recommended by SWISS-Model. Suitable models were available for 19 of the 28 viral proteins. Models were ordered by priority; firstly experimental models over homology models and then by QMean Score. Models were examined in turn and any position not covered by a higher priority model was added to the FoldX analysis pipeline. FoldX’s RepairPDB command was used to pre-process selected SWISS-Model PDB files. All mutations at each position assigned to each model were modeled using the BuildModel command, using the average ΔΔG prediction from three runs.

#### Surface Accessibility

Naccess (Hubbard and Thornton, 1993) was run on each structure using the default settings. Structures were filtered to only include the chain corresponding to the appropriate SARS-CoV-2 protein. This means some surface accessible positions are usually found in interfaces rather than facing the solvent. Since structures are not always complete, surface accessibility is an approximation and will not be accurate in all cases. The code managing the pipeline and analyses are available at https://github.com/allydunham/mutfunc_sars_cov_2. The web service source code is available at https://github.com/allydunham/mutfunc_sars_cov_2_frontend. The pipeline is managed through Snakemake (*91*). For more information about the mutation and structure analysis, see prior work (*92*).

### Affinity purification mass spectrometry (APMS) experiments

#### Constructs

Mutagenesis and subsequent plasmid preparation were performed by Genscript Biotech. Each sequence was codon optimized for expression in mammalian cells by Genscript Biotech. All plasmids were cloned in the lentiviral constitutive expression vector pLVX-EF1alpha-IRES-Puro (Takara Bio). Constructs are available from the authors upon request.

#### Transfections

HEK-293T cells were cultured in Dulbecco’s modified Eagle’s medium (DMEM; Corning) supplemented with 10% fetal bovine serum (FBS; Gibco, Life Technologies) and 1% penicillin– streptomycin (Corning) and maintained at 37 °C in a humidified atmosphere of 5% CO2. HEK-293T cells were procured from the UCSF Cell Culture Facility, now available through UCSF’s Cell and Genome Engineering Core ((https://cgec.ucsf.edu/cell-culture-and-banking-services); cell line collection listed here: https://ucsf.app.box.com/s/6xkydeqhr8a2xes0mbo2333i3k1lndqv (CCLZR076)). STR analysis by the Berkeley Cell Culture Facility on 8 August 2017 authenticated HEK-293T cells with 94% probability. Cells were tested on 4 October 2021 using the MycoAlert PLUS Mycoplasma Detection Kit (Lonza LT07-710) and were negative: B/A ratio < 1 (no detected mycoplasma). For each bait, n = 3 independent biological replicates were prepared for affinity purification and seven million HEK293T cells were plated per 15-cm dish. The WT and mutant baits for each SARS-CoV-2 protein were transfected as one experiment on the same day, and one GFP control and one empty vector (pLVX-EF1α-IRES-Puro) control were included in each experiment. The amount of bait plasmid transfected was determined based on previous affinity purification experiments of wave 1 only SARS-CoV-2 baits (2020 Nature paper). Total plasmid was topped up to 15 μg with empty vector and the following amounts of each bait or control plasmid were transfected: 2 μg GFP; 2.5 μg N, Orf3a, Orf9b; 5.0 μg Orf7b, Nsp2, Nsp6; 6.125 μg Nsp3; 7.5 μg E, Orf8, Nsp7, Nsp9, Nsp10, Nsp16; 10 μg Orf6, Nsp4, Nsp5, Nsp8, Nsp11, Nsp15; 12.5 μg S, M, Nsp12, Nsp14; 15 μg Orf3b, Orf7a, Orf9c, Orf10, Nsp1, Nsp13, empty vector. Each 15 μg plasmid per 15-cm dish was complexed with PolyJet Transfection Reagent (SignaGen Laboratories) at a 1:3 μg:μl ratio of plasmid:transfection reagent based on the manufacturer’s recommendations. After more than 38h, cells were dissociated on ice using 10 ml Dulbecco’s phosphate-buffered saline without calcium and magnesium (DPBS) supplemented with 10 mM EDTA for at least 5 minutes and subsequently washed three times with 10 ml DPBS. Each step was followed by centrifugation at 200g, 4 °C for 5 min. Cell pellets were frozen immediately on dry ice and stored at −80°C.

#### Affinity purification and sample preparation

Frozen cell pellets were thawed on ice for 15–20 min and resuspended in 1 ml IP lysis buffer. IP lysis buffer (50 mM Tris-HCl, pH 7.4 at 4 °C, 150 mM NaCl, 1 mM EDTA) was supplemented immediately before use with 0.5% Non-idet P40 substitute (NP40; Fluka Analytical) and cOmplete mini EDTA-free protease and PhosSTOP phosphatase inhibitor cocktails (Roche). Samples were then frozen on dry ice for 10–20 min and partially thawed at 37 °C before incubation on a tube rotator for 30 min at 4 °C and centrifugation at 13,000g, 4 °C for 15 min to pellet debris. Up to 96 samples were arrayed into a 96-well Deepwell plate for affinity purification on the KingFisher Flex (KFF) Purification System (Thermo Scientific) as follows: MagStrep ‘type3’ beads (30 μl; IBA Lifesciences) were equilibrated twice with 1 ml wash buffer (IP buffer supplemented with 0.05% NP40) and incubated with 0.95 ml lysate for 2h. Beads were washed three times with 1 ml wash buffer and then once with 1 ml IP buffer. To directly digest bead-bound proteins as well as elute proteins with biotin, beads were manually suspended in IP buffer and divided in half before transferring to 50 μl denaturation–reduction buffer (2 M urea, 50 mM Tris-HCl pH 8.0, 1 mM DTT) and 50 μl 1× buffer BXT (IBA Lifesciences) dispensed into a single 96-well KFF microtitre plate. Purified proteins were first eluted at room temperature for 30 min with constant shaking at 1,100 rpm on a ThermoMixer C incubator. After removing eluates, on-bead digestion proceeded. Strep-tagged protein expression in lysates and enrichment in eluates was assessed by western blot and silver stain, respectively. The KFF Purification System was placed in the cold room and allowed to equilibrate to 4 °C overnight before use. All automated protocol steps were performed using the slow mix speed and the following mix times: 30 s for equilibration and wash steps, 2h for binding and 1 min for final bead release. Three 10s bead collection times were used between all steps. Bead-bound proteins were denatured and reduced at 37°C for 30 min and, after being brought to room temperature, alkylated in the dark with 3 mM iodoacetamide for 45 min and quenched with 3 mM DTT for 10 min. Proteins were then incubated at 37 °C, initially for 4 h with 1.5 μl trypsin (0.5 μg/μl; Promega) and then another 1–2 h with 0.5 μl additional trypsin. To offset evaporation, 15 μl 50 mM Tris-HCl, pH 8.0 was added to each sample before trypsin digestion. All steps were performed with constant shaking at 1,100 rpm on a ThermoMixer C incubator. Resulting peptides were combined with 50 μl 50 mM Tris-HCl, pH 8.0 to rinse beads and then acidified with trifluoroacetic acid (0.5% final, pH < 2.0). Acidified peptides were desalted for MS analysis using a BioPureSPE Mini 96-Well Plate (20 mg PROTO 300 C18; The Nest Group) according to standard protocols.

#### Mass spectrometry acquisition and search

Samples were resuspended in 0.1% formic acid and analyzed on a Q-Exactive Plus mass spectrometry system (Thermo Fisher Scientific) equipped with an Easy nLC 1200 ultra-high pressure liquid chromatography system (Thermo Fisher Scientific) interfaced via a Nanospray Flex nanoelectrospray source. For all analyses, samples were injected on a C18 reverse phase column (25 cm × 75 μm packed with ReprosilPur 1.9-μm particles). Analytical columns were equilibrated with 5 μl of mobile phase A with a max pressure of 650 bar. Mobile phase A consisted of 0.1% FA, and mobile phase B consisted of 0.1% FA / 80% ACN. Peptides were separated by an organic gradient from 2% to 7% mobile phase B over 1 minute, followed by an increase to 36% B over 53 min, then held at 95% B for 13 min, then reduced back down to 2% B for 11 minutes at a flow rate of 300 nl min−1. Data-dependent analysis (DDA) was performed by acquiring a full scan over a *m*/*z* range of 300–1500 in the Orbitrap at 70,000 resolving power with an AGC target of 1e6. Top 20 peptides were selected for MS/MS interrogation using in-source collision-induced dissociation (CID), with 20 MS/MS scans per cycle and 17,500 resolving power. Resulting mass spectra were searched using MaxQuant (v.1.6.12.0) (*93, 94*) using default settings and searching against a database of Uniprot *Homo sapiens* sequences (downloaded 22 March 2022), a SARS-CoV-2 proteome of 149 protein sequences spanning five variants of concern (Alpha 20I V1, Beta 20H V2, Gamma 20J V3, Delta 21J, and Omicron 21K) and 32 viral proteins, including N-star (N*), and the enhanced green fluorescence protein (eGFP) sequence. Detected peptides and proteins were filtered to 1% false-discovery rate in MaxQuant.

#### Scoring high-confidence protein-protein interactions

Identified proteins were subjected to protein–protein interaction scoring with both SAINTexpress (v.3.6.3)(*31*) and MiST (https://github.com/kroganlab/mist) (*32, 95*) scoring algorithms. For SAINT scoring, each APMS experimental batch was scored separately relative to independent negative controls: empty vector (EV) and enhanced green fluorescence protein overexpression (eGFP). For MiST scoring, all samples across all batches were scored together. Three scoring thresholds were generated, all included the requirement of average spectral counts (AvgSpec) to be greater than two: (i) SAINT only requiring SAINT BFDR<0.05, (ii) SAINT and MiST requiring BFDR<0.05 & MiST>0.5, and our final selected threshold (iii) SAINT, MiST, and Gordon et al. (2020) requiring [BFDR<0.05 & MiST>0.5] | [BFDR<0.05 & present in Gordon et al. (2020)]. These thresholds had to be satisfied in either the mutant or wave 1, but not both, for every mutant and wave 1 pair. Although (iii) is our final, most high-confidence, scoring scheme, interactions in (i) and (ii) are worth consideration for followup studies (see Table S11 for full list of unthresholded and high-confidence interactions).

#### Differential protein-protein interaction analysis and visualization

We used the R package MSstats (*33*) to quantify changes in “prey” protein abundance between the corresponding mutant (VOC) and wave 1 “bait” forms. We first converted MaxQuant evidence files to MSstats format using MaxQtoMSstatsFormat with proteinID=”Leading.razor.protein”, useUniquePeptides=FALSE, summaryforMultipleRows=sum, removeFewMeasurements=FALSE, removeOxidationMpeptides=FALSE, and removeProtein_with1Peptide=FALSE. We then ran the dataProcess function with featureSubset=”all”, normalization=”equalizeMedians”, MBimpute=FALSE, and summaryMethod=“TMP”. In essence, we performed normalization by median equalization, did not impute missing values, and combined (i.e., “summarized”) intensities for multiple peptide ions or fragments into a single intensity for their protein group. Lastly, statistical tests of differences in intensity between mutant and wave 1 were performed. We used defaults for MSstats for adjusted *P* values, even in cases of *n* = 2. By default, MSstats uses the Student’s *t*-test for *P* value calculation and the Benjamini–Hochberg method of FDR estimation to adjust *P* values. This analysis resulted in log2 fold changes (log2FC) and p-values per interaction between corresponding mutant and wave 1 baits. We defined differential interactions based on two criteria: (1) The prey must be a high-confidence interaction in either the mutant or wave 1 bait (see scoring thresholds above) and (2) the prey must be changing in abundance between the mutant and wave 1 forms with an absolute value log2FC>0.05 and p<0.05. For all proteins that fulfilled these criteria, we extracted information about the stable protein complexes that they participated in from the CORUM database of known protein complexes (*28*). We visualized differential interactions and CORUM complexes using Cytoscape (v.3.8.0) (*96*). We exported the cytoscape network as PDF and imported the PDF into Adobe Illustrator to refine the aesthetics and add annotations for protein complexes and biological processes. Biological process terms were manually refined from a set of GO Biological Process enrichments acquired using the clusterProfiler package (*97*) in R.

#### AP-western analysis

For affinity purification (AP) western blot analysis, HEK293T cells were transfected with the indicated plasmid constructs using lipofectamine 2000 (Invitrogen). Twenty four (24) hours post-transfection, cells were harvested in NP-40 lysis buffer (50mM Tris-Cl pH 7.5, 150mM NaCl, 0.5% NP-40, 1mM EDTA) supplemented with cOmplete mini protease inhibitor cocktail and PhosSTOP phosphatase inhibitor cocktail (Roche). Clarified cell lysates were incubated with MagStrep ‘type3’ beads (IBA Lifesciences) for 2-4 h at 4°C, followed by washing the magnetic beads with NP-40 lysis buffer for five times. Protein complexes were eluted by direct incubation in 1X SDS loading buffer and heating at 95 °C. Eluates and whole-cell lysates were analyzed by western blotting using the indicated antibodies. The ORF clone for Flag-MARK2 was obtained from Genscript (#OHu23943D). The antibodies used in the study include: rabbit polyclonal anti-beta-actin (Cell Signaling Technology #4967, RRID:AB_330288, used at 1:2000); mouse monoclonal anti-Strep tag (QIAGEN #34850, RRID:AB_2810987, used at 1:1000); rabbit polyclonal anti-flag antibody (Sigma Aldrich #F7425, RRID:AB_439687, used at 1:2000); rabbit polyclonal anti-LEO1 antibody (Atlas antibodies #HPA040741, RRID:AB_10794859, used at 1:1000); rabbit polyclonal anti-WDR61 antibody (Atlas antibodies # HPA040065, RRID: AB_10793926, used at 1:1000); rabbit polyclonal anti-PAF1 antibody (Proteintech # 15441-1-AP, RRID: AB_2174457, used at 1:1000); rabbit polyclonal anti-CTR9 antibody (Proteintech # 21264-1-AP, RRID: AB_10734585, used at 1:1000).

### Integrative computational analysis

#### Defining genes and functional modules involved in host response

Infection-regulated host genes were defined by requiring significant changes at any time point versus time-matched mocks using thresholds of absolute log2FC > 1 and p-value < 0.001 for phosphoproteomics and transcriptomics. Using a single unadjusted p-value, as opposed to a FDR threshold, across all viruses allowed for the application of consistent thresholds per virus without favoring any while still keeping the FDR to a reasonable level (0.00x to 0.00x for all contrasts) based on the high number of regulated genes. For phosphoproteomics, we further limited the set to only the likely functionally-important phosphorylation sites by requiring a functional score greater than 0.4 according to Ochoa *et al*. (*98*). For the proteomics data with lower number of regulated proteins, there was more need to control the FDR, so we used thresholds of absolute log2FC > log2(1.5) and FDR-adjusted (Benjamini-Hochberg) p-value < 0.05. Additionally, within each of the three different omics, a gene or phosphosite had to pass these thresholds twice at either different time points, viruses, or batches.

All genes that passed thresholds of regulation in any omics in batches Experiment 1 and 2 were pooled to define the total set of infection-regulated genes. A subnetwork of STRING (version 11.5) was then extracted that contained all matching STRING proteins using STRING’s gene alias table, and all edges between any pair of these STRING proteins with composite score greater than 0.6. A matrix of network distances between all genes was calculated using Diffusion State Distance (DSD) (*99*). DSD evaluates inter-gene network distance based on differences of diffusion by random walks from the two genes, and was a major portion of the best overall method for network module detection in Choobdar *et al*. (*100*). The DSD distance matrix was used as input to the R function hclust with agglomeration method set to “average”. The resultant hierarchical clustering was divided into modules by the function cutreeHybrid in R package dynamicTreeCut (version 1.63-1) with deepSplit set to 3. Modularization was visualized by coloring genes by module on a t-SNE plot of the regulated genes built using the R package Rtsne (version 0.16) with the DSD distance matrix as input. Modules were named by performing over representation analysis on sets of genes annotated by all gene ontology terms using the R package clusterProfiler (version 4.4.1) and the R annotations package org.Hs.eg.db (version 3.15.0). From among the eight GO terms with the lowest p-value per module, the term with the greatest number of genes was chosen for the module’s name, with ties broken by lowest p-value.

#### Correlations of modules

For each unique combination of batch, virus, timepoint, and omics, the response of a module to infection was calculated as the mean of all observed genes’ absolute log2FC vs mock. The mean of all modules’ values from transcriptomics was used as the Average Host Response (AHR). Correlation values between modules, using each of proteomics, phosphoproteomics and transcriptomics separately, and the AHR or virus protein levels were calculated as Pearson’s R using the values from the 11 samples from 7 different viruses in experiments 1 and 2. Each module gets a separate correlation to the AHR for its proteomics, phosphoproteomics and transcriptomics. These three different correlations were summarized to a composite R value using geometric mean to rank them for their correlation to AHR. The geometric mean was used to favor consistently high correlation across the three omics. To rank those least correlated, we defined the composite R value by first transforming by subtracting from 1, taking the geometric mean, and then subtracting from 1 again to convert back to a similar scale as the original R values. This transformation allowed for the handling of negative R values and still enabled the property of geometric means to favor consistently high numbers to find those that are consistently low. For correlating modules with virus protein intensity, the log_2_ virus protein intensities were used.

#### Network propagation analysis

We downloaded the human physical protein-protein interaction network from String, version 11.5 (*101*). We focused on genes with full RNA data across the 11 conditions whose encoded proteins are present in the network. For each gene and each condition, we set its log_2_ fold-change (24hpi relative to mock) in absolute value as a prior value for that condition. We subjected these values to a network propagation process with symmetric normalization and a parameter of 0.5, which smooths the values across the network while accounting for their prior values (*102*). The resulting values were averaged for every module and condition and compared to these averages across all modules (AHR, average host response). We further computed nominal p-values per module by comparing a module’s AHR correlation value to those obtained by randomizing module assignment while preserving module sizes. Similar analyses were conducted for abundance and phosphoproteomics data by completing missing (prior) data with zeros.

### Plitidepsin treatment of SARS-CoV-2 VOC-infected mice

Animal studies using the K18-hACE2 (Strain #034860 from the Jackson Laboratories) were performed in animal biosafety level 3 (BSL3) facility at the Icahn school of Medicine in Mount Sinai Hospital, New York City. All work was conducted under protocols approved by the Institutional Animal Care and Use Committee (IACUC). We utilized five female mice 4-week-old specific pathogen–free per group. Mice were anesthetized with a mixture of ketamine/xylazine before each intranasal infection with different SARS-CoV-2 strains (Alpha, Beta, Gamma, Delta. Omicron and Washington strain, WA). Four (4) days post-infection (dpi) animals were humanely euthanized. Lungs were harvested for viral titration. Silica glass beads tubes were used for the homogenization. For viral titers, whole lung was homogenized in PBS then frozen at −80°C for viral titration via TCID50. Briefly, infectious supernatants were collected at 48 h post-infection and frozen at −80 °C until later use. Infectious titers were quantified by limiting dilution titration using Vero TMPRSS2 cells. Briefly, Vero-TMPRSS2 cells were seeded in 96-well plates at 20,000 cells/well. The next day, SARS-CoV-2-containing supernatant was applied at serial 10-fold dilutions ranging from 10−1 to 10−8 and, after 4 days, viral cytopathic effect (CPE) was detected by staining cell monolayers with crystal violet. TCID50/ml were calculated using the method of Reed and Muench. The R statistical software (package rstatix) was used to determine differences in lung titers using a two-tailed, unpaired t-test assuming equal variances at a confidence interval of 95%. Not all infections were successful in the mice. We filtered outliers using a Hampel filter.

### Virus-like particle (VLP) assays

Virus-like particle assays were performed as previously described (*22, 23*). Plasmids CoV2-N (0.67), CoV2-M-IRES-E (0.33), CoV-2-S (0.0016) and Luc-T20 (1.0) at indicated mass ratios for a total of 375 ng of DNA were diluted in 37.5 µL Opti-MEM containing 1.125 µg PEI (polyethyleneimine, Polysciences). For experiments with N*, mass ratios were 0.33 (N), 0.33 (N*), 0.33 (M-IRES-E), 0.0016 (S), Luc-T20 (1.0). Transfection mixture was incubated for 20 minutes at room temperature and then added to HEK293T cells in each well of a 96-well plate containing 150 µL of DMEM (with fetal bovine serum and penicillin/streptomycin). Media was changed after 24 hours of transfection. At 48 hours post-transfection, VLP containing supernatant was collected and filtered using a 0.45 µm pore size filter plate. VLP containing supernatant (50 µL) was added to 50 µL of cell suspension containing 50,000 receiver cells (HEK293T ACE2/TMPRSS2) in each well of tissue culture treated opaque white 96-well plate (Corning). Cells were allowed to attach and take up VLPs overnight. The next day, supernatant was removed, and cells were lysed in 20 µL passive lysis buffer (Promega) for 15 minutes at room temperature with gentle rocking. Using a TECAN Spark plate reader, 50 µL of reconstituted luciferase assay buffer (Promega) was added to each well and mixed for 15 seconds followed immediately by luminescence measurement. Expression of VOC N protein constructs was evaluated previously (*22*) and found to be equivalent across constructs (*23*).

## REFERENCES

1. Weekly epidemiological update on COVID-19 - 6 January 2022, (available at https://www.who.int/publications/m/item/weekly-epidemiological-update-on-covid-19---6-january-2022).

2. E. Volz, S. Mishra, M. Chand, J. C. Barrett, R. Johnson, L. Geidelberg, W. R. Hinsley, D. J. Laydon, G. Dabrera, Á. O’Toole, R. Amato, M. Ragonnet-Cronin, I. Harrison, B. Jackson, C. V. Ariani, O. Boyd, N. J. Loman, J. T. McCrone, S. Gonçalves, D. Jorgensen, R. Myers, V. Hill, D. K. Jackson, K. Gaythorpe, N. Groves, J. Sillitoe, D. P. Kwiatkowski, COVID-19 Genomics UK (COG-UK) consortium, S. Flaxman, O. Ratmann, S. Bhatt, S. Hopkins, A. Gandy, A. Rambaut, N. M. Ferguson, Assessing transmissibility of SARS-CoV-2 lineage B.1.1.7 in England. Nature. 593, 266–269 (2021).

3. N. G. Davies, S. Abbott, R. C. Barnard, C. I. Jarvis, A. J. Kucharski, J. D. Munday, C. A. B. Pearson, T. W. Russell, D. C. Tully, A. D. Washburne, T. Wenseleers, A. Gimma, W. Waites, K. L. M. Wong, K. van Zandvoort, J. D. Silverman, CMMID COVID-19 Working Group, COVID-19 Genomics UK (COG-UK) Consortium, K. Diaz-Ordaz, R. Keogh, R. M. Eggo, S. Funk, M. Jit, K. E. Atkins, W. J. Edmunds, Estimated transmissibility and impact of SARS-CoV-2 lineage B.1.1.7 in England. Science. 372 (2021), doi:10.1126/science.abg3055.

4. N. L. Washington, K. Gangavarapu, M. Zeller, A. Bolze, E. T. Cirulli, K. M. Schiabor Barrett, B. B. Larsen, C. Anderson, S. White, T. Cassens, S. Jacobs, G. Levan, J. Nguyen, J. M. Ramirez 3rd, C. Rivera-Garcia, E. Sandoval, X. Wang, D. Wong, E. Spencer, R. Robles-Sikisaka, E. Kurzban, L. D. Hughes, X. Deng, C. Wang, V. Servellita, H. Valentine, P. De Hoff, P. Seaver, S. Sathe, K. Gietzen, B. Sickler, J. Antico, K. Hoon, J. Liu, A. Harding, O. Bakhtar, T. Basler, B. Austin, D. MacCannell, M. Isaksson, P. G. Febbo, D. Becker, M. Laurent, E. McDonald, G. W. Yeo, R. Knight, L. C. Laurent, E. de Feo, M. Worobey, C. Y. Chiu, M. A. Suchard, J. T. Lu, W. Lee, K. G. Andersen, Emergence and rapid transmission of SARS-CoV-2 B.1.1.7 in the United States. Cell. 184, 2587–2594.e7 (2021).

5. N. R. Faria, T. A. Mellan, C. Whittaker, I. M. Claro, D. da S. Candido, S. Mishra, M. A. E. Crispim, F. C. S. Sales, I. Hawryluk, J. T. McCrone, R. J. G. Hulswit, L. A. M. Franco, M. S. Ramundo, J. G. de Jesus, P. S. Andrade, T. M. Coletti, G. M. Ferreira, C. A. M. Silva, E. R. Manuli, R. H. M. Pereira, P. S. Peixoto, M. U. G. Kraemer, N. Gaburo Jr, C. da C. Camilo, H. Hoeltgebaum, W. M. Souza, E. C. Rocha, L. M. de Souza, M. C. de Pinho, L. J. T. Araujo, F. S. V. Malta, A. B. de Lima, J. do P. Silva, D. A. G. Zauli, A. C. de S. Ferreira, R. P. Schnekenberg, D. J. Laydon, P. G. T. Walker, H. M. Schlüter, A. L. P. Dos Santos, M. S. Vidal, V. S. Del Caro, R. M. F. Filho, H. M. Dos Santos, R. S. Aguiar, J. L. Proença-Modena, B. Nelson, J. A. Hay, M. Monod, X. Miscouridou, H. Coupland, R. Sonabend, M. Vollmer, A. Gandy, C. A. Prete Jr, V. H. Nascimento, M. A. Suchard, T. A. Bowden, S. L. K. Pond, C.-H. Wu, O. Ratmann, N. M. Ferguson, C. Dye, N. J. Loman, P. Lemey, A. Rambaut, N. A. Fraiji, M. do P. S. S. Carvalho, O. G. Pybus, S. Flaxman, S. Bhatt, E. C. Sabino, Genomics and epidemiology of the P.1 SARS-CoV-2 lineage in Manaus, Brazil. Science. 372, 815–821 (2021).

6. H. Tegally, E. Wilkinson, M. Giovanetti, A. Iranzadeh, V. Fonseca, J. Giandhari, D. Doolabh, S. Pillay, E. J. San, N. Msomi, K. Mlisana, A. von Gottberg, S. Walaza, M. Allam, A. Ismail, T. Mohale, A. J. Glass, S. Engelbrecht, G. Van Zyl, W. Preiser, F. Petruccione, A. Sigal, D. Hardie, G. Marais, M. Hsiao, S. Korsman, M.-A. Davies, L. Tyers, I. Mudau, D. York, C. Maslo, D. Goedhals, S. Abrahams, O. Laguda-Akingba, A. Alisoltani-Dehkordi, A. Godzik, C. K. Wibmer, B. T. Sewell, J. Lourenço, L. C. J. Alcantara, S. L. Kosakovsky Pond, S. Weaver, D. Martin, R. J. Lessells, J. N. Bhiman, C. Williamson, T. de Oliveira, Emergence and rapid spread of a new severe acute respiratory syndrome-related coronavirus 2 (SARS-CoV-2) lineage with multiple spike mutations in South Africa. medRxiv, 2020.12.21.20248640 (2020).

7. S. Cherian, V. Potdar, S. Jadhav, P. Yadav, N. Gupta, M. Das, P. Rakshit, S. Singh, P. Abraham, S. Panda, NIC team, Convergent evolution of SARS-CoV-2 spike mutations, L452R, E484Q and P681R, in the second wave of COVID-19 in Maharashtra, India. bioRxiv (2021), p. 2021.04.22.440932.

8. D. Planas, N. Saunders, P. Maes, F. Guivel-Benhassine, C. Planchais, J. Buchrieser, W.-H. Bolland, F. Porrot, I. Staropoli, F. Lemoine, H. Péré, D. Veyer, J. Puech, J. Rodary, G. Baele, S. Dellicour, J. Raymenants, S. Gorissen, C. Geenen, B. Vanmechelen, T. Wawina-Bokalanga, J. Martí-Carreras, L. Cuypers, A. Sève, L. Hocqueloux, T. Prazuck, F. A. Rey, E. Simon-Loriere, T. Bruel, H. Mouquet, E. André, O. Schwartz, Considerable escape of SARS-CoV-2 Omicron to antibody neutralization. Nature. 602, 671–675 (2022).

9. R. Viana, S. Moyo, D. G. Amoako, H. Tegally, C. Scheepers, C. L. Althaus, U. J. Anyaneji, P. A. Bester, M. F. Boni, M. Chand, W. T. Choga, R. Colquhoun, M. Davids, K. Deforche, D. Doolabh, L. du Plessis, S. Engelbrecht, J. Everatt, J. Giandhari, M. Giovanetti, D. Hardie, V. Hill, N.-Y. Hsiao, A. Iranzadeh, A. Ismail, C. Joseph, R. Joseph, L. Koopile, S. L. Kosakovsky Pond, M. U. G. Kraemer, L. Kuate-Lere, O. Laguda-Akingba, O. Lesetedi-Mafoko, R. J. Lessells, S. Lockman, A. G. Lucaci, A. Maharaj, B. Mahlangu, T. Maponga, K. Mahlakwane, Z. Makatini, G. Marais, D. Maruapula, K. Masupu, M. Matshaba, S. Mayaphi, N. Mbhele, M. B. Mbulawa, A. Mendes, K. Mlisana, A. Mnguni, T. Mohale, M. Moir, K. Moruisi, M. Mosepele, G. Motsatsi, M. S. Motswaledi, T. Mphoyakgosi, N. Msomi, P. N. Mwangi, Y. Naidoo, N. Ntuli, M. Nyaga, L. Olubayo, S. Pillay, B. Radibe, Y. Ramphal, U. Ramphal, J. E. San, L. Scott, R. Shapiro, L. Singh, P. Smith-Lawrence, W. Stevens, A. Strydom, K. Subramoney, N. Tebeila, D. Tshiabuila, J. Tsui, S. van Wyk, S. Weaver, C. K. Wibmer, E. Wilkinson, N. Wolter, A. E. Zarebski, B. Zuze, D. Goedhals, W. Preiser, F. Treurnicht, M. Venter, C. Williamson, O. G. Pybus, J. Bhiman, A. Glass, D. P. Martin, A. Rambaut, S. Gaseitsiwe, A. von Gottberg, T. de Oliveira, Rapid epidemic expansion of the SARS-CoV-2 Omicron variant in southern Africa. Nature. 603, 679–686 (2022).

10. M. Hoffmann, N. Krüger, S. Schulz, A. Cossmann, C. Rocha, A. Kempf, I. Nehlmeier, L. Graichen, A.-S. Moldenhauer, M. S. Winkler, M. Lier, A. Dopfer-Jablonka, H.-M. Jäck, G. M. N. Behrens, S. Pöhlmann, The Omicron variant is highly resistant against antibody-mediated neutralization: Implications for control of the COVID-19 pandemic. Cell. 185, 447–456.e11 (2022).

11. T. P. Peacock, J. C. Brown, J. Zhou, N. Thakur, K. Sukhova, J. Newman, R. Kugathasan, A. W. C. Yan, W. Furnon, G. De Lorenzo, V. M. Cowton, D. Reuss, M. Moshe, J. L. Quantrill, O. K. Platt, M. Kaforou, A. H. Patel, M. Palmarini, D. Bailey, W. S. Barclay, The altered entry pathway and antigenic distance of the SARS-CoV-2 Omicron variant map to separate domains of spike protein,, doi:10.1101/2021.12.31.474653.

12. H. Tegally, M. Moir, J. Everatt, M. Giovanetti, C. Scheepers, E. Wilkinson, K. Subramoney, Z. Makatini, S. Moyo, D. G. Amoako, C. Baxter, C. L. Althaus, U. J. Anyaneji, D. Kekana, R. Viana, J. Giandhari, R. J. Lessells, T. Maponga, D. Maruapula, W. Choga, M. Matshaba, M. B. Mbulawa, N. Msomi, NGS-SA consortium, Y. Naidoo, S. Pillay, T. J. Sanko, J. E. San, L. Scott, L. Singh, N. A. Magini, P. Smith-Lawrence, W. Stevens, G. Dor, D. Tshiabuila, N. Wolter, W. Preiser, F. K. Treurnicht, M. Venter, G. Chiloane, C. McIntyre, A. O’Toole, C. Ruis, T. P. Peacock, C. Roemer, S. L. Kosakovsky Pond, C. Williamson, O. G. Pybus, J. N. Bhiman, A. Glass, D. P. Martin, B. Jackson, A. Rambaut, O. Laguda-Akingba, S. Gaseitsiwe, A. von Gottberg, T. de Oliveira, Emergence of SARS-CoV-2 Omicron lineages BA.4 and BA.5 in South Africa. Nat. Med. 2022), doi:10.1038/s41591-022-01911-2.

13. N. P. Hachmann, J. Miller, A.-R. Y. Collier, J. D. Ventura, J. Yu, M. Rowe, E. A. Bondzie, O. Powers, N. Surve, K. Hall, D. H. Barouch, Neutralization Escape by SARS-CoV-2 Omicron Subvariants BA.2.12.1, BA.4, and BA.5. N. Engl. J. Med. 387, 86–88 (2022).

14. L. G. Thorne, M. Bouhaddou, A.-K. Reuschl, L. Zuliani-Alvarez, B. Polacco, A. Pelin, J. Batra, M. V. X. Whelan, M. Hosmillo, A. Fossati, R. Ragazzini, I. Jungreis, M. Ummadi, A. Rojc, J. Turner, M. L. Bischof, K. Obernier, H. Braberg, M. Soucheray, A. Richards, K.-H. Chen, B. Harjai, D. Memon, J. Hiatt, R. Rosales, B. L. McGovern, A. Jahun, J. M. Fabius, K. White, I. G. Goodfellow, Y. Takeuchi, P. Bonfanti, K. Shokat, N. Jura, K. Verba, M. Noursadeghi, P. Beltrao, M. Kellis, D. L. Swaney, A. García-Sastre, C. Jolly, G. J. Towers, N. J. Krogan, Evolution of enhanced innate immune evasion by SARS-CoV-2. Nature. 602, 487–495 (2022).

15. L. G. Thorne, A.-K. Reuschl, L. Zuliani-Alvarez, M. V. X. Whelan, J. Turner, M. Noursadeghi, C. Jolly, G. J. Towers, SARS-CoV-2 sensing by RIG-I and MDA5 links epithelial infection to macrophage inflammation. EMBO J. 40, e107826 (2021).

16. A. K. Banerjee, M. R. Blanco, E. A. Bruce, D. D. Honson, L. M. Chen, A. Chow, P. Bhat, N. Ollikainen, S. A. Quinodoz, C. Loney, J. Thai, Z. D. Miller, A. E. Lin, M. M. Schmidt, D. G. Stewart, D. Goldfarb, G. De Lorenzo, S. J. Rihn, R. M. Voorhees, J. W. Botten, D. Majumdar, M. Guttman, SARS-CoV-2 Disrupts Splicing, Translation, and Protein Trafficking to Suppress Host Defenses. Cell. 183, 1325–1339.e21 (2020).

17. M. Thoms, R. Buschauer, M. Ameismeier, L. Koepke, T. Denk, M. Hirschenberger, H. Kratzat, M. Hayn, T. Mackens-Kiani, J. Cheng, J. H. Straub, C. M. Stürzel, T. Fröhlich, O. Berninghausen, T. Becker, F. Kirchhoff, K. M. J. Sparrer, R. Beckmann, Structural basis for translational shutdown and immune evasion by the Nsp1 protein of SARS-CoV-2. Science. 369, 1249–1255 (2020).

18. M. Bouhaddou, D. Memon, B. Meyer, K. M. White R. V. v., M. Correa Marrero, B. J. Polacco, J. E. Melnyk, S. Ulferts, R. M. Kaake, J. Batra, A. L. Richards, E. Stevenson, D. E. Gordon, A. Rojc, K. Obernier, J. M. Fabius, M. Soucheray, L. Miorin, E. Moreno, C. Koh, Q. D. Tran, A. Hardy, R. Robinot, T. Vallet, B. E. Nilsson-Payant, C. Hernandez-Armenta, A. Dunham, S. Weigang, J. Knerr, M. Modak, D. Quintero, Y. Zhou, A. Dugourd, A. Valdeolivas, T. Patil, Q. Li, R. Hüttenhain, M. Cakir, M. Muralidharan, M. Kim, G. Jang, B. Tutuncuoglu, J. Hiatt, J. Z. Guo, J. Xu, S. Bouhaddou, C. J. P. Mathy, A. Gaulton, E. J. Manners, E. Félix, Y. Shi, M. Goff, J. K. Lim, T. McBride, M. C. O’Neal, Y. Cai, J. C. J. Chang, D. J. Broadhurst, S. Klippsten, E, A. R. Leach, T. Kortemme, B. Shoichet, M. Ott, J. Saez-Rodriguez, Br, R. D. Mullins, E. R. Fischer, G. Kochs, R. Grosse, A. García-Sastre, M. Vignuzzi, S. Johnson Jr, S. Km, B. Dl, K. p, N.j., The Global Phosphorylation Landscape of SARS-CoV-2 Infection. Cell. 182 (2020).

19. J. Cui, F. Li, Z.-L. Shi, Origin and evolution of pathogenic coronaviruses. Nat. Rev. Microbiol. 17, 181–192 (2018).

20. E. Simon-Loriere, E. C. Holmes, Why do RNA viruses recombine? Nat. Rev. Microbiol. 9, 617–626 (2011).

21. M. D. Parker, B. B. Lindsey, D. R. Shah, S. Hsu, A. J. Keeley, D. G. Partridge, S. Leary, A. Cope, Amy State, K. Johnson, N. Ali, R. Raghei, J. Heffer, N. Smith, P. Zhang, M. Gallis, S. F. Louka, H. R. Hornsby, M. Whiteley, B. H. Foulkes, S. Christou, P. Wolverson, M. Pohare, S. E. Hansford, L. R. Green, C. Evans, M. Raza, D. Wang, S. Gaudieri, S. Mallal, The COVID-19 Genomics UK (COG-UK) consortium, T. I. de Silva, Altered Subgenomic RNA Expression in SARS-CoV-2 B.1.1.7 Infections. bioRxiv (2021), p. 2021.03.02.433156.

22. A. M. Syed, T. Y. Taha, T. Tabata, I. P. Chen, A. Ciling, M. M. Khalid, B. Sreekumar, P.-Y. Chen, J. M. Hayashi, K. M. Soczek, M. Ott, J. A. Doudna, Rapid assessment of SARS-CoV-2-evolved variants using virus-like particles. Science. 374, 1626–1632 (2021).

23. A. M. Syed, A. Ciling, T. Y. Taha, I. P. Chen, M. M. Khalid, B. Sreekumar, P.-Y. Chen, G. R. Kumar, R. Suryawanshi, I. Silva, B. Milbes, N. Kojima, V. Hess, M. Shacreaw, L. Lopez, M. Brobeck, F. Turner, L. Spraggon, T. Tabata, M. Ott, J. A. Doudna, Omicron mutations enhance infectivity and reduce antibody neutralization of SARS-CoV-2 virus-like particles. Proc. Natl. Acad. Sci. U. S. A. 119, e2200592119 (2022).

24. S. Lu, Q. Ye, D. Singh, Y. Cao, J. K. Diedrich, J. R. Yates 3rd, E. Villa, D. W. Cleveland, K. D. Corbett, The SARS-CoV-2 nucleocapsid phosphoprotein forms mutually exclusive condensates with RNA and the membrane-associated M protein. Nat. Commun. 12, 502 (2021).

25. C. Wang, H. Xu, S. Lin, W. Deng, J. Zhou, Y. Zhang, Y. Shi, D. Peng, Y. Xue, GPS 5.0: An Update on the Prediction of Kinase-specific Phosphorylation Sites in Proteins. Genomics, Proteomics & Bioinformatics. 18 (2020), pp. 72–80.

26. C. R. Carlson, J. B. Asfaha, C. M. Ghent, C. J. Howard, N. Hartooni, M. Safari, A. D. Frankel, D. O. Morgan, Phosphoregulation of Phase Separation by the SARS-CoV-2 N Protein Suggests a Biophysical Basis for its Dual Functions. Mol. Cell. 80, 1092–1103.e4 (2020).

27. B. A. Johnson, Y. Zhou, K. G. Lokugamage, M. N. Vu, N. Bopp, P. A. Crocquet-Valdes, B. Kalveram, C. Schindewolf, Y. Liu, D. Scharton, J. A. Plante, X. Xie, P. Aguilar, S. C. Weaver, P.-Y. Shi, D. H. Walker, A. L. Routh, K. S. Plante, V. D. Menachery, Nucleocapsid mutations in SARS-CoV-2 augment replication and pathogenesis. PLoS Pathog. 18, e1010627 (2022).

28. M. Giurgiu, J. Reinhard, B. Brauner, I. Dunger-Kaltenbach, G. Fobo, G. Frishman, C. Montrone, A. Ruepp, CORUM: the comprehensive resource of mammalian protein complexes-2019. Nucleic Acids Res. 47, D559–D563 (2019).

29. D. E. Gordon, G. M. Jang, M. Bouhaddou, J. Xu, K. Obernier, K. M. White, M. J. O’Meara R. V. v., J. Z. Guo, D. L. Swaney, T. A. Tummino, R. Hüttenhain, R. M. Kaake, A. L. Richards, B. Tutuncuoglu, H. Foussard, J. Batra, K. Haas, M. Modak, M. Kim, P. Haas, B. J. Polacco, H. Braberg, J. M. Fabius, M. Eckhardt, M. Soucheray, M. J. Bennett, M. Cakir, M. J. McGregor, Q. Li, B. Meyer, F. Roesch, T. Vallet, A. Kain, L. Miorin, E. Moreno, Z. Z. C. Naing, Y. Zhou, S. Peng, Y. Shi, Z. Zhang, W. Shen, I. T. Kirby, J. E. Melnyk, J. S. Chorba, K. Lou, S. A. Dai, I. Barrio-Hernandez, D. Memon, C. Hernandez-Armenta, J. Lyu, C. J. P. Mathy, T. Perica, K. B. Pilla, S. J. Ganesan, D. J. Saltzberg, R. Rakesh, X. Liu, S. B. Rosenthal, L. Calviello, S. Venkataramanan, J. Liboy-Lugo, Y. Lin, X. P. Huang, Y. F. Liu, S. A. Wankowicz, M. Bohn, M. Safari, F. S. Ugur, C. Koh, N. S. Savar, Q. D. Tran, D. Shengjuler, S. J. Fletcher, M. C. O’Neal, Y. Cai, J. C. J. Chang, D. J. Broadhurst, S. Klippsten, P. P. Sharp, N. A. Wenzell, D. Kuzuoglu-Ozturk, H. Y. Wang, R. Trenker, J. M. Young, D. A. Cavero, J. Hiatt, T. L. Roth, U. Rathore, A. Subramanian, J. Noack, M. Hubert, R. M. Stroud, A. D. Frankel, O. S. Rosenberg, K. A. Verba, D. A. Agard, M. Ott, M. Emerman, N. Jura, M. Zastrow, E. Verdin, A. Ashworth, O. Schwartz, C. d’Enfert, S. Mukherjee, M. Jacobson, H. S. Malik, D. G. Fujimori, T. Ideker, C. S. Craik, S. N. Floor, J. S. Fraser, J. D. Gross, A. Sali, B. L. Roth, D. Ruggero, J. Taunton, T. Kortemme, P. Beltrao, M. Vignuzzi, A. García-Sastre, K. M. Shokat, B. K. Shoichet, N. J. Krogan, A SARS-CoV-2 protein interaction map reveals targets for drug repurposing. Nature. 583 (2020).

30. D. E. Gordon, J. Hiatt, M. Bouhaddou, V. V. Rezelj, S. Ulferts, H. Braberg, A. S. Jureka, K. Obernier, J. Z. Guo, J. Batra, R. M. Kaake, A. R. Weckstein, T. W. Owens, M. Gupta, S. Pourmal, E. W. Titus, M. Cakir, M. Soucheray, M. McGregor, Z. Cakir, G. Jang, M. J. O’Meara, T. A. Tummino, Z. Zhang, H. Foussard, A. Rojc, Y. Zhou, D. Kuchenov, R. Hüttenhain, J. Xu, M. Eckhardt, D. L. Swaney, J. M. Fabius, M. Ummadi, B. Tutuncuoglu, U. Rathore, M. Modak, P. Haas, K. M. Haas, Z. Z. C. Naing, E. H. Pulido, Y. Shi, I. Barrio-Hernandez, D. Memon, E. Petsalaki, A. Dunham, M. C. Marrero, D. Burke, C. Koh, T. Vallet, J. A. Silvas, C. M. Azumaya, C. Billesbølle, A. F. Brilot, M. G. Campbell, A. Diallo, M. S. Dickinson, D. Diwanji, N. Herrera, N. Hoppe, H. T. Kratochvil, Y. Liu, G. E. Merz, M. Moritz, H. C. Nguyen, C. Nowotny, C. Puchades, A. N. Rizo, U. Schulze-Gahmen, A. M. Smith, M. Sun, I. D. Young, J. Zhao, D. Asarnow, J. Biel, A. Bowen, J. R. Braxton, J. Chen, C. M. Chio, U. S. Chio, I. Deshpande, L. Doan, B. Faust, S. Flores, M. Jin, K. Kim, V. L. Lam, F. Li, J. Li, Y.-L. Li, Y. Li, X. Liu, M. Lo, K. E. Lopez, A. A. Melo, F. R. Moss 3rd, P. Nguyen, J. Paulino, K. I. Pawar, J. K. Peters, T. H. Pospiech Jr, M. Safari, S. Sangwan, K. Schaefer, P. V. Thomas, A. C. Thwin, R. Trenker, E. Tse, T. K. M. Tsui, F. Wang, N. Whitis, Z. Yu, K. Zhang, Y. Zhang, F. Zhou, D. Saltzberg, QCRG Structural Biology Consortium, A. J. Hodder, A. S. Shun-Shion, D. M. Williams, K. M. White, R. Rosales, T. Kehrer, L. Miorin, E. Moreno, A. H. Patel, S. Rihn, M. M. Khalid, A. Vallejo-Gracia, P. Fozouni, C. R. Simoneau, T. L. Roth, D. Wu, M. A. Karim, M. Ghoussaini, I. Dunham, F. Berardi, S. Weigang, M. Chazal, J. Park, J. Logue, M. McGrath, S. Weston, R. Haupt, C. J. Hastie, M. Elliott, F. Brown, K. A. Burness, E. Reid, M. Dorward, C. Johnson, S. G. Wilkinson, A. Geyer, D. M. Giesel, C. Baillie, S. Raggett, H. Leech, R. Toth, N. Goodman, K. C. Keough, A. L. Lind, Zoonomia Consortium, R. J. Klesh, K. R. Hemphill, J. Carlson-Stevermer, J. Oki, K. Holden, T. Maures, K. S. Pollard, A. Sali, D. A. Agard, Y. Cheng, J. S. Fraser, A. Frost, N. Jura, T. Kortemme, A. Manglik, D. R. Southworth, R. M. Stroud, D. R. Alessi, P. Davies, M. B. Frieman, T. Ideker, C. Abate, N. Jouvenet, G. Kochs, B. Shoichet, M. Ott, M. Palmarini, K. M. Shokat, A. García-Sastre, J. A. Rassen, R. Grosse, O. S. Rosenberg, K. A. Verba, C. F. Basler, M. Vignuzzi, A. A. Peden, P. Beltrao, N. J. Krogan, Comparative host-coronavirus protein interaction networks reveal pan-viral disease mechanisms. Science. 370 (2020), doi:10.1126/science.abe9403.

31. G. Teo, G. Liu, J. Zhang, A. I. Nesvizhskii, A.-C. Gingras, H. Choi, SAINTexpress: improvements and additional features in Significance Analysis of INTeractome software. J. Proteomics. 100, 37–43 (2014).

32. S. Jäger, P. Cimermancic, N. Gulbahce, J. R. Johnson, K. E. McGovern, S. C. Clarke, M. Shales, G. Mercenne, L. Pache, K. Li, H. Hernandez, G. M. Jang, S. L. Roth, E. Akiva, J. Marlett, M. Stephens, I. D’Orso, J. Fernandes, M. Fahey, C. Mahon, A. J. O’Donoghue, A. Todorovic, J. H. Morris, D. A. Maltby, T. Alber, G. Cagney, F. D. Bushman, J. A. Young, S. K. Chanda, W. I. Sundquist, T. Kortemme, R. D. Hernandez, C. S. Craik, A. Burlingame, A. Sali, A. D. Frankel, N. J. Krogan, Global landscape of HIV-human protein complexes. Nature. 481, 365–370 (2011).

33. M. Choi, C.-Y. Chang, T. Clough, D. Broudy, T. Killeen, B. MacLean, O. Vitek, MSstats: an R package for statistical analysis of quantitative mass spectrometry-based proteomic experiments. Bioinformatics. 30 (2014), pp. 2524–2526.

34. N. J. Krogan, M. Kim, S. H. Ahn, G. Zhong, M. S. Kobor, G. Cagney, A. Emili, A. Shilatifard, S. Buratowski, J. F. Greenblatt, RNA polymerase II elongation factors of Saccharomyces cerevisiae: a targeted proteomics approach. Mol. Cell. Biol. 22, 6979–6992 (2002).

35. B. Zhu, S. S. Mandal, A.-D. Pham, Y. Zheng, H. Erdjument-Bromage, S. K. Batra, P. Tempst, D. Reinberg, The human PAF complex coordinates transcription with events downstream of RNA synthesis. Genes Dev. 19, 1668–1673 (2005).

36. S. B. Van Oss, C. E. Cucinotta, K. M. Arndt, Emerging Insights into the Roles of the Paf1 Complex in Gene Regulation. Trends Biochem. Sci. 42, 788–798 (2017).

37. I. Marazzi, J. S. Y. Ho, J. Kim, B. Manicassamy, S. Dewell, R. A. Albrecht, C. W. Seibert, U. Schaefer, K. L. Jeffrey, R. K. Prinjha, K. Lee, A. García-Sastre, R. G. Roeder, A. Tarakhovsky, Suppression of the antiviral response by an influenza histone mimic. Nature. 483, 428–433 (2012).

38. C. New, Z.-Y. Lee, K. S. Tan, A. H.-P. Wong, D. Y. Wang, T. Tran, Tetraspanins: Host Factors in Viral Infections. Int. J. Mol. Sci. 22 (2021), doi:10.3390/ijms222111609.

39. J. T. Earnest, M. P. Hantak, K. Li, P. B. McCray Jr, S. Perlman, T. Gallagher, The tetraspanin CD9 facilitates MERS-coronavirus entry by scaffolding host cell receptors and proteases. PLoS Pathog. 13, e1006546 (2017).

40. I. P. Chen, J. E. Longbotham, S. McMahon, R. K. Suryawanshi, M. M. Khalid, T. Y. Taha, T. Tabata, J. M. Hayashi, F. W. Soveg, J. Carlson-Stevermer, M. Gupta, M. Y. Zhang, V. L. Lam, Y. Li, Z. Yu, E. W. Titus, A. Diallo, J. Oki, K. Holden, N. Krogan, D. G. Fujimori, M. Ott, Viral E protein neutralizes BET protein-mediated post-entry antagonism of SARS-CoV-2. Cell Rep. 40, 111088 (2022).

41. D. Szklarczyk, A. L. Gable, K. C. Nastou, D. Lyon, R. Kirsch, S. Pyysalo, N. T. Doncheva, M. Legeay, T. Fang, P. Bork, L. J. Jensen, C. von Mering, The STRING database in 2021: customizable protein–protein networks, and functional characterization of user-uploaded gene/measurement sets. Nucleic Acids Research. 49 (2021), pp. D605–D612.

42. K. M. White, R. Rosales, S. Yildiz, T. Kehrer, L. Miorin, E. Moreno, S. Jangra, M. B. Uccellini, R. Rathnasinghe, L. Coughlan, C. Martinez-Romero, J. Batra, A. Rojc, M. Bouhaddou, J. M. Fabius, K. Obernier, M. Dejosez, M. J. Guillén, A. Losada, P. Avilés, M. Schotsaert, T. Zwaka, M. Vignuzzi, K. M. Shokat, N. J. Krogan, A. García-Sastre, Plitidepsin has potent preclinical efficacy against SARS-CoV-2 by targeting the host protein eEF1A. Science. 371 (2021).

43. Trial to Determine the Efficacy/Safety of Plitidepsin vs Control in Patients With Moderate COVID-19 Infection - Full Text View - ClinicalTrials.gov, (available at https://clinicaltrials.gov/ct2/show/NCT04784559).

44. J. F. Varona, P. Landete, J. A. Lopez-Martin, V. Estrada, R. Paredes, P. Guisado-Vasco, L. Fernandez de Orueta, M. Torralba, J. Fortun, R. Vates, J. Barberan, B. Clotet, J. Ancochea, D. Carnevali, N. Cabello, L. Porras, P. Gijon, A. Monereo, D. Abad, S. Zuñiga, I. Sola, J. Rodon, J. Vergara-Alert, N. Izquierdo-Useros, S. Fudio, M. J. Pontes, B. de Rivas, P. Giron de Velasco, A. Nieto, J. Gomez, P. Aviles, R. Lubomirov, A. Belgrano, B. Sopesen, K. M. White, R. Rosales, S. Yildiz, A.-K. Reuschl, L. G. Thorne, C. Jolly, G. J. Towers, L. Zuliani-Alvarez, M. Bouhaddou, K. Obernier, B. L. McGovern, M. L. Rodriguez, L. Enjuanes, J. M. Fernandez-Sousa, N. J. Krogan, J. M. Jimeno, A. Garcia-Sastre, Preclinical and randomized phase I studies of plitidepsin in adults hospitalized with COVID-19. Life Sci Alliance. 5 (2022), doi:10.26508/lsa.202101200.

45. J. L. Tenthorey, M. Emerman, H. S. Malik, Evolutionary Landscapes of Host-Virus Arms Races. Annu. Rev. Immunol. 40, 271–294 (2022).

46. A. G. Laing, A. Lorenc, I. Del Molino Del Barrio, A. Das, M. Fish, L. Monin, M. Muñoz-Ruiz, D. R. McKenzie, T. S. Hayday, I. Francos-Quijorna, S. Kamdar, M. Joseph, D. Davies, R. Davis, A. Jennings, I. Zlatareva, P. Vantourout, Y. Wu, V. Sofra, F. Cano, M. Greco, E. Theodoridis, J. D. Freedman, S. Gee, J. N. E. Chan, S. Ryan, E. Bugallo-Blanco, P. Peterson, K. Kisand, L. Haljasmägi, L. Chadli, P. Moingeon, L. Martinez, B. Merrick, K. Bisnauthsing, K. Brooks, M. A. A. Ibrahim, J. Mason, F. Lopez Gomez, K. Babalola, S. Abdul-Jawad, J. Cason, C. Mant, J. Seow, C. Graham, K. J. Doores, F. Di Rosa, J. Edgeworth, M. Shankar-Hari, A. C. Hayday, A dynamic COVID-19 immune signature includes associations with poor prognosis. Nat. Med. 26, 1623–1635 (2020).

47. J. Hadjadj, N. Yatim, L. Barnabei, A. Corneau, J. Boussier, N. Smith, H. Péré, B. Charbit, V. Bondet, C. Chenevier-Gobeaux, P. Breillat, N. Carlier, R. Gauzit, C. Morbieu, F. Pène, N. Marin, N. Roche, T.-A. Szwebel, S. H. Merkling, J.-M. Treluyer, D. Veyer, L. Mouthon, C. Blanc, P.-L. Tharaux, F. Rozenberg, A. Fischer, D. Duffy, F. Rieux-Laucat, S. Kernéis, B. Terrier, Impaired type I interferon activity and inflammatory responses in severe COVID-19 patients. Science. 369, 718–724 (2020).

48. G. Chen, D. Wu, W. Guo, Y. Cao, D. Huang, H. Wang, T. Wang, X. Zhang, H. Chen, H. Yu, X. Zhang, M. Zhang, S. Wu, J. Song, T. Chen, M. Han, S. Li, X. Luo, J. Zhao, Q. Ning, Clinical and immunological features of severe and moderate coronavirus disease 2019. J. Clin. Invest. 130, 2620–2629 (2020).

49. M. S. Abers, O. M. Delmonte, E. E. Ricotta, J. Fintzi, D. L. Fink, A. A. A. de Jesus, K. A. Zarember, S. Alehashemi, V. Oikonomou, J. V. Desai, S. W. Canna, B. Shakoory, K. Dobbs, L. Imberti, A. Sottini, E. Quiros-Roldan, F. Castelli, C. Rossi, D. Brugnoni, A. Biondi, L. R. Bettini, M. D’Angio’, P. Bonfanti, R. Castagnoli, D. Montagna, A. Licari, G. L. Marseglia, E. F. Gliniewicz, E. Shaw, D. E. Kahle, A. T. Rastegar, M. Stack, K. Myint-Hpu, S. L. Levinson, M. J. DiNubile, D. W. Chertow, P. D. Burbelo, J. I. Cohen, K. R. Calvo, J. S. Tsang, NIAID COVID-19 Consortium, H. C. Su, J. I. Gallin, D. B. Kuhns, R. Goldbach-Mansky, M. S. Lionakis, L. D. Notarangelo, An immune-based biomarker signature is associated with mortality in COVID-19 patients. JCI Insight. 6 (2021), doi:10.1172/jci.insight.144455.

50. D. M. Del Valle, S. Kim-Schulze, H.-H. Huang, N. D. Beckmann, S. Nirenberg, B. Wang, Y. Lavin, T. H. Swartz, D. Madduri, A. Stock, T. U. Marron, H. Xie, M. Patel, K. Tuballes, O. Van Oekelen, A. Rahman, P. Kovatch, J. A. Aberg, E. Schadt, S. Jagannath, M. Mazumdar, A. W. Charney, A. Firpo-Betancourt, D. R. Mendu, J. Jhang, D. Reich, K. Sigel, C. Cordon-Cardo, M. Feldmann, S. Parekh, M. Merad, S. Gnjatic, An inflammatory cytokine signature predicts COVID-19 severity and survival. Nat. Med. 26, 1636–1643 (2020).

51. C. Lucas, P. Wong, J. Klein, T. B. R. Castro, J. Silva, M. Sundaram, M. K. Ellingson, T. Mao, J. E. Oh, B. Israelow, T. Takahashi, M. Tokuyama, P. Lu, A. Venkataraman, A. Park, S. Mohanty, H. Wang, A. L. Wyllie, C. B. F. Vogels, R. Earnest, S. Lapidus, I. M. Ott, A. J. Moore, M. C. Muenker, J. B. Fournier, M. Campbell, C. D. Odio, A. Casanovas-Massana, Yale IMPACT Team, R. Herbst, A. C. Shaw, R. Medzhitov, W. L. Schulz, N. D. Grubaugh, C. Dela Cruz, S. Farhadian, A. I. Ko, S. B. Omer, A. Iwasaki, Longitudinal analyses reveal immunological misfiring in severe COVID-19. Nature. 584, 463–469 (2020).

52. A.-K. Reuschl, L. G. Thorne, M. V. X. Whelan, D. Mesner, R. Ragazzini, G. Dowgier, N. Bogoda, J. L. E. Turner, W. Furnon, V. M. Cowton, G. de Lorenzo, P. Bonfanti, M. Palmarini, A. H. Patel, C. Jolly, G. J. Towers, Enhanced innate immune suppression by SARS-CoV-2 Omicron subvariants BA.4 and BA.5. bioRxiv (2022), p. 2022.07.12.499603.

53. L. Miorin, T. Kehrer, M. T. Sanchez-Aparicio, K. Zhang, P. Cohen, R. S. Patel, A. Cupic, T. Makio, M. Mei, E. Moreno, O. Danziger, K. M. White, R. Rathnasinghe, M. Uccellini, S. Gao, T. Aydillo, I. Mena, X. Yin, L. Martin-Sancho, N. J. Krogan, S. K. Chanda, M. Schotsaert, R. W. Wozniak, Y. Ren, B. R. Rosenberg, B. M. A. Fontoura, A. García-Sastre, SARS-CoV-2 Orf6 hijacks Nup98 to block STAT nuclear import and antagonize interferon signaling. Proc. Natl. Acad. Sci. U. S. A. 117, 28344–28354 (2020).

54. S. Wang, T. Dai, Z. Qin, T. Pan, F. Chu, L. Lou, L. Zhang, B. Yang, H. Huang, H. Lu, F. Zhou, Targeting liquid-liquid phase separation of SARS-CoV-2 nucleocapsid protein promotes innate antiviral immunity by elevating MAVS activity. Nat. Cell Biol. 23, 718–732 (2021).

55. H. Liu, Y. Bai, X. Zhang, T. Gao, Y. Liu, E. Li, X. Wang, Z. Cao, L. Zhu, Q. Dong, Y. Hu, G. Wang, C. Song, X. Niu, T. Zheng, D. Wang, Z. Liu, Y. Jin, P. Li, X. Bian, C. Cao, X. Liu, SARS-CoV-2 N Protein Antagonizes Stress Granule Assembly and IFN Production by Interacting with G3BPs to Facilitate Viral Replication. J. Virol. 96, e0041222 (2022).

56. Nextstrain, (available at https://nextstrain.org/).

57. Nextstrain, (available at https://nextstrain.org/).

58. A. Addetia, N. A. P. Lieberman, Q. Phung, T.-Y. Hsiang, H. Xie, P. Roychoudhury, L. Shrestha, M. A. Loprieno, M.-L. Huang, M. Gale Jr, K. R. Jerome, A. L. Greninger, SARS-CoV-2 ORF6 Disrupts Bidirectional Nucleocytoplasmic Transport through Interactions with Rae1 and Nup98. MBio. 12 (2021), doi:10.1128/mBio.00065-21.

59. T. Li, Y. Wen, H. Guo, T. Yang, H. Yang, X. Ji, Molecular Mechanism of SARS-CoVs Orf6 Targeting the Rae1-Nup98 Complex to Compete With mRNA Nuclear Export. Front Mol Biosci. 8, 813248 (2021).

60. X. Huang, W. Zheng, R. Pearce, Y. Zhang, SSIPe: accurately estimating protein-protein binding affinity change upon mutations using evolutionary profiles in combination with an optimized physical energy function. Bioinformatics. 36, 2429–2437 (2020).

61. L. Z. Alvarez, M. L. Govasli, J. Rasaiyaah, C. Monit, S. O. Perry, R. P. Sumner, S. McAlpine-Scott, C. Dickson, K. M. Rifat Faysal, L. Hilditch, R. J. Miles, F. Bibollet-Ruche, B. H. Hahn, T. Boecking, N. Pinotsis, L. C. James, D. A. Jacques, G. J. Towers, Macrophage activation of cGAS and TRIM5 distinguish pandemic and non-pandemic HIV. bioRxiv (2022), p. 2022.01.21.477263.

62. K. P. Y. Hui, J. C. W. Ho, M.-C. Cheung, K.-C. Ng, R. H. H. Ching, K.-L. Lai, T. T. Kam, H. Gu, K.-Y. Sit, M. K. Y. Hsin, T. W. K. Au, L. L. M. Poon, M. Peiris, J. M. Nicholls, M. C. W. Chan, SARS-CoV-2 Omicron variant replication in human bronchus and lung ex vivo. Nature. 603, 715–720 (2022).

63. H. Shuai, J. F.-W. Chan, B. Hu, Y. Chai, T. T.-T. Yuen, F. Yin, X. Huang, C. Yoon, J.-C. Hu, H. Liu, J. Shi, Y. Liu, T. Zhu, J. Zhang, Y. Hou, Y. Wang, L. Lu, J.-P. Cai, A. J. Zhang, J. Zhou, S. Yuan, M. A. Brindley, B.-Z. Zhang, J.-D. Huang, K. K.-W. To, K.-Y. Yuen, H. Chu, Attenuated replication and pathogenicity of SARS-CoV-2 B.1.1.529 Omicron. Nature. 603, 693–699 (2022).

64. A. A. Butt, S. R. Dargham, H. Chemaitelly, A. Al Khal, P. Tang, M. R. Hasan, P. V. Coyle, A. G. Thomas, A. M. Borham, E. G. Concepcion, A. H. Kaleeckal, A. N. Latif, R. Bertollini, A.-B. Abou-Samra, L. J. Abu-Raddad, Severity of Illness in Persons Infected With the SARS-CoV-2 Delta Variant vs Beta Variant in Qatar. JAMA Intern. Med. 182, 197–205 (2022).

65. K. A. Twohig, T. Nyberg, A. Zaidi, S. Thelwall, M. A. Sinnathamby, S. Aliabadi, S. R. Seaman, R. J. Harris, R. Hope, J. Lopez-Bernal, E. Gallagher, A. Charlett, D. De Angelis, A. M. Presanis, G. Dabrera, COVID-19 Genomics UK (COG-UK) consortium, Hospital admission and emergency care attendance risk for SARS-CoV-2 delta (B.1.617.2) compared with alpha (B.1.1.7) variants of concern: a cohort study. Lancet Infect. Dis. 22, 35–42 (2022).

66. S. W. X. Ong, C. J. Chiew, L. W. Ang, T.-M. Mak, L. Cui, M. P. H. S. Toh, Y. D. Lim, P. H. Lee, T. H. Lee, P. Y. Chia, S. Maurer-Stroh, R. T. P. Lin, Y.-S. Leo, V. J. Lee, D. C. Lye, B. E. Young, Clinical and Virological Features of SARS-CoV-2 Variants of Concern: A Retrospective Cohort Study Comparing B.1.1.7 (Alpha), B.1.315 (Beta), and B.1.617.2 (Delta). SSRN Electronic Journal,, doi:10.2139/ssrn.3861566.

67. A. Sigal, R. Milo, W. Jassat, Estimating disease severity of Omicron and Delta SARS-CoV-2 infections. Nat. Rev. Immunol. 22, 267–269 (2022).

68. F. Abdullah, J. Myers, D. Basu, G. Tintinger, V. Ueckermann, M. Mathebula, R. Ramlall, S. Spoor, T. de Villiers, Z. Van der Walt, J. Cloete, P. Soma-Pillay, P. Rheeder, F. Paruk, A. Engelbrecht, V. Lalloo, M. Myburg, J. Kistan, W. van Hougenhouck-Tulleken, M. T. Boswell, G. Gray, R. Welch, L. Blumberg, W. Jassat, Decreased severity of disease during the first global omicron variant covid-19 outbreak in a large hospital in tshwane, south africa. International Journal of Infectious Diseases. 116 (2022), pp. 38–42.

69. L. Veneti, H. Bøås, A. Bråthen Kristoffersen, J. Stålcrantz, K. Bragstad, O. Hungnes, M. L. Storm, N. Aasand, G. Rø, J. Starrfelt, E. Seppälä, R. Kvåle, L. Vold, K. Nygård, E. A. Buanes, R. Whittaker, Reduced risk of hospitalisation among reported COVID-19 cases infected with the SARS-CoV-2 Omicron BA.1 variant compared with the Delta variant, Norway, December 2021 to January 2022. Euro Surveill. 27 (2022), doi:10.2807/1560-7917.ES.2022.27.4.2200077.

70. T. Nyberg, N. M. Ferguson, S. G. Nash, H. H. Webster, S. Flaxman, N. Andrews, W. Hinsley, J. L. Bernal, M. Kall, S. Bhatt, P. B. Blomquist, A. Zaidi, E. Volz, N. A. Aziz, K. Harman, R. Hope, A. Charlett, M. A. Chand, A. Ghani, S. Seaman, G. Dabrera, D. DeAngelis, A. M. Presanis, S. Thelwall, Comparative Analysis of the Risks of Hospitalisation and Death Associated with SARS-CoV-2 Omicron (B.1.1.529) and Delta (B.1.617.2) Variants in England. SSRN Electronic Journal,, doi:10.2139/ssrn.4025932.

71. C. Sievers, B. Zacher, A. Ullrich, M. Huska, S. Fuchs, S. Buda, W. Haas, M. Diercke, M. An der Heiden, S. Kröger, SARS-CoV-2 Omicron variants BA.1 and BA.2 both show similarly reduced disease severity of COVID-19 compared to Delta, Germany, 2021 to 2022. Euro Surveill. 27 (2022), doi:10.2807/1560-7917.ES.2022.27.22.2200396.

72. I. Kislaya, P. Casaca, V. Borges, C. Sousa, B. I. Ferreira, E. Fernandes, C. M. Dias, S. Duarte, J. P. Almeida, I. Grenho, L. Coelho, R. Ferreira, P. P. Ferreira, J. Isidro, M. Pinto, L. Menezes, D. Sobral, A. Nunes, D. Santos, A. M. Gonçalves, L. Vieira, J. P. Gomes, P. P. Leite, B. Nunes, A. Machado, A. Peralta-Santos, SARS-CoV-2 BA.5 vaccine breakthrough risk and severity compared with BA.2: a case-case and cohort study using Electronic Health Records in Portugal. medRxiv (2022), p. 2022.07.25.22277996.

73. A. Waterhouse, M. Bertoni, S. Bienert, G. Studer, G. Tauriello, R. Gumienny, F. T. Heer, T. A. P. de Beer, C. Rempfer, L. Bordoli, R. Lepore, T. Schwede, SWISS-MODEL: homology modelling of protein structures and complexes. Nucleic Acids Res. 46, W296–W303 (2018).

74. Y. Perez-Riverol, J. Bai, C. Bandla, D. García-Seisdedos, S. Hewapathirana, S. Kamatchinathan, D. J. Kundu, A. Prakash, A. Frericks-Zipper, M. Eisenacher, M. Walzer, S. Wang, A. Brazma, J. A. Vizcaíno, The PRIDE database resources in 2022: a hub for mass spectrometry-based proteomics evidences. Nucleic Acids Res. 50, D543–D552 (2022).

75. T. Barrett, S. E. Wilhite, P. Ledoux, C. Evangelista, I. F. Kim, M. Tomashevsky, K. A. Marshall, K. H. Phillippy, P. M. Sherman, M. Holko, A. Yefanov, H. Lee, N. Zhang, C. L. Robertson, N. Serova, S. Davis, A. Soboleva, NCBI GEO: archive for functional genomics data sets--update. Nucleic Acids Res. 41, D991–5 (2013).

76. J. Zhou, T. P. Peacock, J. C. Brown, D. H. Goldhill, A. M. E. Elrefaey, R. Penrice-Randal, V. M. Cowton, G. De Lorenzo, W. Furnon, W. T. Harvey, R. Kugathasan, R. Frise, L. Baillon, R. Lassaunière, N. Thakur, G. Gallo, H. Goldswain, I. ‘ah Donovan-Banfield, X. Dong, N. P. Randle, F. Sweeney, M. C. Glynn, J. L. Quantrill, P. F. McKay, A. H. Patel, M. Palmarini, J. A. Hiscox, D. Bailey, W. S. Barclay, Mutations that adapt SARS-CoV-2 to mink or ferret do not increase fitness in the human airway. Cell Reports. 38 (2022), p. 110344.

77. T. Thi Nhu Thao, F. Labroussaa, N. Ebert, P. V’kovski, H. Stalder, J. Portmann, J. Kelly, S. Steiner, M. Holwerda, A. Kratzel, M. Gultom, K. Schmied, L. Laloli, L. Hüsser, M. Wider, S. Pfaender, D. Hirt, V. Cippà, S. Crespo-Pomar, S. Schröder, D. Muth, D. Niemeyer, V. M. Corman, M. A. Müller, C. Drosten, R. Dijkman, J. Jores, V. Thiel, Rapid reconstruction of SARS-CoV-2 using a synthetic genomics platform. Nature. 582, 561–565 (2020).

78. C. Pattabiraman, F. Habib, P. K. Harsha, R. Rasheed, V. Reddy, P. Dinesh, T. Damodar, K. Hosallimath, A. K. George, N. V. K. Reddy, B. John, A. Pattanaik, N. Kumar, R. S. Mani, M. M. Venkataswamy, S. K. Shahul Hameed, B. G. Prakash Kumar, A. Desai, R. Vasanthapuram, Genomic epidemiology reveals multiple introductions and spread of SARS-CoV-2 in the Indian state of Karnataka,, doi:10.1101/2020.07.10.20150045.

79. J. Schindelin, I. Arganda-Carreras, E. Frise, V. Kaynig, M. Longair, T. Pietzsch, S. Preibisch, C. Rueden, S. Saalfeld, B. Schmid, J.-Y. Tinevez, D. J. White, V. Hartenstein, K. Eliceiri, P. Tomancak, A. Cardona, Fiji: an open-source platform for biological-image analysis. Nat. Methods. 9, 676–682 (2012).

80. L. Miorin, C. E. Mire, S. Ranjbar, A. J. Hume, J. Huang, N. A. Crossland, K. M. White, M. Laporte, T. Kehrer, V. Haridas, E. Moreno, A. Nambu, S. Jangra, A. Cupic, M. Dejosez, K. A. Abo, A. E. Tseng, R. B. Werder, R. Rathnasinghe, T. Mutetwa, I. Ramos, J. S. de Aja, C. G. de Alba Rivas, M. Schotsaert, R. B. Corley, J. V. Falvo, A. Fernandez-Sesma, C. Kim, J.-F. Rossignol, A. A. Wilson, T. Zwaka, D. N. Kotton, E. Mühlberger, A. García-Sastre, A. E. Goldfeld, The oral drug nitazoxanide restricts SARS-CoV-2 infection and attenuates disease pathogenesis in Syrian hamsters. bioRxiv (2022), p. 2022.02.08.479634.

81. C. Ye, K. Chiem, J.-G. Park, F. Oladunni, R. N. Platt 2nd, T. Anderson, F. Almazan, J. C. de la Torre, L. Martinez-Sobrido, Rescue of SARS-CoV-2 from a Single Bacterial Artificial Chromosome. MBio. 11 (2020), doi:10.1128/mBio.02168-20.

82. F. Amanat, K. M. White, L. Miorin, S. Strohmeier, M. McMahon, P. Meade, W. Liu, R. A. Albrecht, V. Simon, L. Martinez-Sobrido, T. Moran, A. García-Sastre, F. Krammer, An In Vitro Microneutralization Assay for SARS-CoV-2 Serology and Drug Screening. Current Protocols in Microbiology. 58 (2020),, doi:10.1002/cpmc.108.

83. D. Kim, J. M. Paggi, C. Park, C. Bennett, S. L. Salzberg, Graph-based genome alignment and genotyping with HISAT2 and HISAT-genotype. Nat. Biotechnol. 37, 907–915 (2019).

84. S. Kovaka, A. V. Zimin, G. M. Pertea, R. Razaghi, S. L. Salzberg, M. Pertea, Transcriptome assembly from long-read RNA-seq alignments with StringTie2. Genome Biol. 20, 278 (2019).

85. M. I. Love, W. Huber, S. Anders, Moderated estimation of fold change and dispersion for RNA-seq data with DESeq2. Genome Biol. 15, 550 (2014).

86. A. Bankevich, S. Nurk, D. Antipov, A. A. Gurevich, M. Dvorkin, A. S. Kulikov, V. M. Lesin, S. I. Nikolenko, S. Pham, A. D. Prjibelski, A. V. Pyshkin, A. V. Sirotkin, N. Vyahhi, G. Tesler, M. A. Alekseyev, P. A. Pevzner, SPAdes: a new genome assembly algorithm and its applications to single-cell sequencing. J. Comput. Biol. 19, 455–477 (2012).

87. UniProt Consortium, UniProt: the universal protein knowledgebase in 2021. Nucleic Acids Res. 49, D480–D489 (2021).

88. M. Steinegger, J. Söding, MMseqs2 enables sensitive protein sequence searching for the analysis of massive data sets. Nat. Biotechnol. 35, 1026–1028 (2017).

89. S. Bienert, A. Waterhouse, T. A. P. de Beer, G. Tauriello, G. Studer, L. Bordoli, T. Schwede, The SWISS-MODEL Repository-new features and functionality. Nucleic Acids Res. 45, D313–D319 (2017).

90. P. Benkert, M. Biasini, T. Schwede, Toward the estimation of the absolute quality of individual protein structure models. Bioinformatics. 27, 343–350 (2011).

91. J. Köster, S. Rahmann, Snakemake--a scalable bioinformatics workflow engine. Bioinformatics. 28, 2520–2522 (2012).

92. A. Dunham, G. M. Jang, M. Muralidharan, D. Swaney, P. Beltrao, A missense variant effect prediction and annotation resource for SARS-CoV-2. bioRxiv (2021), p. 2021.02.24.432721.

93. J. Cox, M. Mann, MaxQuant enables high peptide identification rates, individualized p.p.b.-range mass accuracies and proteome-wide protein quantification. Nature Biotechnology. 26 (2008), pp. 1367–1372.

94. J. Cox, M. Y. Hein, C. A. Luber, I. Paron, N. Nagaraj, M. Mann, Accurate proteome-wide label-free quantification by delayed normalization and maximal peptide ratio extraction, termed MaxLFQ. Mol. Cell. Proteomics. 13, 2513–2526 (2014).

95. E. Verschueren, J. Von Dollen, P. Cimermancic, N. Gulbahce, A. Sali, N. J. Krogan, Scoring Large-Scale Affinity Purification Mass Spectrometry Datasets with MiST. Curr. Protoc. Bioinformatics. 49, 8.19.1–8.19.16 (2015).

96. P. Shannon, A. Markiel, O. Ozier, N. S. Baliga, J. T. Wang, D. Ramage, N. Amin, B. Schwikowski, T. Ideker, Cytoscape: a software environment for integrated models of biomolecular interaction networks. Genome Res. 13, 2498–2504 (2003).

97. T. Wu, E. Hu, S. Xu, M. Chen, P. Guo, Z. Dai, T. Feng, L. Zhou, W. Tang, L. Zhan, X. Fu, S. Liu, X. Bo, G. Yu, clusterProfiler 4.0: A universal enrichment tool for interpreting omics data. The Innovation. 2 (2021), p. 100141.

98. D. Ochoa, A. F. Jarnuczak, C. Viéitez, M. Gehre, M. Soucheray, A. Mateus, A. A. Kleefeldt, A. Hill, L. Garcia-Alonso, F. Stein, N. J. Krogan, M. M. Savitski, D. L. Swaney, J. A. Vizcaíno, K.-M. Noh, P. Beltrao, The functional landscape of the human phosphoproteome. Nat. Biotechnol. 38, 365–373 (2019).

99. M. Cao, H. Zhang, J. Park, N. M. Daniels, M. E. Crovella, L. J. Cowen, B. Hescott, Going the Distance for Protein Function Prediction: A New Distance Metric for Protein Interaction Networks. PLoS One. 8, e76339 (2013).

100. S. Choobdar, M. E. Ahsen, J. Crawford, M. Tomasoni, T. Fang, D. Lamparter, J. Lin, B. Hescott, X. Hu, J. Mercer, T. Natoli, R. Narayan, A. Subramanian, J. D. Zhang, G. Stolovitzky, Z. Kutalik, K. Lage, D. K. Slonim, J. Saez-Rodriguez, L. J. Cowen, S. Bergmann, D. Marbach, Assessment of network module identification across complex diseases. Nat. Methods. 16, 843–852 (2019).

101. D. Szklarczyk, A. L. Gable, D. Lyon, A. Junge, S. Wyder, J. Huerta-Cepas, M. Simonovic, N. T. Doncheva, J. H. Morris, P. Bork, L. J. Jensen, C. von Mering, STRING v11: protein-protein association networks with increased coverage, supporting functional discovery in genome-wide experimental datasets. Nucleic Acids Res. 47, D607–D613 (2019).

102. L. Cowen, T. Ideker, B. J. Raphael, R. Sharan, Network propagation: a universal amplifier of genetic associations. Nat. Rev. Genet. 18, 551–562 (2017).

